# Alignment of spatial transcriptomics data using diffeomorphic metric mapping

**DOI:** 10.1101/2023.04.11.534630

**Authors:** Kalen Clifton, Manjari Anant, Gohta Aihara, Lyla Atta, Osagie K. Aimiuwu, Justus M. Kebschull, Michael I. Miller, Daniel Tward, Jean Fan

## Abstract

Spatial transcriptomics (ST) technologies enable high throughput gene expression characterization within thin tissue sections. However, comparing spatial observations across sections, samples, and technologies remains challenging. To address this challenge, we developed STalign to align ST datasets in a manner that accounts for partially matched tissue sections and other local non-linear distortions using diffeomorphic metric mapping. We apply STalign to align ST datasets within and across technologies as well as to align ST datasets to a 3D common coordinate framework. We show that STalign achieves high gene expression and cell-type correspondence across matched spatial locations that is significantly improved over landmark-based affine alignments. Applying STalign to align ST datasets of the mouse brain to the 3D common coordinate framework from the Allen Brain Atlas, we highlight how STalign can be used to lift over brain region annotations and enable the interrogation of compositional heterogeneity across anatomical structures. STalign is available as an open-source Python toolkit at https://github.com/JEFworks-Lab/STalign and as supplementary software with additional documentation and tutorials available at https://jef.works/STalign.

## Introduction

Spatial transcriptomics (ST) technologies have enabled high-throughput, quantitative profiling of gene expression within individual cells and small groups of cells in fixed, thin tissue sections. Comparative analysis of ST datasets at matched spatial locations across tissues, individuals, and samples provides the opportunity to interrogate spatial gene expression and cell-type compositional variation in the context of health and disease. Such comparative analysis is complicated by technical challenges such as in sample collection, where the experimental process may induce tissue rotations, tears, and other structural distortions. Other challenges include biological variation such as natural inter-individual tissue structural differences. In order to reliably characterize spatial molecular differences between ST datasets along comparative axes of interest, it is integral to control for potentially confounding tissue structural variation by spatially aligning these tissue structures across ST datasets.

Considering the recent development of such ST technologies, options for spatially aligning across ST datasets are still limited. Previous computational methods have focused on spatial alignment of ST datasets for which each dataset is assayed using the same pixel-resolution ST technology with only a few hundred to a few thousand spatial measurements^1,2^. These methods face challenges in scaling to larger, single-cell resolution ST datasets with tens to hundreds of thousands of spatial measurements. Further, spatial alignment of datasets across different ST technologies remains challenging. Other alignment methods are limited to rigid, affine transformation such as based on landmarks^3^ and cannot accommodate non-linear distortions. To address these challenges, we present an approach called STalign that builds on recent developments in Large Deformation Diffeomorphic Metric Mapping^4,5^ (LDDMM) to align ST datasets using image varifolds. STalign is amenable to data from single-cell resolution ST technologies as well as data from multi-cellular pixel-resolution ST technologies for which a corresponding registered single-cell resolution image such as a histology image is available. STalign is further able to accommodate alignment in both 2D and 3D coordinate systems. STalign is available as an open-source Python toolkit at https://github.com/JEFworks-Lab/STalign and as supplementary software with additional documentation and tutorials available at https://jef.works/STalign.

## Results

### Overview of Method

To align two ST datasets, STalign solves a mapping that minimizes the dissimilarity between a source and a target ST dataset subject to regularization penalties (Online Methods). Within single-cell resolution ST technologies, both the source and target ST datasets are represented as cellular positions (*x*^*ρ,s*^, *y*^*ρ,s*^) and (*x*^*ρ,T*^,*y*^*ρ,T*^) respectively (Fig 1a). Solving the mapping with respect to single cells has quadratic complexity and is computationally intractable, so STalign applies a rasterization approach to reduce computational time (Fig 1b). Briefly, STalign models the positions of single cells as a marginal space measure *ρ*within the varifold measure framework^6^. STalign then convolves the space measure *ρ*with Gaussian kernels *k*to obtain the smooth, rasterized function 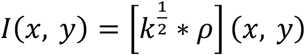. Finally, STalign samples from the continuous *I*(*x,y*) to get a discrete image of a specified size with a specified pixel resolution. STalign focuses on minimizing the dissimilarity between the source and target images *I*^*S*^ and *I*^*T*^ rather than minimizing the dissimilarity between the source and target space measures because, while approximately equivalent, the former can be calculated more efficiently (Online Methods). To solve for a mapping that minimizes the dissimilarity between source and target images *I*^*S*^ and *I*^*T*^, STalign utilizes the LDDMM framework (Fig 1c). Using LDDMM to identify a diffeomorphic solution allows us to have a smooth, continuous, invertible transformation which permits mapping back and forth from the rasterized image and original cell positions while respecting the biological constraints such that cell neighbor relationships stay relatively the same^7^. The mapping *ϕ*^*A,v*^ is constructed from two transformations, an affine transformation A and a diffeomorphism 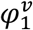 such that 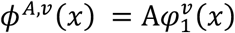, where 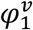 is generated by integrating a time varying velocity field *v*_*t*_ over time and A acts on 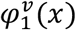 through matrix vector multiplication in homogeneous coordinates. The optimal *ϕ*^*A,v*^ is computed by minimizing an objective function that is the sum of a regularization term, *R*(*v*) and a matching term, *M*_*Έ*_ (*ϕ*^*A,v*^ · *I*^*S*^, *I*^*T*^). The relative weights of the regularization term and matching term can be tuned with 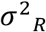and 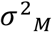. The regularization term controls spatial smoothness. In this term, we optimize over *v*_*t*_ t ∈ [0, 1] noting that if *v*_*t*_ is constricted to being a smooth function, the 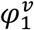 constructed from *v*_*t*_ is guaranteed to be diffeomorphic. The matching term incorporates a Gaussian mixture model *W*(*x*) to estimate matching, background, and artifact components of the image to account for missing tissue such as due to partial tissue matches or tears. Additionally, the matching term contains an image contrast function *f*_θ_ to account for differences due to variations in cell density and/or imaging modalities. To solve all parameters in each term a steepest gradient descent is performed over a user-specified number of epochs. Once *ϕ*^*A,v*^ is computed, STalign applies this computed transformation to the source’s original cell positions (*x*^*ρS*^, *y*^*ρS*^) to generate aligned source coordinates (*x*^*ρS A*^, *y*^*ρ SA*^) (Fig 1d).

**Figure 1.**
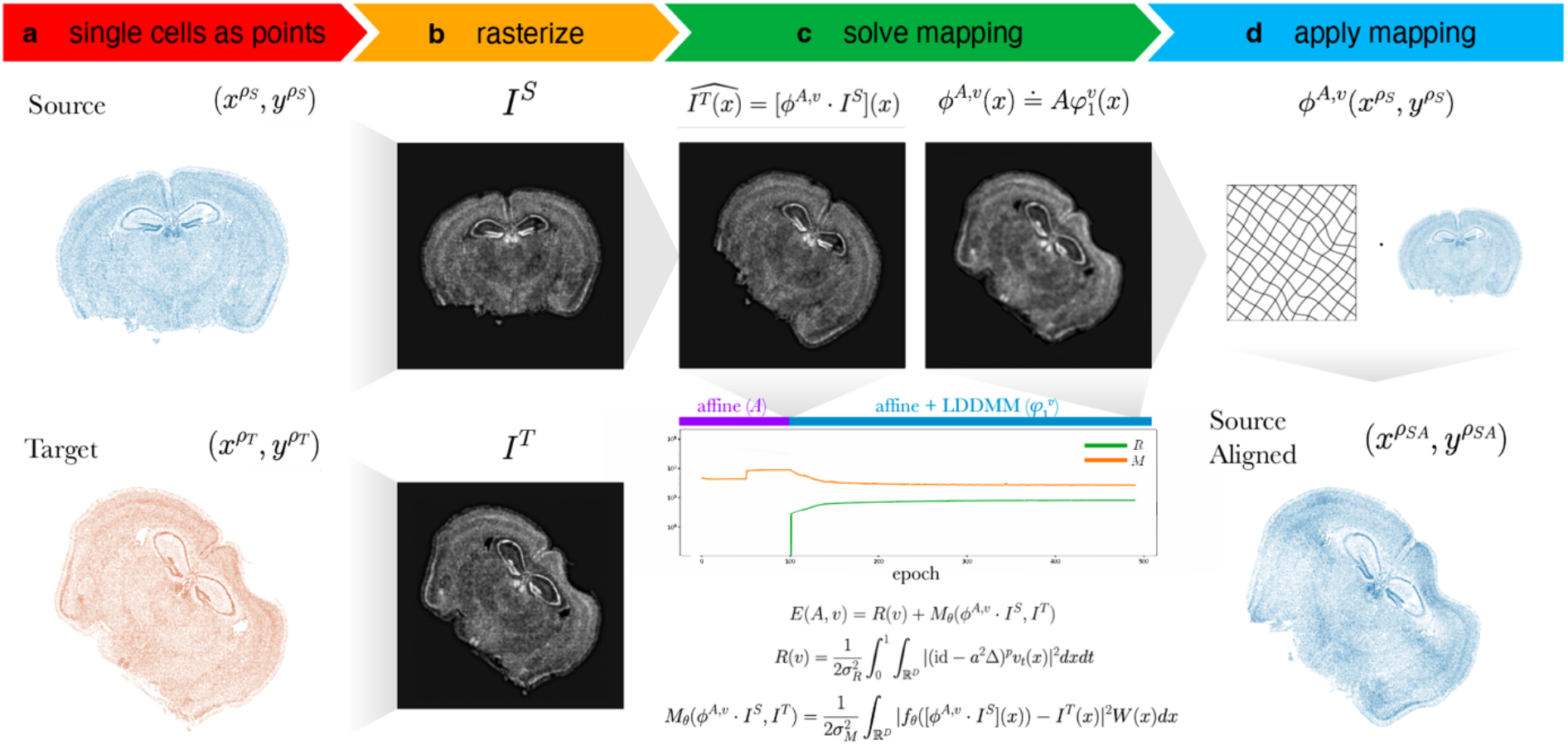
Overview of STalign on ST data from a single-cell resolution technology. **a**. STalign takes as input a source and target ST dataset as x- and y-coordinates of cellular positions. **b**. Source and target coordinates are then rasterized into images *I*^*S*^ and *I*^*T*^. **c**. To align *I*^*S*^ and *I*^*T*^, STalign solves for the mapping *ϕ*^*A,v*^ that when applied to *I*^*S*^ estimates *I*^*T*^ such that *I*^*T*^(*x*) *=* [*ϕ*^*A,v*^ · *I*^*S*^](*x*). Gradient descent is used to solve affine transformation *A* and large deformation diffeomorphic metric mapping (LDDMM) 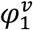 that compose *ϕ*^*A,v*^ such that 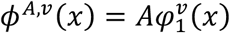. The objective function minimized includes a regularization term *R*(*v*) to penalize non-smooth solutions and a matching term *M*_*Έ*_(*ϕ*^*A,v*^ · *I*^*S*^, *I*^*T*^) that minimizes the dissimilarity between the transformed source image and the target image while accounting for tissue and technical artifacts with *W*(*x*) and *f*_*Έ*_, respectively. Balance between regularization and matching accuracy can be tuned with the parameters 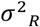 and 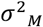. Components of the objective function decrease over epochs with transforms at different stages of the diffeomorphism. **d**. Once *ϕ*^*A,v*^ is solved, visualized as a deformation field, the mapping is applied to the coordinates of the source to obtain the coordinates for the aligned source.

### STalign enables alignment of single-cell resolution ST datasets within technologies

As a proof of concept, we first applied STalign to align two single-cell resolution ST datasets from the same technology. Specifically, we aligned, in a pairwise manner at matched locations, ST data from 9 full coronal slices of the adult mouse brain representing 3 biological replicates spanning 3 different locations with respect to bregma assayed by MERFISH (Methods). Inherent local spatial dissimilarities between slices, due to biological variability and further exacerbated by technical variation as well as tears and distortions sustained in the data acquisition process, render affine transformations such as rotations and translations often insufficient for alignment.

To evaluate the performance of STalign, we first evaluated the spatial proximity of manually identified structural landmarks between the source and target ST datasets, expecting the landmarks to be closer together after alignment. We manually placed 12 to 13 landmarks that could be reproducibly identified (Supp Fig 1, Supp Table 1). To establish a supervised affine transformation for comparison with STalign, we solved for the affine transformation that minimized the error between these landmarks using least squares. We then compared the positions of the corresponding landmarks after both the supervised affine alignment and STalign alignment using root-mean-square error (RMSE). When the supervised affine transformations were used for alignment, RMSE was 202 +/-17.1 μm, 170 +/-3.47 μm, and 266 +/-6.65 μm for biological replicates of each slice location respectively. When STalign based on an LDDMM transformation model was used for alignment, RMSE was 113 +/-10.5 μm, 169 +/-4.53 μm, and 175 +/-5.47 μm for biological replicates of each slice location respectively. STalign was thus able to consistently reduce the RMSE between landmarks after alignment compared to an affine transformation, suggestive of higher alignment accuracy.

Given the ambiguity of where landmarks may be manually reproducibly placed and their inability to evaluate alignment performance for the entire ST dataset, we next took advantage of the available gene expression measurements to further evaluate the performance of STalign. Because of the highly prototypic spatial organization of the brain, we expect high gene expression correspondence across matched spatial locations after alignment. We focused our evaluation on one pair of ST datasets of coronal slices from matched locations (Methods). We visually confirm that alignment results in a high degree of spatial gene expression correspondence (Fig 2a, Supp Fig 2a). To further quantify this spatial gene expression correspondence, we evaluated the gene expression magnitudes at matched spatial locations across the aligned ST datasets. Specifically, we aggregated cells into pixels in a 200μm grid to accommodate the differing numbers of cells across slices and then quantified gene expression magnitude correspondence at spatially matched 200μm pixels using cosine similarity (Fig 2b-c, Supp Fig 2b). For a good alignment, we would expect a high cosine similarity approaching 1, particularly for spatially patterned genes. To identify such spatially patterned genes, we applied MERINGUE^8^ to identify 457 genes with highly significant spatial autocorrelation (Methods). For these genes, we observe a high spatial correspondence after alignment as captured by the high median cosine similarity of 0.73. In contrast, for the remaining 192 non-spatially patterned genes, we visually confirm as well as quantify the general lack of spatial correspondence (Fig 2d-f, Supp Fig 3a-b). We note that these non-spatially patterned genes are enriched in negative control blanks (57%), which do not encode any specific gene but instead represent noise such that we would not expect spatial correspondence even after alignment. Further, we observe a low median cosine similarity of 0.21 across non-spatially patterned genes that is significantly lower than for spatially patterned genes (Wilcoxon rank-sum test p-value < 2.2e-16).

**Figure 2.**
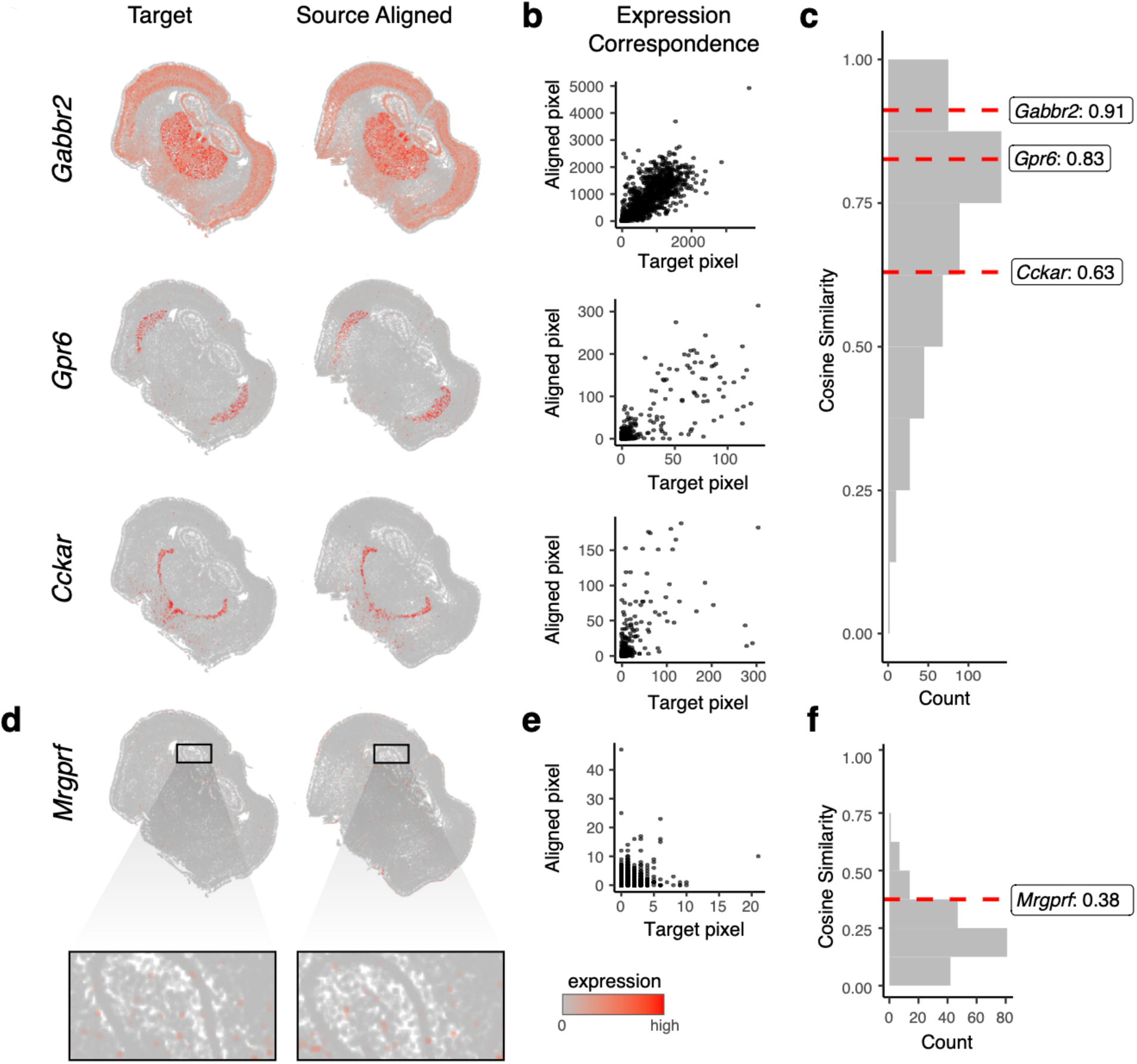
Evaluation of STalign based on spatial gene expression correspondence. **a**. Correspondence of gene expression spatial organization between the target and aligned source for select spatially patterned genes. **b**. Transcript counts in the target compared to the aligned source at matched pixels for select genes: *Gabbr2, Gpr6*, and *Cckar*. **c**. Distribution of cosine similarities between transcript counts in target versus aligned source at matched pixels for 457 spatially patterned genes with select genes marked. **d**. Spatial pattern of expression for a select non-spatially patterned gene in the target and aligned source (inset displays cells at higher magnification). **e**. Counts for the target versus aligned source at matched pixels for a select non-spatially patterned gene, *Mrgprf*. **f**. Distribution of cosine similarities between counts in target compared to the aligned source at matched pixels for 192 non-spatially patterned genes.

**Figure 3.**
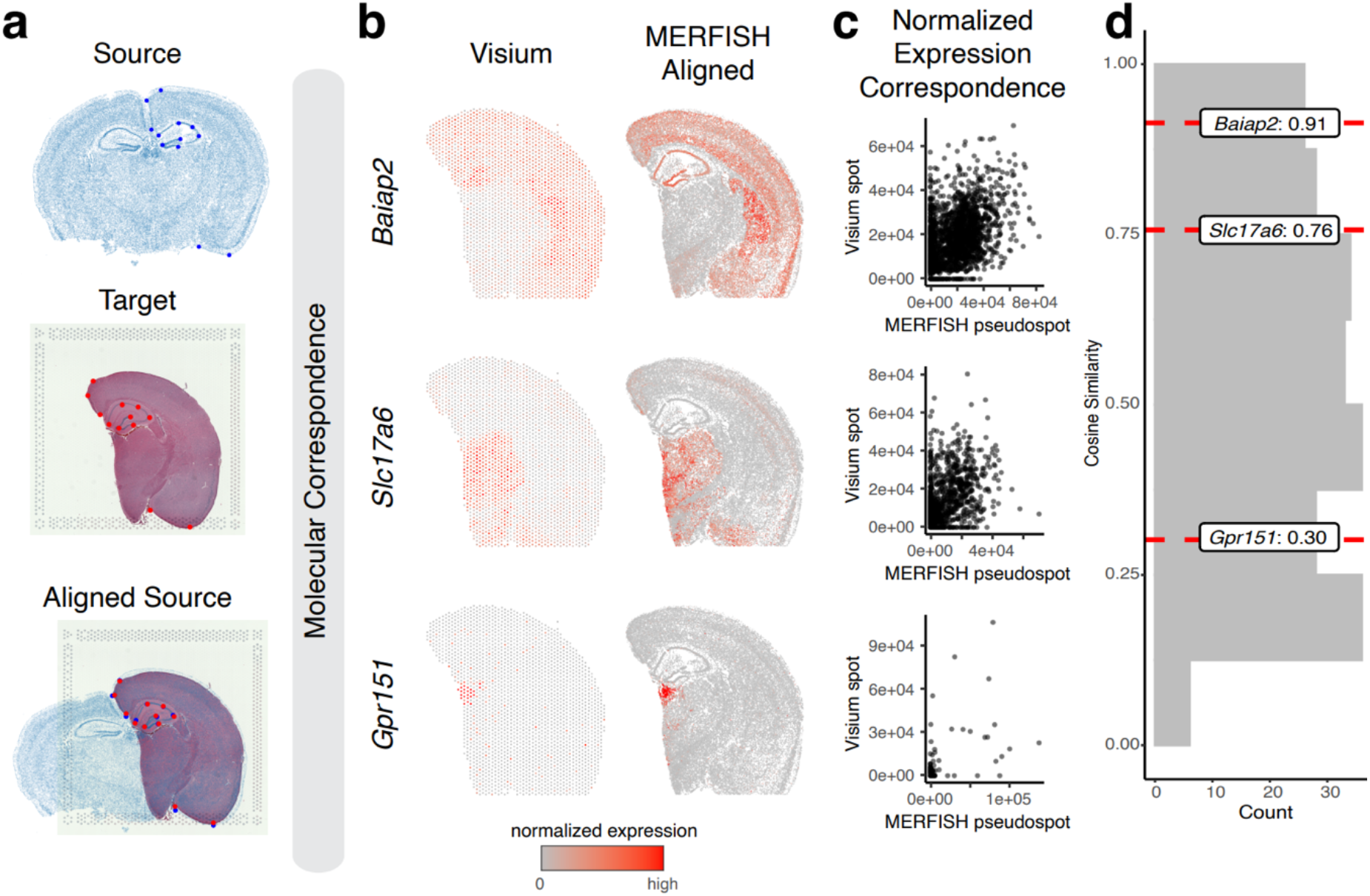
Application and evaluation of STalign on spatial transcriptomics data from different ST technologies based on normalized spatial gene expression correspondence. **a**. Overview of STalign on ST data from different ST technologies. Single-cell resolution ST is used as the source, with the initial image being produced from the x- and y-coordinates of each cell’s position (top). For the multi-cellular resolution ST technologies, the corresponding single-cell resolution histological image is used as target (middle). STalign aligns the source to target (bottom). The manually placed landmarks that were utilized to improve alignment for these partially matched tissues are marked. **b**. Correspondence of gene expression spatial organization between the Visium target and aligned MERFISH source for select spatially patterned genes. **c**. Normalized gene expression in the Visium target compared to the aligned MERFISH source at matched spots and pseudospots respectively for select spatially patterned genes: *Baiap2, Slc17a6* and *Gpr151*. **d**. Distribution of cosine similarities between normalized gene expression in the Visium target versus aligned MERFISH source at matched spots and pseudospots for 227 spatially patterned genes detected by both ST technologies with select genes marked.

We next compare the alignment achieved with STalign to the alignment from a supervised affine transformation based on our previously manually placed landmarks (Supp Fig 4a, Methods). We visually confirm that a supervised affine alignment results in a lower degree of spatial gene expression correspondence than alignment by STalign (Supp Fig4b). We again evaluate performance of the supervised affine transformation using a pixel-based cosine similarity quantification (Supp Fig4c). We find that for spatially patterned genes, the cosine similarity is consistently higher with a mean difference of 0.09 for the alignment by STalign compared to supervised affine (Supp Fig 4d). In contrast, for non-spatially patterned genes, the cosine similarity is more comparable with a mean difference of 0.02 for the alignment by STalign compared to supervised affine (Supp Fig 4e). This greater improvement in spatial gene expression correspondence for the alignment achieved with STalign compared to supervised affine transformation for spatially patterned genes suggests that modeling non-linearity in alignment with approaches like STalign can achieve a higher alignment accuracy compared to linear alignment approaches.

**Figure 4.**
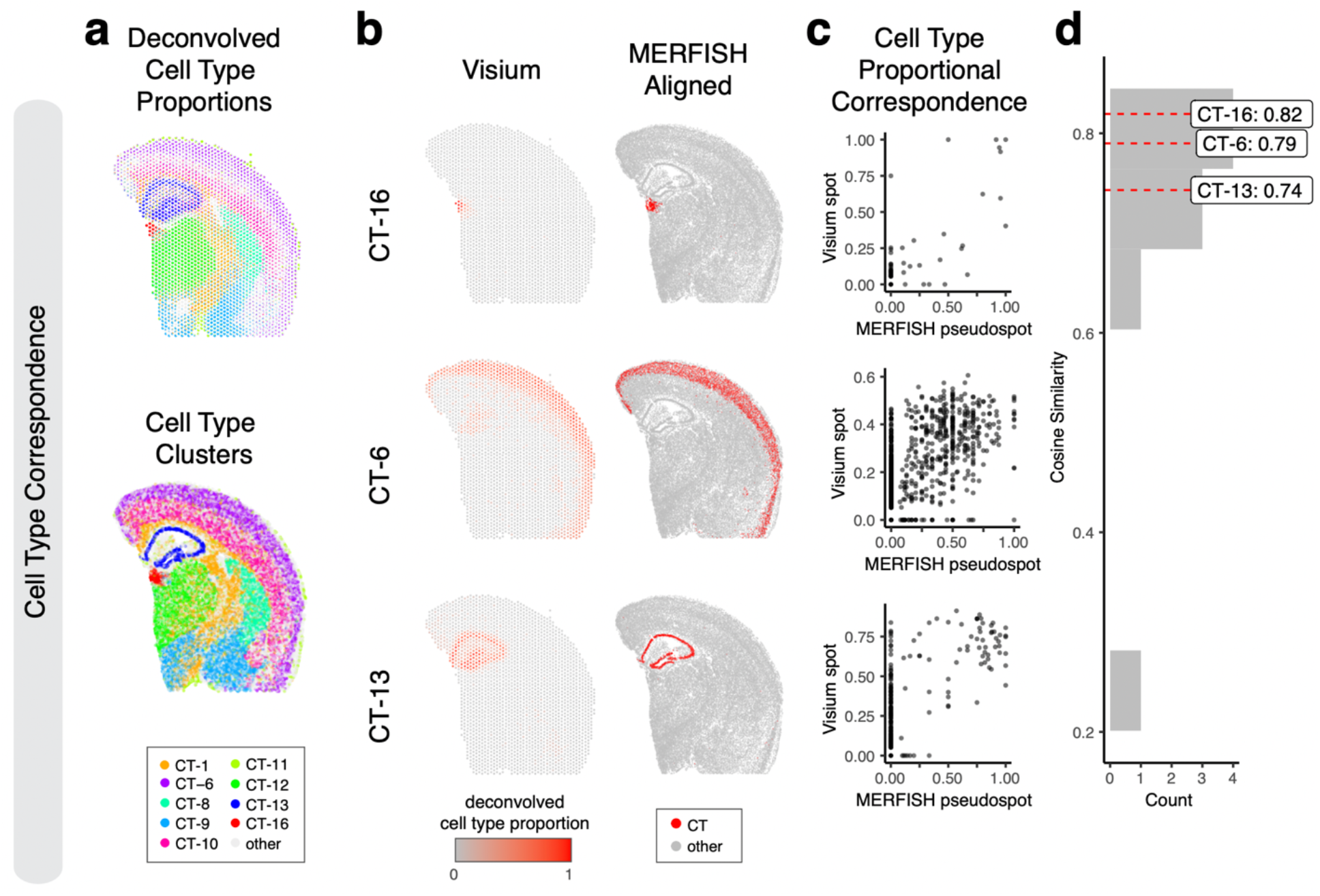
Evaluation of STalign on ST data from technologies at different resolutions based on cell-type correspondence. **a**. Transcriptionally matched cell-types from deconvolution analysis of spot-resolution Visium data (top) and clustering analysis of spatially aligned single-cell-resolution MERFISH data (bottom). **b**. Cell type correspondence between the Visium target and aligned MERFISH source with select cell-types shown. **c**. Correspondence of cell-type proportion between the Visium target and aligned MERFISH source at matched spots and pseudospots respectively for select cell-types. **d**. Distribution of cosine similarities between cell-type proportions in the Visium target and aligned MERFISH source at matched spots and pseudospots respectively for all matched cell-types with cell-types marked.

### STalign enables alignment of ST datasets across technologies

Many technologies for spatially resolved transcriptomic profiling are available, varying in experimental throughput and spatial resolution^9^. We thus applied STalign to align two ST datasets from two such different ST technologies. Specifically, we applied STalign to align the previously analyzed single-cell resolution ST dataset of a full coronal slice of the adult mouse brain assayed by MERFISH to a multi-cellular pixel resolution ST dataset of an analogous hemi-brain slice assayed by Visium (Fig 3a). As such, in addition to being from different ST technologies, these two ST datasets further represent partially matched tissue sections. Because of this partial matching, we incorporated manually placed landmarks to initialize the alignment as well as further help steer our gradient descent towards an appropriate solution (Online Methods). For the Visium dataset, we leveraged a corresponding registered single-cell resolution hematoxylin and eosin (H&E) staining image obtained from the same tissue section for the alignment (Methods).

To evaluate the performance of this alignment, we again take advantage of the available gene expression measurements. Due to partially matched tissue sections, we restricted downstream comparisons to tissue regions STalign assessed with a matching probability > 0.85 (Methods). We again visually confirm that the spatial alignment results in a high spatial gene expression correspondence albeit at differing resolutions across the two technologies (Fig 3b, Supp Fig 5a). To further quantify this spatial gene expression correspondence, we evaluated the gene expression magnitudes at matched spatial locations across the aligned tissue sections for the 415 genes with non-zero expression in both ST datasets. We evaluated these genes for spatial autocorrelation on the Visium data to identify 227 spatially patterned genes and 188 non-spatially patterned genes (Methods). Due to the resolution differences between the two technologies, to ensure appropriate comparisons, we used the positions of the Visium spots to aggregate MERFISH cells into matched resolution pseudospots. Likewise, to control for detection efficiency differences between the two technologies, we performed the same counts-per-million normalization on the Visium spot gene expression measurements and the aggregated MERFISH pseudospots gene expression measurements (Fig 3c, Supp Fig 5b). We again evaluated gene expression correspondence at spatially matched spots using cosine similarity and observed a median cosine similarity of 0.55 across spatially patterned genes (Fig 3d) and a median cosine similarity of 0.06 across non-spatially patterned genes (Supp Fig 6). We note that this gene expression correspondence after spatial alignment is lower than what was previously observed within technologies most likely due to variation in detection efficiency across technologies in addition to variation in tissue preservation rather than poor spatial alignment. While MERFISH detects targeted genes at high sensitivity, Visium enables untargeted transcriptome-wide profiling though sensitivity for individual genes may be lower^9^. Likewise, while the MERFISH dataset was generated with fresh, frozen tissue, the Visium dataset was generated with FFPE preserved tissue. Still, we anticipate that while sensitivity to specific genes may vary across technologies and with different tissue preservation techniques, the underlying cell-types should be consistent.

**Figure 5.**
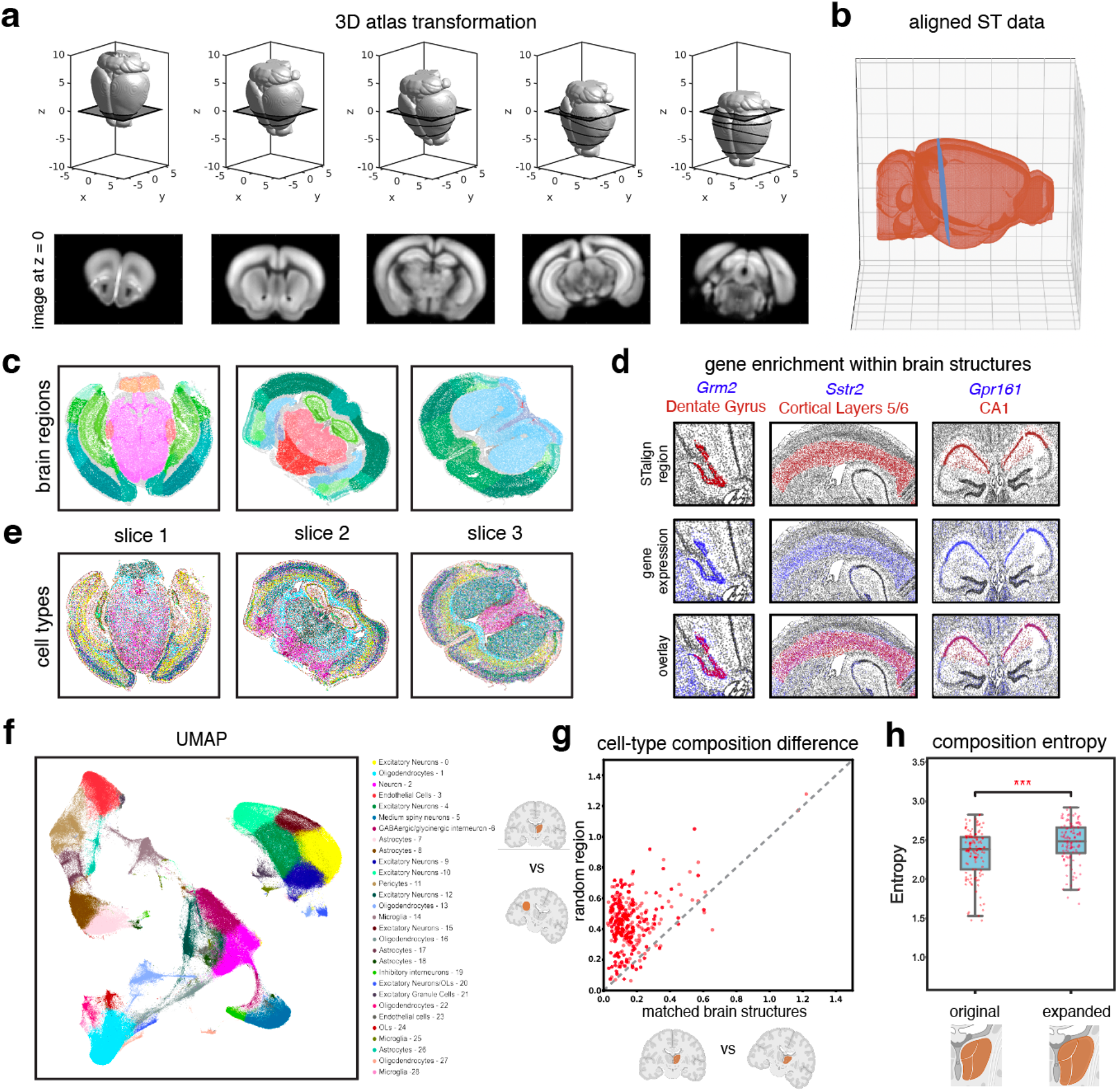
Evaluation of 3D-2D alignment using STalign. **a**. Transformation of 3D CCF atlas to align to ST data at z=0. **b**. Aligned ST data (MERFISH Slice 1 Replicate 1) plotted in 3D Allen Brain Atlas coordinates. **c**. Lift-over brain regions from aligning to the Allen Brain Atlas CCF with STalign. **d**. Brain regions (top) labeled by STalign with expression of expected genes (middle) and overlay (bottom). **e**. Spatial location of cell types on MERFISH brain slices. **f**. UMAP embedding of different cell types defined by differential gene expression and Leiden clustering. **g**. Cell-type composition difference between paired brain regions from two MERFISH replicates. The x axis represents cell-type composition difference within matched brain structures annotated by STalign across replicates and the y axis represents cell-type composition difference between STalign-annotated regions and size-matched random brain regions. **h**. Significant difference between distribution of cell-type composition entropy for brain regions labeled by STalign versus regions expanded by 100 nearest neighbors (center line, median; box limits, upper and lower quartiles; whiskers, 1.5x interquartile range; all data points shown)

**Figure 6.**
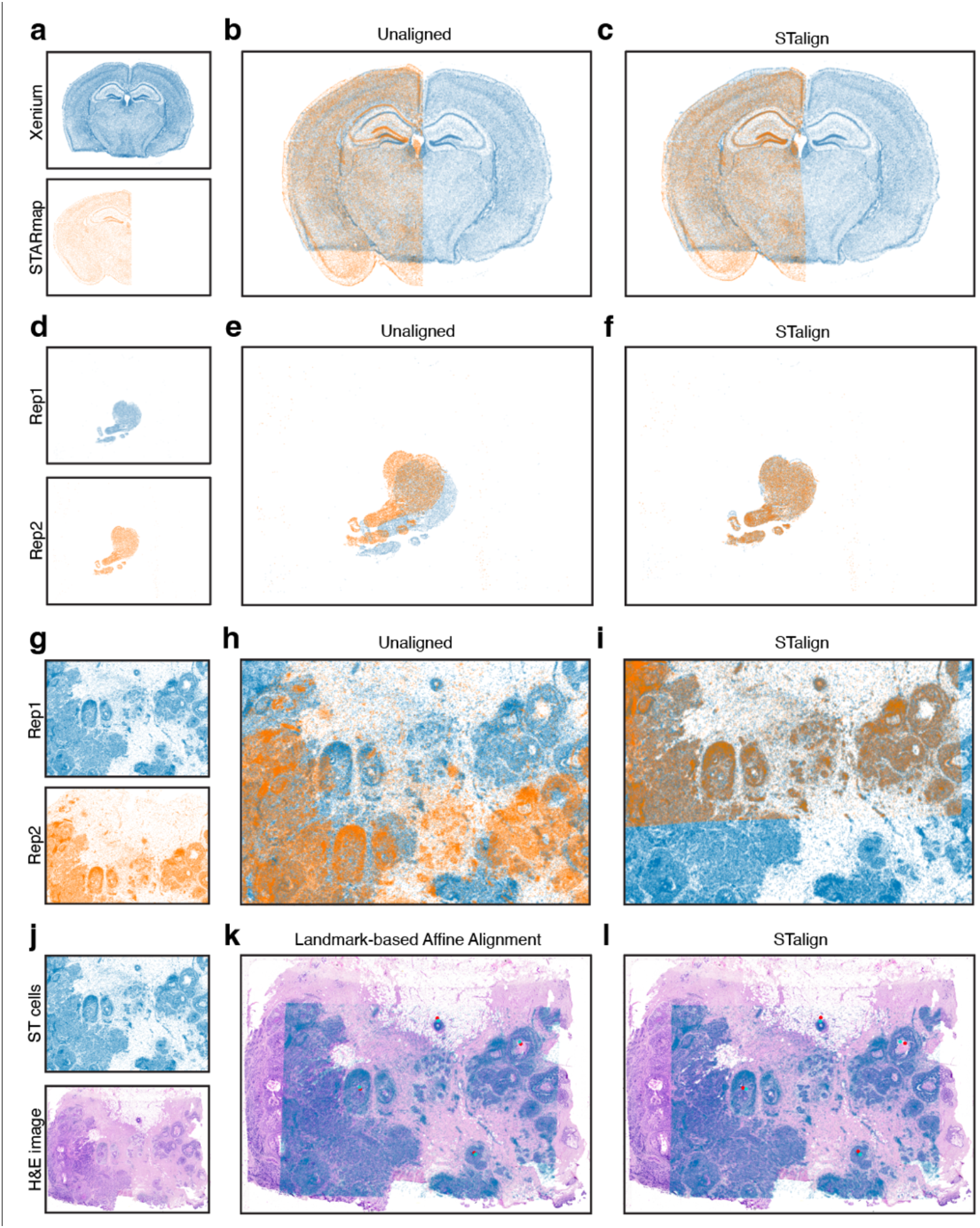
Application of STalign to ST data of diverse tissues. **a**. Two coronal slices of the adult mouse brain assayed by two different single-cell resolution ST technologies, Xenium and STARmap PLUS **b**. Overlay of cellular positions before alignment. **c**. Overlay of cellular positions after alignment with STalign. **d**. Two single-cell resolution datasets from serial sections of the developing human heart. **e**. Overlay of cellular positions before alignment. **f**. Overlay of cellular positions after alignment with STalign. **g**. Two single-cell resolution ST datasets from partially matched, serial breast cancer sections visualized as x- and y-coordinates of cellular positions. **h**. Overlay of cellular positions before alignment. **i**. Overlay of cellular positions after alignment with STalign. **j**. A single-cell resolution ST dataset with a corresponding H&E image from the same tissue section. **k**. Overlay of cellular positions and H&E image based on affine transformation by minimizing distances between manually placed landmarks, shown as points in red and turquoise. **l**. Overlay of cellular positions and H&E image after alignment with STalign.

Therefore, we sought to evaluate the performance of our alignment based on cell-type spatial correspondence. To identify putative cell-types, we performed transcriptional clustering analysis on the single-cell resolution MERFISH data (Supp Fig 7a) and deconvolution analysis^10^ on the multi-cellular pixel-resolution Visium data (Fig 4a, Methods). We matched cell-types based on transcriptional similarity between cell clusters and deconvolved cell-types (Supp Fig 7b). Indeed, we visually observe high spatial correspondence across matched cell-types (Fig 4a-b). We evaluated the proportional correspondence of cell-types at aligned spot and pseudospot spatial locations by cosine similarity and observed a high median cosine similarity of 0.75 across cell-types (Fig 4c-d). As such, STalign achieves high cell-type spatial correspondence across aligned ST datasets, suggestive of high alignment accuracy.

### STalign enables alignment of ST datasets to a 3D common coordinate framework

Tissues are inherently 3-dimensional (3D), and tissue sections are subject to distortions in 3D as well as 2D. As such, a more precise spatial alignment of 2D tissue sections must accommodate this 3D distortion. The underlying mathematical framework for STalign is amenable to alignment in 2D as well as 3D (Online Methods). We thus applied STalign to align ST datasets to a 3D common coordinate framework (CCF). Specifically, we applied STalign to align 9 ST datasets of the adult mouse brain assayed by MERFISH to a 50μm resolution 3D adult mouse brain CCF established by the Allen Brain Atlas^11^ (Methods, Fig 5a). We note that such a 3D alignment can accommodate deformations in and out of 2D planes (Fig 5b). In the construction of the Allen Brain Atlas CCF, brain regions were delineated based on several features like cellular architecture, differential gene expression, and functional properties via modalities such as histological stains, in situ hybridization, and connectivity experiments to generate a set of reference brain region annotations^11^. By aligning to this CCF, we can lift over these annotations to each cell (Fig 5c, Supp Fig 8a), enabling further evaluation of variations of gene expression and cell-type composition within and across these annotated brain regions.

To assess the performance of our atlas alignment and lift-over annotations, we first confirmed the enrichment of genes within certain brain regions. Numerous previous studies have shown that some brain regions can be demarcated based on the expression of particular genes^12,13^. We use these characteristic gene expression patterns to evaluate whether the brain regions lifted over from the Allen Brain Atlas CCF by STalign indeed contain expression of known marker genes. Consistent with previous studies, we found *Grm2* to be visually primarily enriched in the dentate gyrus brain region^14^, *Sstr2* to be enriched in cerebral cortical layers 5 and 6 brain region^15^, and *Gpr161* to be enriched in the CA1 brain region^16^ (Fig 5d), which was consistent across replicates (Supp Fig 8b).

Next, we took a more agnostic approach to assess the performance of our atlas alignment and lift-over annotations by evaluating the consistency of cell-type compositional heterogeneity within brain structures across replicates. To identify cell-types, we perform unified transcriptional clustering analysis on these 9 ST datasets to identify transcriptionally distinct cell clusters and annotate them as cell-types based on known differentially expressed marker genes (Methods, Fig 5e-f, Supp Fig 9a). Many brain regions are known to have a characteristic cell type distribution^17– 19^. Consistent with previous studies^20^, we observed cell-types to be spatially and compositionally variable across brain regions (Fig5c, Fig 5e). We visually confirmed that this spatial and compositional variability is consistent across replicates (Supp Fig8a, Supp Fig 9b). To further quantify this consistency, for each brain region, we evaluated whether its cell-type composition was more similar between replicates than compared to a randomly demarcated brain region of matched size (Methods). For an accurate atlas alignment, we would expect the lift-over brain region annotations to be more similar in cell-type composition across replicates, particularly for brain structures with distinct cell-type compositions, as compared to random brain regions of matched size. Indeed, we found that in 93% of evaluated brain structures (131/141), the cell-type composition was significantly more similar (Paired t-test p-value = 6.805e-121) between replicates than compared to a random brain region of matched size. (Fig 5g). For the 7% (10/141) of brain regions that were less similar across replicates, we found that the number of cells in these brain regions were significantly fewer (Wilcoxon rank-sum test p-value = 0.002) than other brain regions (Supp Fig 10a). Notably, 60% of these brain regions had a minimum width of under 50μm, including both compact and long, thin structures (Supp Fig 10b), highlighting potential limitations with respect to alignment accuracy of such structures at this given resolution of alignment.

Finally, we also sought to assess the performance of our atlas alignment and lift-over annotations by evaluating cell-type compositions within and beyond annotated brain region boundaries (Methods). Specifically, we compare the entropy of each brain region based on the region’s cell-type composition to entropy if the boundaries of these regions were expanded (Fig 5h). Again, due to the characteristic cell-type distributions within brain regions in which one or a few cell-types predominate, we would expect accurate lift-over brain region annotations to exhibit entropies that are comparatively lower than if the boundaries of these regions were expanded, as more cell-types would be incorporated into the region and entropy would increase. We therefore expanded the brain structures lifted over by STalign by 100 nearest neighbors (NN), or approximately 100μm, and evaluated the change in entropy. We performed the same analysis on randomly demarcated brain regions of matched size, which were expanded by 100 NN to account for increases in entropy due to an incorporation of more cells. We found that the entropies for the original brain region annotations lifted over by STalign were significantly lower (paired t-test p-value=8.6e-18) than for the expanded regions. In contrast, the entropies for random brain regions were not significantly lower (paired t-test p-value = 0.12) for the expanded regions (Supp Fig 11). Taken together, these results demonstrate that STalign can align ST datasets to a 3D CCF to consistently lift over atlas annotations that recapitulate the unique gene expression and cell-type composition within brain regions.

### STalign applicable to diverse tissues profiled by diverse ST technologies

STalign relies on variation in cell densities that generally form visible structures that can be used for alignment. As we have shown, alignment across samples and animals is possible for tissues with highly prototypic structures such as the brain. We further highlight the applicability of STalign to the diverse ST technologies that can assay this tissue by demonstrating that we can apply STalign to achieve structural correspondence for coronal slices of the adult mouse brain assayed by two different single-cell resolution ST technologies, Xenium^21^ and STARmap PLUS^22^ (Methods, Fig 6a-c).

For other tissues with substantially more inter-sample and inter-animal variation, alignment across serial sections is still achievable. For example, for serial sections of the developing human heart^23^, we can apply STalign to achieve structural correspondence (Methods, Fig 6d-f). Likewise, even for cancer tissues, which are highly non-prototypic in structure, there is still often sufficient structural consistency across serial sections to enable alignment. As such, we have applied STalign to align single-cell resolution ST datasets arising from partially matched serial sections of the same breast cancer sample assayed by Xenium (Methods, Fig 6g-i). Likewise, we have applied STalign to align a single-cell ST dataset assayed by Xenium to a corresponding H&E image of the same tissue section (Methods, Fig 6j-l). We visually observe a high degree of spatial correspondence and overlap of structural features after alignment, highlighting STalign’s applicability to diverse tissues.

## Discussion

Alignment of ST datasets is a prerequisite step to enable comparisons across samples, subjects, and technologies. Alignment can also enable pooling of measurements across biological replicates to construct consensus ST profiles^1^ as well as enable 3D reconstruction by serial registration^24^. Here, we presented STalign, which builds on advancements in LDDMM, to perform alignment of ST datasets in a pairwise manner within ST technologies, across ST technologies, as well as to a 3D common coordinate system. We have shown that STalign achieves high accuracy based on the spatial proximity of manually identified shared landmarks as well as gene expression and cell-type correspondence at matched spatial locations after alignment. We note that based on these metrics, STalign outperforms affine transformations alone, highlighting the utility of local, non-linear transformations in alignment. STalign can further accommodate partially matched tissue sections, where one tissue section may be a fraction of another. We further apply STalign to align ST datasets to a 3D CCF to enable automated lift-over of CCF annotations such as brain regions in a scalable manner. We confirm that lift-over brain region annotations identify cells that express expected genes for a variety of brain regions. We also show that brain region annotations lifted over by STalign exhibit consistent cell-type compositions across replicates and within boundaries compared to random brain regions matched in size.

We anticipate that future applications of STalign to ST data particularly across ST technologies will enable cross-technology comparisons as well as cross-technology integration through spatial alignment. In particular, aligning ST data for similar tissues across different ST technology platforms may allow us to better interrogate platform-specific differences and strengths. Given that different ST technologies currently generally prioritize either resolution or genome-wide capabilities, we may wish to apply different ST technologies on serial sections to leverage their unique strengths to characterize matched spatial location. With atlasing efforts like The Human BioMolecular Atlas Program and others producing 3D CCFs^25^, application of STalign to align ST data to such CCFs to enable automated lift over of atlas structural annotations will facilitate standardization and unification of biological insights regarding annotated structures. Likewise, STalign complements gene-expression-based approaches for sample alignment^26^ by focusing on the real space rather than a higher-order transcriptomic manifold. We further anticipate future applications of STalign to ST data from structurally matched tissues in case-control settings will enhance the throughput for yielding meaningful comparisons regarding gene expression and cell-type distributions in space as evidenced by recent applications of ST technologies to characterize spatially-resolved age-related^27^ and injury-related^28^ gene expression variation.

As ST technologies continue to evolve, we anticipate STalign will continue to be applicable due to our use of rasterization to convert the positions of single cells into an image with specified resolution. The runtime of each iteration of the STalign alignment algorithm scales with respect to the number of pixels in this image. For most evaluated datasets, we find that STalign is generally able to converge onto an optimal alignment within a few minutes to a few hours, depending on the number of pixels, the number of iterations, and other system variables (Methods, Supp Table 2). Whereas other alignment algorithms generally scale in memory and runtime with the number of spatially resolved measurements (spots or cells)^1,2^, which will likely make them computationally untenable as ST technologies evolve to increase the number of spatially resolved measurements that can be assayed. Overall, we anticipate that the ability for users to choose the rasterization resolution, and therefore the number of pixels in the rasterized image, will allow STalign to maintain its utility for larger datasets.

Still, among the limitations of STalign with respect to ST data, it is currently applicable to only ST datasets with single-cell resolution or those accompanied with a registered single-cell resolution histology image from same assayed tissue section, which may not be available to all non-single-cell resolution ST technologies. STalign further relies on the representative nature of cell segmentations in ST data to reflect underlying tissue structures. As such, limitations in cell segmentations that render the derived cell density to be no longer representative of the profiled tissue structure could present challenges for alignment with STalign.

Further, as STalign is based on an LDDMM transformation model for alignment, it inherits the same limitations. As LDDMM relies on optimization using gradient descent, the resulting alignment solution may converge on local minima. Strategies to guide the optimization away from potential local minima may be applied in the future. Likewise, the more different the source and targets for alignment, particularly for partially matching sections, the more important the initialization will be for this optimization. As we have shown, landmark points may be used to guide the initialization of an orientation and scaling for alignment. In addition, LDDMM enforces an inverse consistency constraint such that every observation in the target must have some correspondence in the source in a manner that cannot accommodate holes or other topological differences in the tissue through the deformation only^7^. As such, when performing alignments, we advise choosing the more complete tissue section as the source because our Gaussian mixture modeling for accommodating partially matched tissues and other artifacts applies to the target image intensity only.

Still, alignment accuracy at the resolution of single cells is limited by the fact that there is generally no one-to-one correspondence between cells across samples, particular for complex tissues. As such, accuracy can typically only be expected to be achieved up to a “mesoscopic scale” at which it is reasonable to define cell density^29^. As we have shown, this presents challenges particularly in aligning thin structures. While STalign currently uses an isotropic (Gaussian) kernel to estimate cell densities, future work considering non-isotropic kernels may improve accuracy for these thin structures. However, generally, our choice of kernel will inherently bias our alignment towards accuracy at a certain structural scale. Likewise, although we focused here on aligning based on cell densities, STalign and the underlying LDDMM framework can also be applied to align using cellular features such as gene expression magnitude, reduced dimensional representations of gene expression such as via principal components, or cell-type annotations, which may improve the accuracy of alignment for regions with homogenous cell density but heterogeneous gene expression and cell-type composition. However, integration of such features in the alignment process necessitates orthogonal means of performance evaluation beyond the correspondences in gene expression magnitude and cell-type proportions that we have used here. By aligning based on cell densities, we do not require shared gene expression quantifications or unified cell-type annotations, potentially enhancing flexibility and providing opportunities for integrating across other data modalities for which spatially resolved single cell resolution information is available such as other spatial omics data in the future.

Overall, we anticipate that moving forward STalign will help provide a unified mathematical framework for ST data alignment to enable integration and downstream analyses requiring spatial alignment to reveal new insights regarding transcriptomic differences between different tissue structures and across various physiological axes.

## Supporting information

supplementary software

## Acknowledgments

KC, MA, GA, LA, and JF are supported by the National Institute of General Medical Sciences of the National Institutes of Health under Award Number R35-GM142889 and the National Science Foundation under Grant No. 2047611. MA and JMK are supported by the National Institute of Drug Abuse under Award Numbers DP1-DA056668. Fig. 5 and Supp Fig. 11 were created with component graphics licensed from Biorender.com.

## Author Contributions

DT led the development of the STalign software and mathematical modeling with input from JF, KC, MA, and MIM. JF, KC, and MA led the application of STalign to various ST datasets with input from DT. KC evaluated the performance of STalign for 2D alignment under the guidance of JF. MA evaluated the performance of STalign for 3D alignment under the guidance of JF and JMK. OKA evaluated the performance of STalign using landmark-based approaches under the guidance of DT. GA performed runtime benchmarks and code revisions under the guidance of KC, DT, and JF. LA contributed to the revision under the guidance of KC and JF. All authors contributed to the writing of the manuscript. All authors approved the final manuscript.

## Methods

### Datasets

Nine MERFISH datasets consisting of 734,696 cells and 483 total genes, across 9 brain slices (3 replicates of 3 coronal sections from matched locations with respect to bregma) were obtained from the Vizgen website for *MERFISH Mouse Brain Receptor Map data release* (https://info.vizgen.com/mouse-brain-map).

A Visium dataset of an FFPE preserved adult mouse brain were obtained from the 10X Datasets website for *Spatial Gene Expression Dataset by Space Ranger 1*.*3*.*0* (https://www.10xgenomics.com/resources/datasets/adult-mouse-brain-ffpe-1-standard-1-3-0)

A Xenium dataset (In Situ Replicate 1) of a fresh frozen mouse brain coronal section was obtained from the 10X Datasets website for *Mouse Brain Dataset Explorer* (https://www.10xgenomics.com/products/xenium-in-situ/mouse-brain-dataset-explorer)

STARMAP Plus data (well11_spatial.csv) of coronal slices of the adult mouse brain was downloaded from the Broad Single Cell Portal (https://singlecell.broadinstitute.org/single_cell/study/SCP1830/spatial-atlas-of-molecular-[…]pes-and-aav-accessibility-across-the-whole-mouse-brain)

Developing heart data for samples CN73_E1 and CN73_E2 were downloaded from the Human Developmental Cell Atlas https://hdca-sweden.scilifelab.se/a-study-on-human-heart-development via ST_heart_all_detected_nuclei.RData from https://github.com/MickanAsp/Developmental_heart

Two Xenium datasets (In Situ Replicate 1 and In Situ Replicate 2) of a single breast cancer FFPE tissue block were obtained from the 10X Datasets website for *High resolution mapping of the breast cancer tumor microenvironment using integrated single cell, spatial and in situ analysis of FFPE tissue* (https://www.10xgenomics.com/products/xenium-in-situ/preview-dataset-human-breast)

The 50μm resolution 3D adult mouse brain CCF was obtained from the Allen Brain Atlas website (https://download.alleninstitute.org/informatics-archive/current-release/mouse_ccf/annotation/ccf_2017/annotation_50.nrrd).

### Application of STalign

To align MERFISH datasets, we applied STalign in a pairwise manner across replicates for sections from matched locations with respect to bregma, rasterized at a 50μm resolution, and iterated over 1000 epochs, with the following changes to default parameters (sigmaM: 0.2).

To align a MERFISH dataset to a Visium dataset, we applied STalign with MERFISH Slice 2 Replicate 3, rasterized at a 50μm resolution, as the source and the high resolution Visium hematoxylin and eosin (H&E) staining image as the target. We utilized the landmark points stored in Merfish_S2_R3_points.npy and tissue_hires_image_points.npy as inputs pointsI and pointsJ. We iterated for 200 epochs with the following changes to default parameters (sigmaP:

0.2, sigmaM: 0.18, sigmaB: 0.18, sigmaA: 0.18, diffeo_start: 100, epL: 5e-11, epT: 5e-4, epV:5e1).

To align MERFISH to the Allen CCF, we applied STalign using the 3D reconstructed Nissl image from the Allen CCF atlas as a source, and each of our 9 MERFISH images as a target.

To align Xenium and STARmap datasets of mouse brain coronal sections, we applied STalign with Xenium In Situ Replicate 1, rasterized at 30 μm resolution, as the source and STARmap well 11, rasterized at 30 μm resolution, as the target. Prior to rasterization, STARmap cell centroid positions were scaled by 1/5 such that the overlay of unaligned sections showed both Xenium and STARmap cells positions at a similar scale. We iterated for 1000 epochs with the following changes to default parameters (sigmaM:1.5, sigmaB:1.0, sigmaA:1.5, epV: 100, muB: black).

To align serial developing heart sections, we applied STalign with sample CN73_E1 as the source and CN73_E2 as the target, both rasterized at 100 μm resolution. We iterated for 1000 epochs with the following changes to default parameters (diffeo_start:100, a: 250, sigmaB:0.1, epV: 1000, muB: black).

To align Xenium datasets, we applied STalign with Xenium Breast Cancer Replicate 1 as the source and with Xenium Breast Cancer Replicate 2 as the target, rasterized at 30μm resolution. We placed a set of 3 manually chosen landmark points to compute an initial affine transformation.

We iterated for 200 epochs with the following changes to default parameters (sigmaM:1.5, sigmaB:1.0, sigmaA:1.5, epV: 100).

To align Xenium to H&E, we applied STalign with Xenium Breast Cancer Replicate 1, rasterized at 30μm resolution, as the source and the corresponding H&E image from the same tissue as the target. We placed a set of 3 manually chosen landmark points to compute an initial affine transformation. We iterated for 2000 epochs with the following changes to default parameters (sigmaM:0.15, sigmaB:0.10, sigmaA:0.11, epV: 10, muB: black, muA: white) where muB and muA initializes the mixture model for the background and artifact components as corresponding to black and white colors respectively in the target image.

### Expression based performance evaluation for STalign-based alignment of single-cell resolution ST datasets within technologies

To evaluate the performance of STalign on aligning datasets from the same technologies based on expression correspondence, we focused on the alignment of Slice 2 Replicate 3 and Slice 2 Replicate 2 from the MERFISH datasets, with the former as the source and the latter as the target. A grid was created to partition all cells into 200μm square pixels. For each 200μm pixel, the gene expression of cells in the pixel was summed for the aligned source and for the target to get gene expression at 200μm resolution.

MERINGUE (v1.0) was applied to calculate Moran’s I on the 200μm resolution summed gene expression of the target. Genes with an adjusted p-value < 0.05 were identified as significantly spatially patterned genes and genes with an adjusted p-value >= 0.05 were identified as non-significantly spatially patterned genes.

For each gene, the cosine similarity was calculated between the 200μm resolution summed gene expression counts in the aligned source and the 200μm resolution summed gene expression counts in the target across pixels. A Wilcoxon rank sum test was used to compare the distributions of cosine similarities for spatially patterned and non-significantly spatially patterned genes.

### Comparison to supervised affine alignment of single-cell resolution ST datasets within technologies

In addition to alignment by STalign, we performed supervised affine alignment of Slice 2 Replicate 3 and Slice 2 Replicate 2 from the MERFISH datasets, with the former as the source and the latter as the target. We manually placed 13 landmarks in the source and target that could be reproducibly identified (Supp Fig 1, Supp Table 1) using our script point_annotator.py. We solved for the affine transformation that minimized the error between these landmarks using least squares and applied the affine transformation to the cell positions of the source. With the supervised affine aligned source and target, we repeated the expression-based performance evaluation described in section “Expression based performance evaluation for STalign-based alignment of single-cell resolution ST datasets within technologies.”

### Evaluation alignment across technologies

#### Expression based performance

Given that the MERFISH tissue section is larger than the Visium, we considered the aligned region to be limited to the MERFISH tissue that had a matching probability > 0.85 based on the posterior probability of pixels belonging to the matched class in the Gaussian mixture modeling, with the 0.85 threshold being manually chosen based on visual inspection. We restricted the set of cells in the MERFISH dataset to only those in this aligned region for downstream evaluation.

To aggregate the cells in the aligned MERFISH dataset into pseudospots that match with the Visium spots, we calculated the distances between the positions of the MERFISH cells and the positions of the Visium spot centroids. Cells were classified as within the pseudospot that corresponds to the Visium spot if the distance of the cell to the Visium centroid was less than the Visium spot radius. The Visium spot radius information was obtained by multiplying the ‘spot_diameter_fullres’ by the “tissue_hires_scalef” in the Visium scalefactors_json.json file and dividing by 2. For each pseudospot, the gene expression of all cells within the pseudospot was summed.

For gene expression correspondence analysis, we restricted to the 415 genes that had at least one copy in both the MERFISH and Visium datasets and that were detected in more than one spot in the Visium dataset.

MERINGUE (v1.0) was applied to calculate Moran’s I on the Visium counts-per-million (CPM) normalized counts. Genes with an adjusted p-value < 0.05 were identified as significantly spatially patterned genes and genes with an adjusted p-value >= 0.05 were identified as non-significantly spatially patterned genes.

CPM normalization and log10 transformation with a pseudocount of 1 were applied on the gene expression of the MERFISH pseudospots and Visium spots. For each gene, the cosine similarity was calculated between the normalized and log-transformed gene expression magnitudes across matched MERFISH pseudospots and Visium spots.

#### Cell-type correspondence performance

To identify cell-types in the Visium data, we applied STdeconvolve on a corpus of 838 genes after filtering out lowly expressed genes (<100 copies), genes present in < 5% of spots and genes present in > 95% of spots and restricting to significantly over-dispersed with alpha =1e-16 to obtain a corpus < 1000 genes, resulting in 16 deconvolved cell-types.

To identify cell-types in the aligned MERFISH data, PCA was performed on the CPM normalized cell by gene matrix. Louvain clustering was performed on a neighborhood graph of cells using the top 30 PCs and 90 nearest neighbors to identify 16 transcriptionally distinct clusters of cells.

To match deconvolved cell-types and single-cell clusters, we used the deconvolved cell-type-specific transcriptomic profiles from STdeconvolve and averaged the transcriptional profiles per cluster from single-cell clustering. We restricted to the 257 shared genes, CPM normalized, and correlated the resulting normalized transcriptional profiles using Spearman correlation. We considered a Visium deconvolved cell-type and MERFISH single-cell cluster as a match if they had transcriptional similarity > 0.5.

For each matched cell-type, we evaluated spatial compositional correspondence using cosine similarity of the cell-types proportional representation across matched MERFISH pseudospots and Visium spots.

### Evaluation of 2D to 3D CCF alignment

#### Unified transcriptional clustering analysis and cell-type annotation

All MERFISH datasets were combined. Transcriptional clustering analysis and cell type annotation was performed using the SCANPY package^30^ [version 1.9.1]. Data were normalized to counts per million (scanpy: normalize_total) and log transformed (scanpy: log1p). PCA (scanpy: pca) was computed on the cell by gene matrix. A neighborhood graph of cells using the top 10 PCs and 10 nearest neighbors was created (scanpy: neighbors), and Leiden clustering was performed on this graph (scanpy: leiden) to identify 29 clusters. Differentially expressed genes were extracted from each cluster (scanpy: rank_genes_groups), and cell-types were annotated based on marker genes in each cluster.

#### Annotated brain region composition analysis

To generate randomly demarcated brain regions, a random number generator (random.randint) defined the x, y coordinate of the center of the random region, and the random region was composed of the N closest points to the center, where N is the number of cells in the brain region. A slice/replicate with random regions was constructed for all slice/replicates with STalign annotated regions, and the number of cells (N) were the same for STalign and randomly demarcated regions.

To compare cell-type compositions, each region was represented by a cell-type vector, which was composed by the proportion of each cell type in the region (29×1 vector). We calculate the Euclidean distance between cell type vectors of the same region across replicates in Slice 2 using the regions annotated by STalign. The Euclidean distance was also found across replicates in Slice 2 using randomly demarcated brain regions and STalign brain regions. The Euclidean distances of both groups were compared using a paired t-test. 431 data points for each group were used, comparing replicate 1 to replicate 2, replicate 2 to replicate 3, and replicate 3 to replicate 1.

To evaluate annotated brain region boundaries, brain regions were expanded using k-nearest neighbors (k=100) using the ‘ball tree’ algorithm for each region and each replicate in Slice 2 (sklearn.neighbors.NearestNeighbors). The procedure was conducted for STalign annotated brain regions and randomly demarcated brain regions. Shannon’s entropy was evaluated for STalign annotated and randomly demarcated brain regions that were expanded by 100 nearest neighbors. Paired t-tests were used to compute p-values between original and expanded brain regions for STalign and random groups. Effect size was computed as a difference in the means of the compared distributions. PP plots were used to visualize normality, and we used a Gaussian fit with R>0.8 and a variance ratio less than 4 to confirm normality and equal variances. 431 data points for each group were used.

To evaluate regions that had a greater Euclidean distance between two STalign regions compared to random versus STalign regions, we calculated the number of cells and Shannon’s entropy of each region and tested for significance using a Wilcoxon Rank Sum test due to the small sample size. Shannon’s entropy was calculated using the formula ∑ *p*(*x*) * logE*p*(*x*)F where p(x) is the probability of picking cell-type x from the given brain region (scipy.special.entr).

### Implementation and software availability

STalign is available as an open-source Python toolkit at https://github.com/JEFworks-Lab/STalign and as supplementary software with additional documentation and tutorials available at https://jef.works/STalign.

The implementation of STalign uses the following parameters and default values.

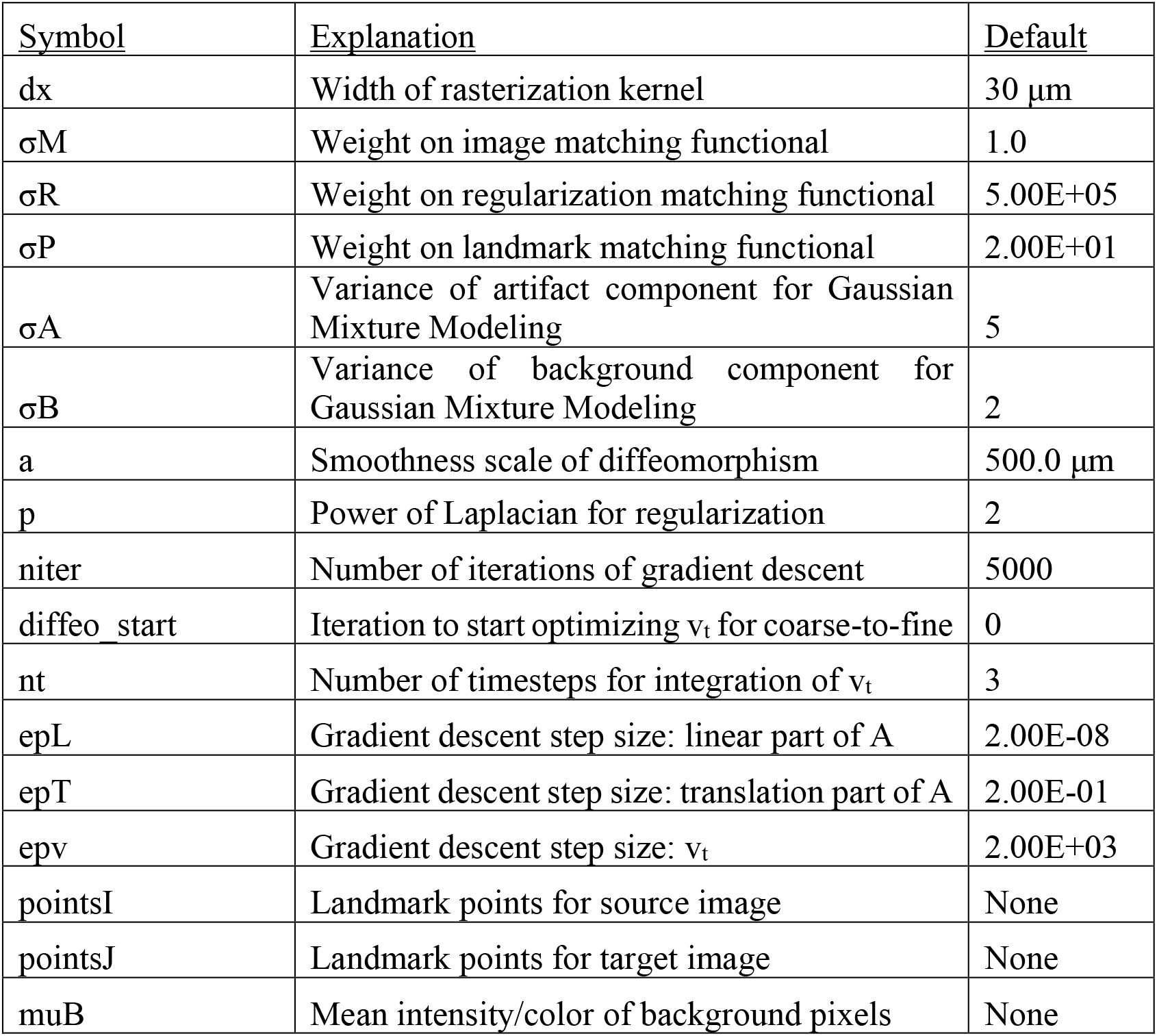
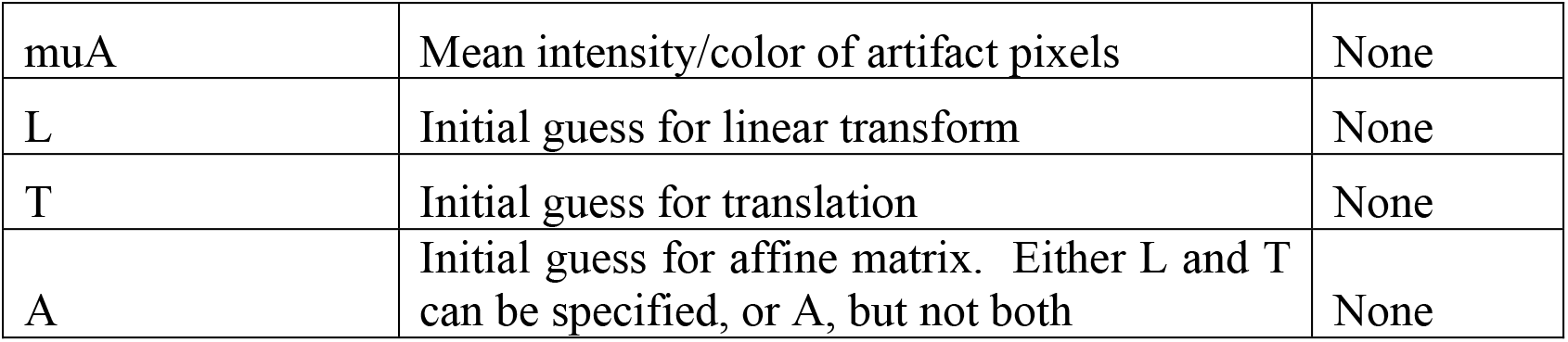

The PyTorch framework was used for automatic gradient calculations. Based on the PyTorch backend, STalign supports parallelization across multiple cores or on GPUs. Derivatives (covectors) are converted to gradient vectors^5,31^ for natural gradient descent^32^.

For improved robustness, Stalign allows users to input pairs of corresponding points in the source and target images. These points can be used to initialize the affine transformation *A* through least squares to steer our gradient based solution toward an appropriate local minimum in this challenging nonconvex optimization problem as well as be added to the objective function to drive the optimization problem itself. Landmark based optimization in the LDDMM framework has been studied extensively^33^. A script point_annotator.py is provided to assist with interactive placement of these points.

#### Runtime Estimate

Runtime for the Stalign.LDDMM function was estimated for CPU settings using a MacBook Pro with an 2.4 GHz 8-Core Intel Core i9 processor and 32 GB 2400 MHz DDR4 memory, and for GPU settings using an Intel Xeon W-3365 2.7GHz Thirty-Two Core 48MB 270W processor with 8 x DDR4-3200 16GB ECC Reg memory and a NVIDIA RTX A5000 24GB PCI-E video card.

## Supplementary Tables

**Supplementary Table 1.**
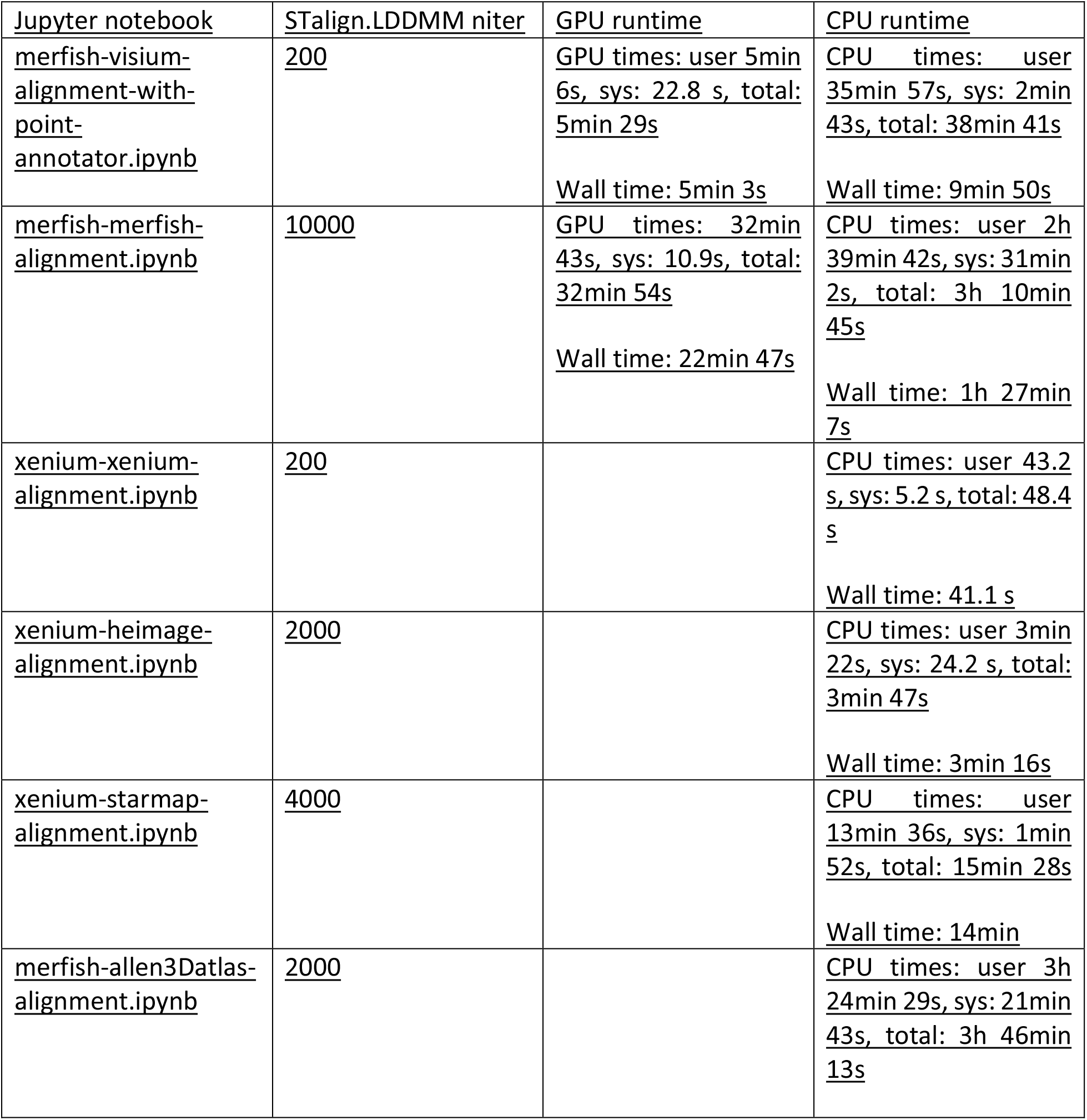
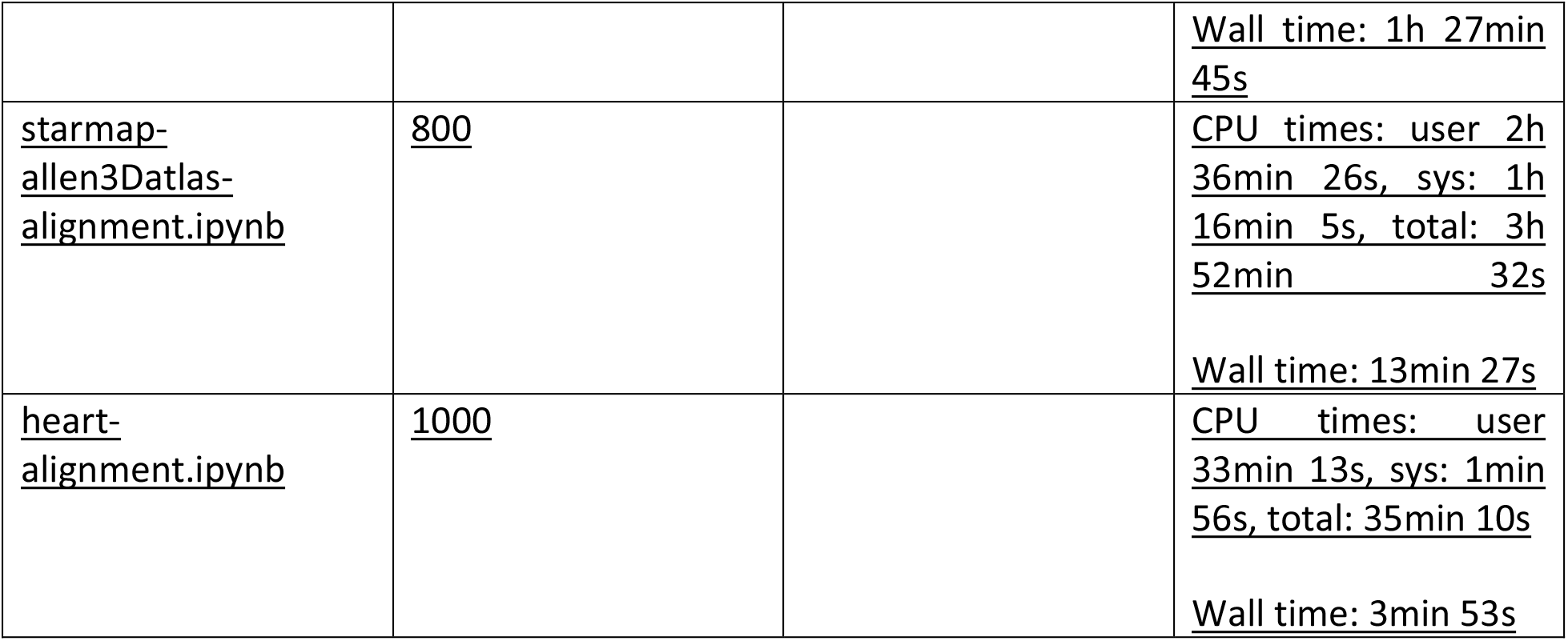
Description of manually placed landmark locations on ST datasets of the mouse brain. (.xlsx)

| <u>Jupyter notebook</u> | <u>STalign.LDDMM niter</u> | <u>GPU runtime</u> | <u>CPU runtime</u> |
| --- | --- | --- | --- |
| <u>merfish-visium-alignment-with-point-annotator.ipynb</u> | <u>200</u> | GPU times: user 5min 6s, sys: 22.8 s, total: 5min 29s<br>Wall time: 5min 3s | CPU times: user 35min 57s, sys: 2min 43s, total: 38min 41s<br>Wall time: 9min 50s |
| <u>merfish-merfish-alignment.ipynb</u> | <u>10000</u> | GPU times: 32min 43s, sys: 10.9s, total: 32min 54s<br>Wall time: 22min 47s | CPU times: user 2h 39min 42s, sys: 31min 2s, total: 3h 10min 45s<br>Wall time: 1h 27min 7s |
| <u>xenium-xenium-alignment.ipynb</u> | <u>200</u> |  | CPU times: user 43.2 s, sys: 5.2 s, total: 48.4 s<br>Wall time: 41.1 s |
| <u>xenium-heimage-alignment.ipynb</u> | <u>2000</u> |  | CPU times: user 3min 22s, sys: 24.2 s, total: 3min 47s<br>Wall time: 3min 16s |
| <u>xenium-starmap-alignment.ipynb</u> | <u>4000</u> |  | CPU times: user 13min 36s, sys: 1min 52s, total: 15min 28s<br>Wall time: 14min |
| <u>merfish-allen3Datlas-alignment.ipynb</u> | <u>2000</u> |  | CPU times: user 3h 24min 29s, sys: 21min 43s, total: 3h 46min 13s |
|  |  |  | Wall time: 1h 27min<br>45s |
| <u>starmap-</u><br><u>allen3Datlas-</u><br><u>alignment.ipynb</u> | <u>800</u> |  | CPU times: user 2h<br>36min 26s, sys: 1h<br>16min 5s, total: 3h<br>52min 32s<br>Wall time: 13min 27s |
| <u>heart-</u><br><u>alignment.ipynb</u> | <u>1000</u> |  | CPU times: user<br>33min 13s, sys: 1min<br>56s, total: 35min 10s<br>Wall time: 3min 53s |

**Supplementary Table 2.** Runtime estimates for the STalign.LDDMM and the STalign.LDDMM_3D_to_slice functions for different ST data alignments.

## Supplementary Figures

**Supplementary Figure 1.**
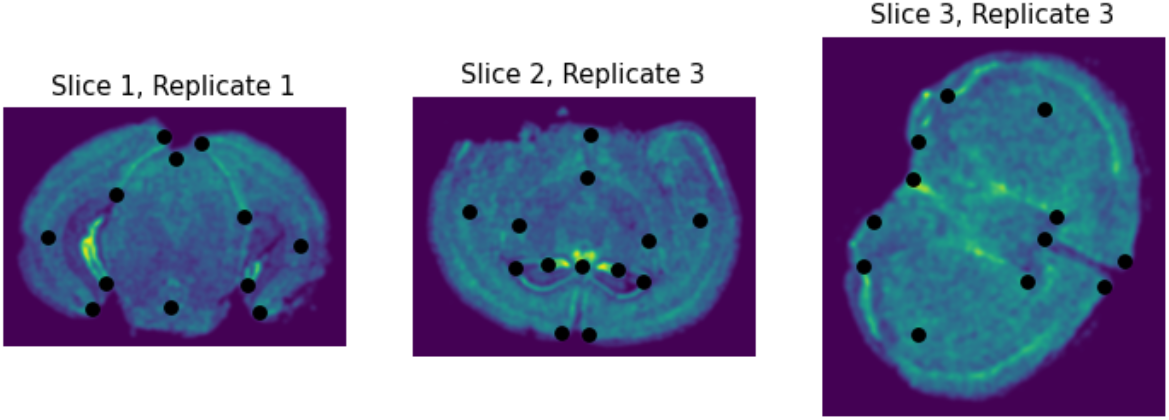
Manually placed landmark locations on ST datasets for one representative biological replicate spanning 3 different locations with respect to bregma.

**Supplementary Figure 2.**
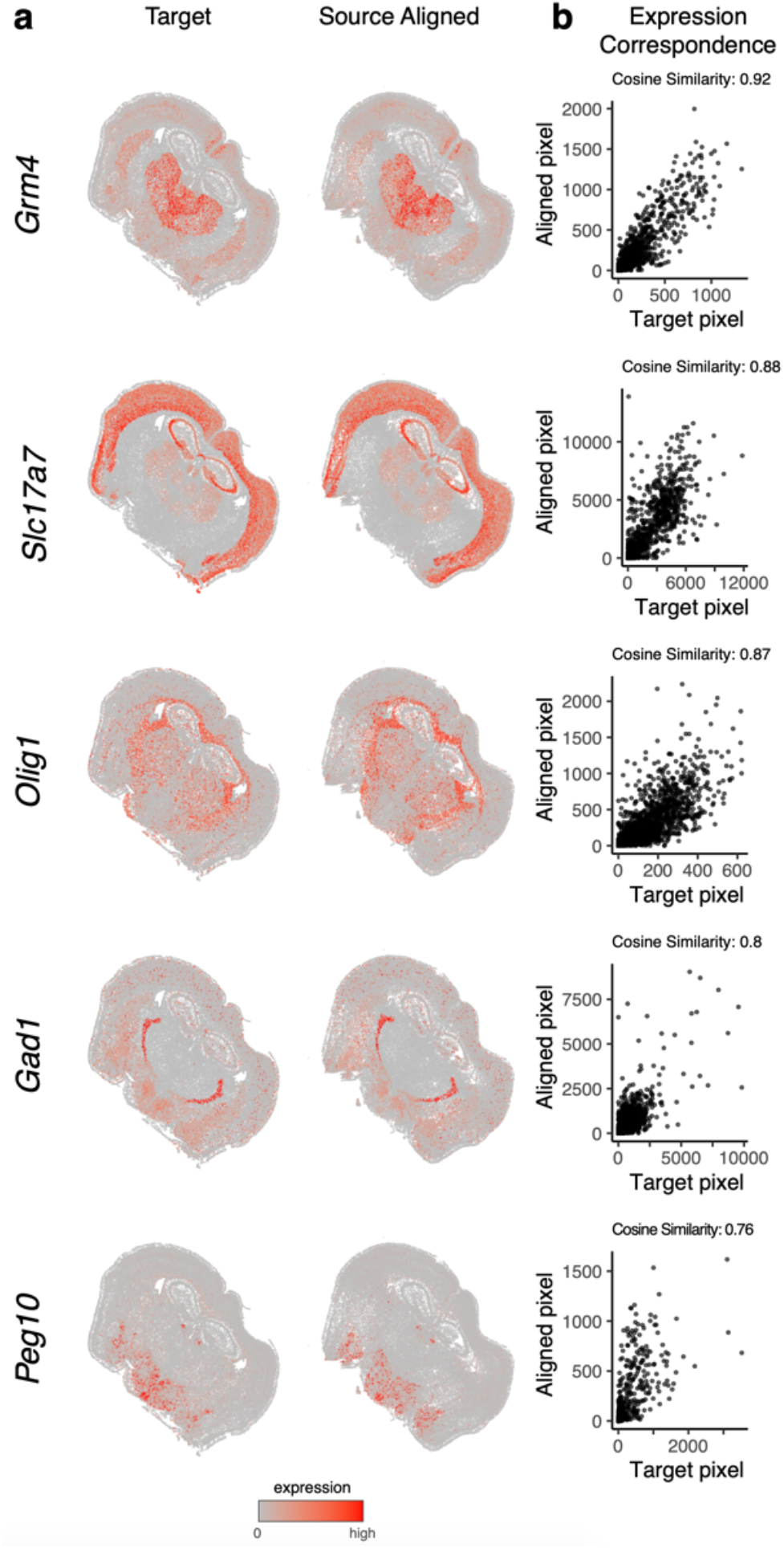
Additional examples of MERFISH to MERFISH alignment for spatially patterned genes. **a**. Correspondence of gene expression spatial organization between the target and aligned source for select spatially patterned genes. **b**. Transcript counts in the target compared to the aligned source at matched pixels for select genes with cosine similarities between transcript counts in target versus aligned source marked.

**Supplementary Figure 3.**
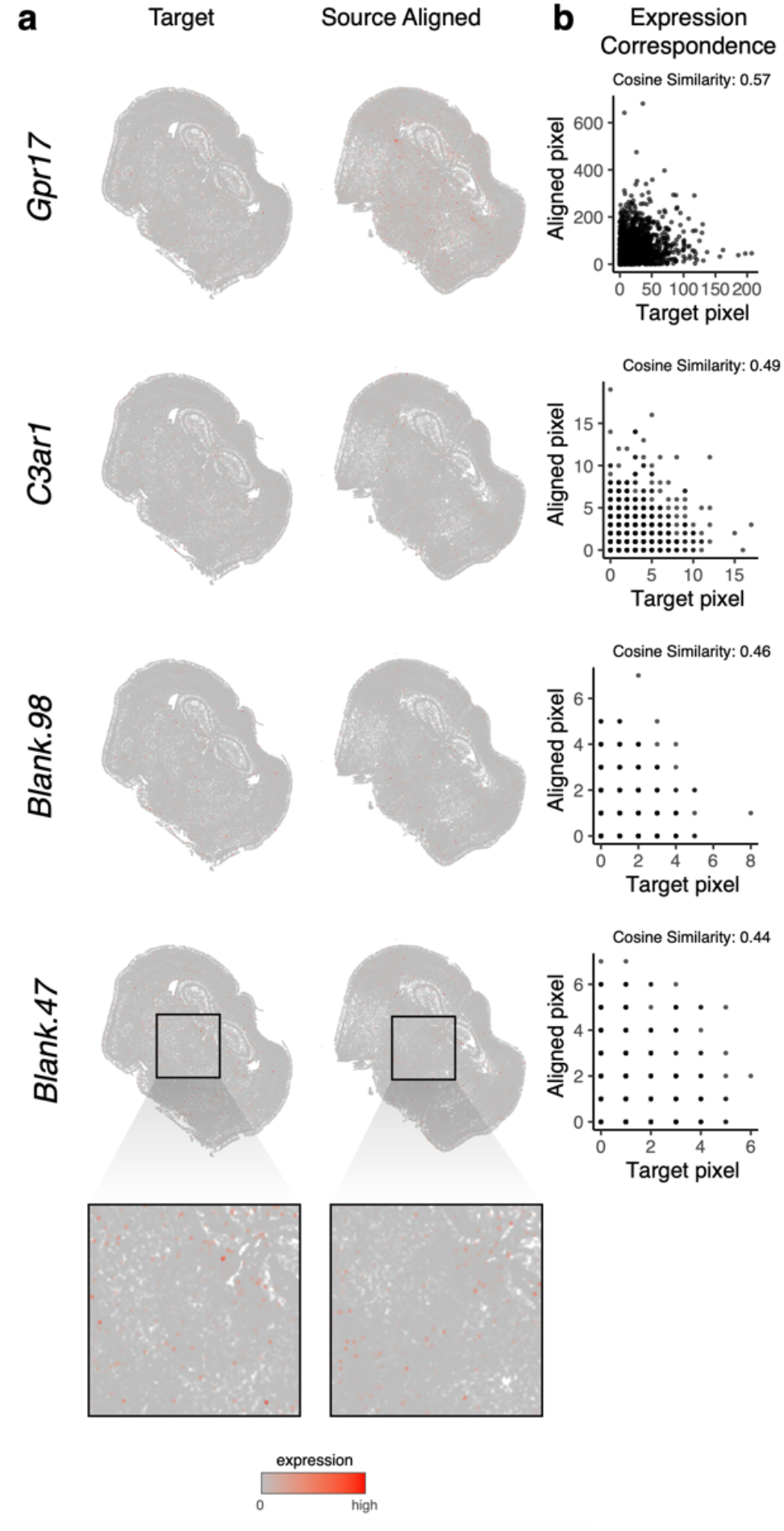
Additional examples of MERFISH to MERFISH alignment for non-spatially patterned genes. **a**. Correspondence of gene expression spatial organization between the target and aligned source for select non-spatially patterned genes (inset displays cells at higher magnification). **b**. Transcript counts in the target compared to the aligned source at matched pixels for select genes with cosine similarities between transcript counts in target versus aligned source marked.

**Supplementary Figure 4.**
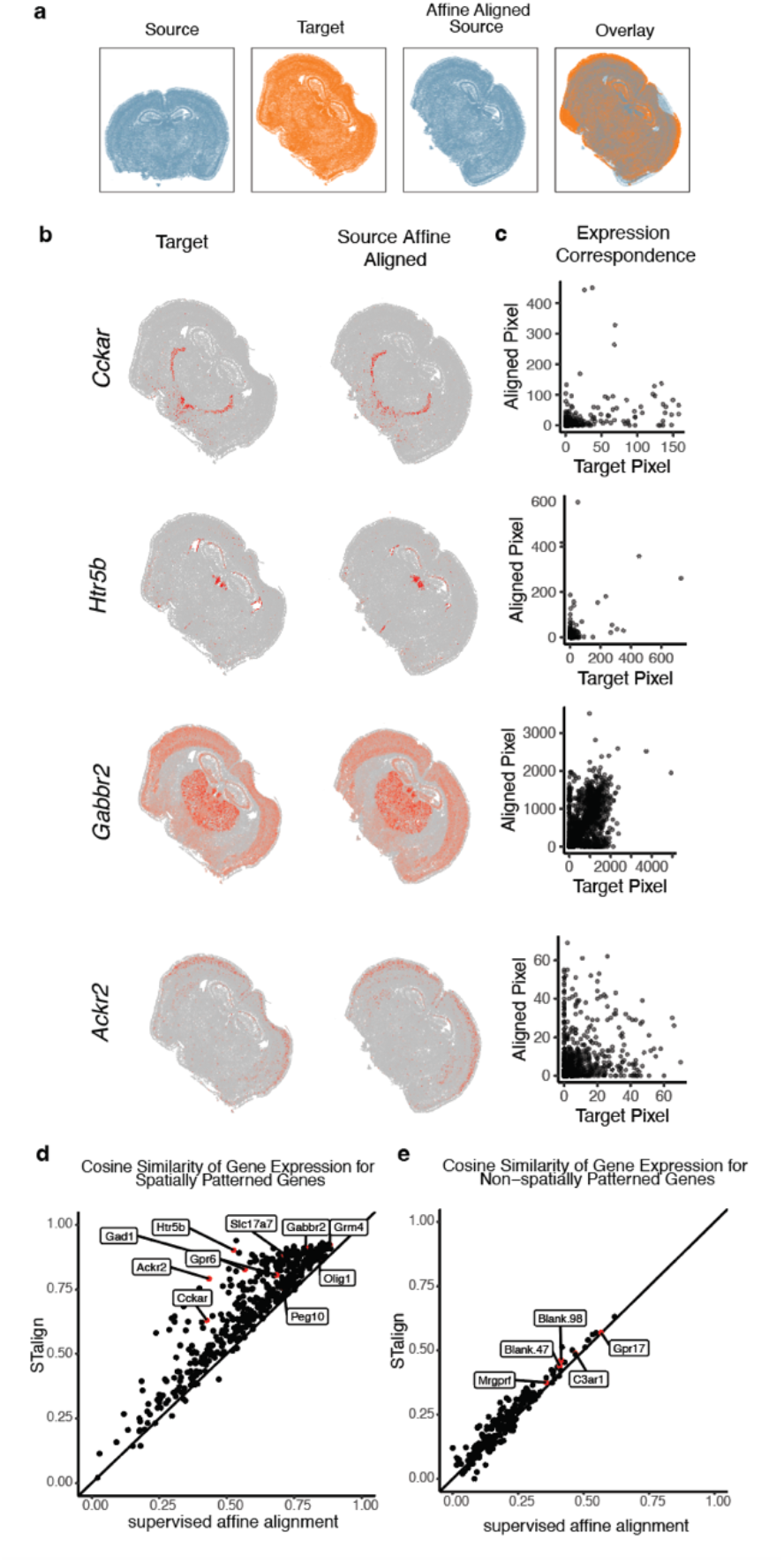
Evaluation of STalign against supervised affine alignment. **a**. Spatial agreement of target and source that has been aligned based on a simple affine transformation based on manually placed landmarks. **b**. Correspondence of gene expression spatial organization between the target and supervised affine aligned source for select spatially patterned genes. **c**. Transcript counts in the target compared to the supervised affine aligned source at matched pixels for select genes: *Cckar, Htr5b, Gabbr2* and *Ackr2*. **d**. Cosine similarities between transcript counts in target versus aligned source for STalign compared to affine alignment for 457 spatially patterned genes. (mean difference = 0.09) Genes featured in Supplemental Figure 4b-c, Figure 2a-c, and Supplemental Figure 2 are highlighted. **e**. Cosine similarities between transcript counts in target versus aligned source for STalign compared to affine alignment for 192 non-spatially patterned genes. (mean difference = 0.02) Genes featured in Figure 2d-f and Supplemental Figure 3 are highlighted.

**Supplementary Figure 5.**
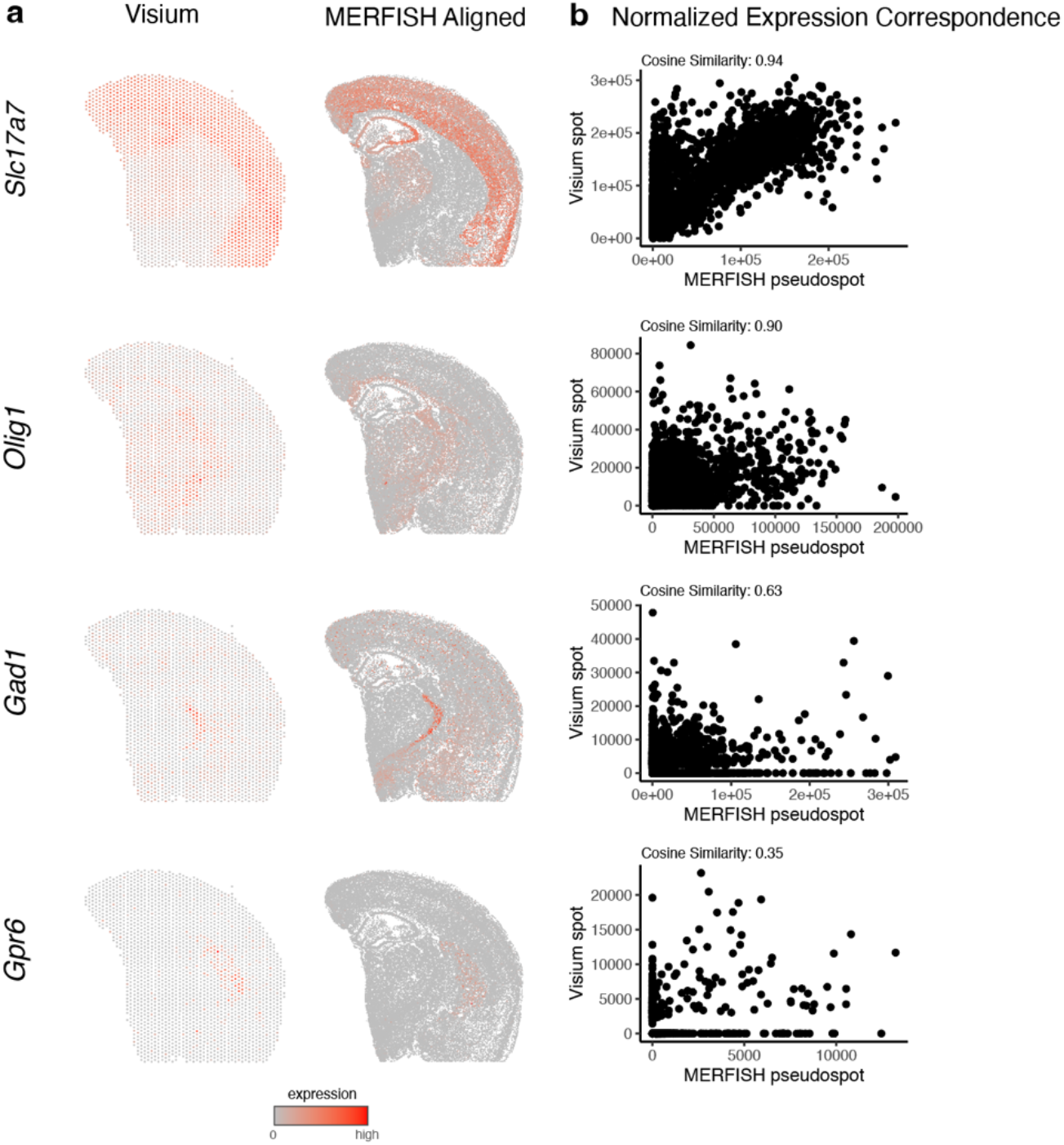
Additional examples of MERFISH to Visium alignment for spatially patterned genes. **a**. Correspondence of gene expression spatial organization between the target and aligned source for select spatially patterned genes. **b**. Normalized gene expression in the Visium target compared to the aligned MERFISH source at matched spots and pseudospots for select genes with cosine similarities between transcript counts in target versus aligned source marked.

**Supplementary Figure 6.**
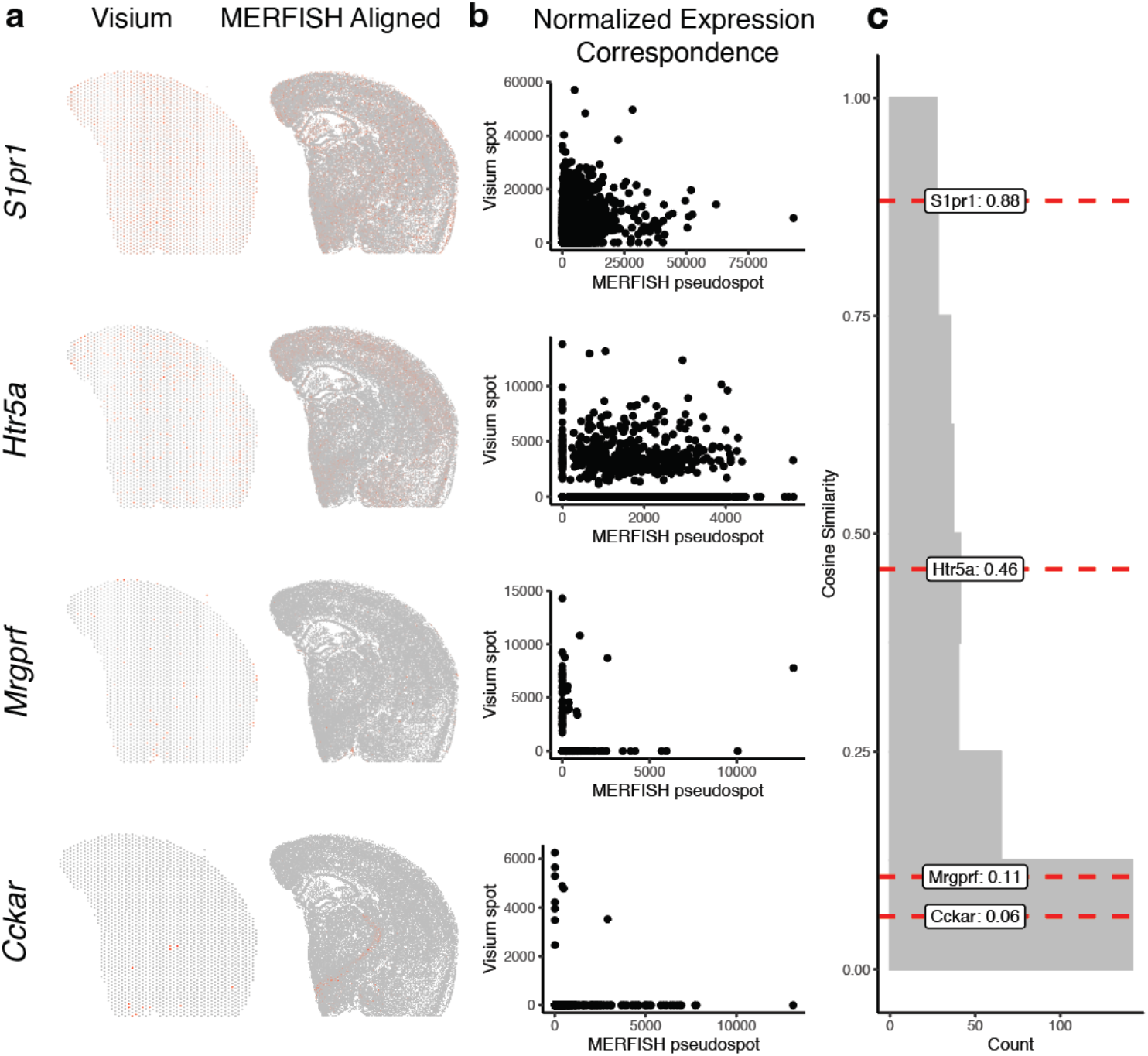
Examples of MERFISH to Visium alignment for spatially non-patterned genes. **a**. Correspondence of gene expression spatial organization between the target and aligned source for select non-spatially patterned genes. **b**. Normalized gene expression in the Visium target compared to the aligned MERFISH source at matched spots and pseudospots for select non spatially patterned genes. **c**. Distribution of cosine similarities between normalized gene expression in the Visium target versus aligned MERFISH source at matched spots and pseudospots for 188 non-spatially patterned genes detected by both ST technologies with select genes marked.

**Supplementary Figure 7.**
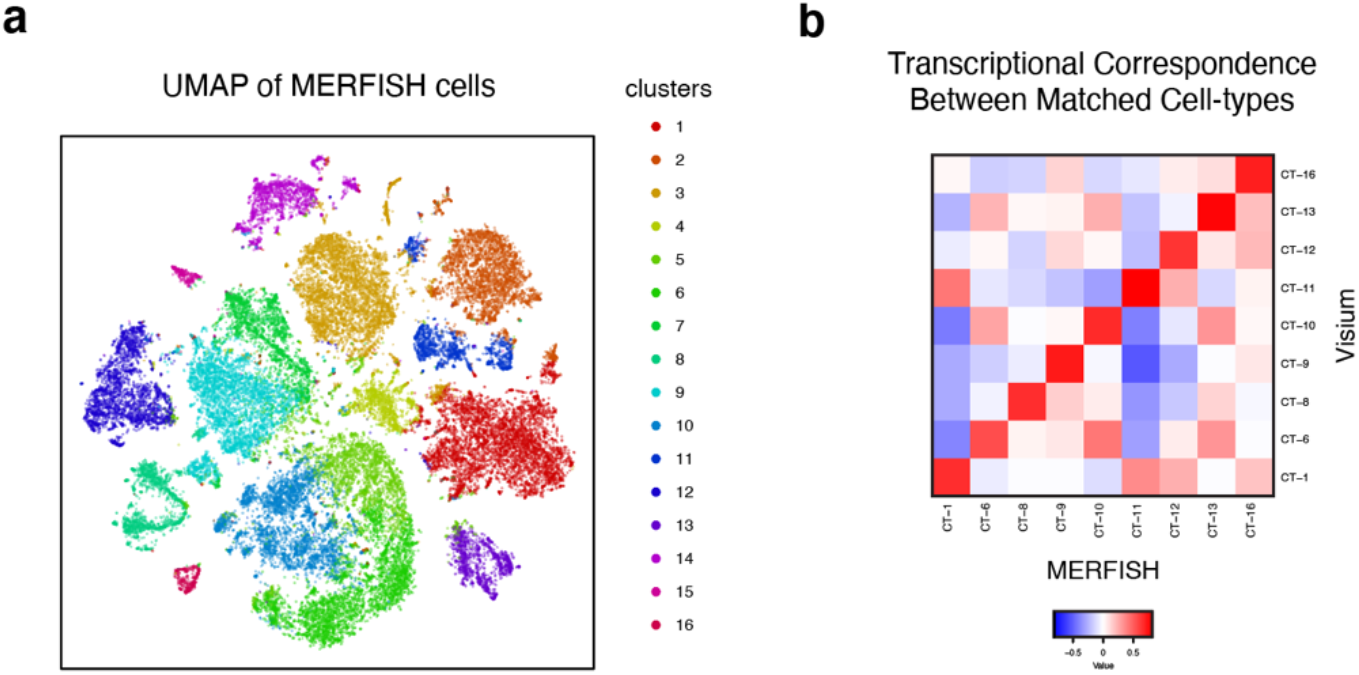
Cell-type correspondence between clustering of MERFISH data and deconvolution of Visium data. **a**. UMAP embedding of MERFISH cells colored by cluster. **b**. Heatmap of transcriptional correlation between the average expression for MERFISH clusters and deconvolved expression for Visium cell-types from STdeconvolve.

**Supplementary Figure 8.**
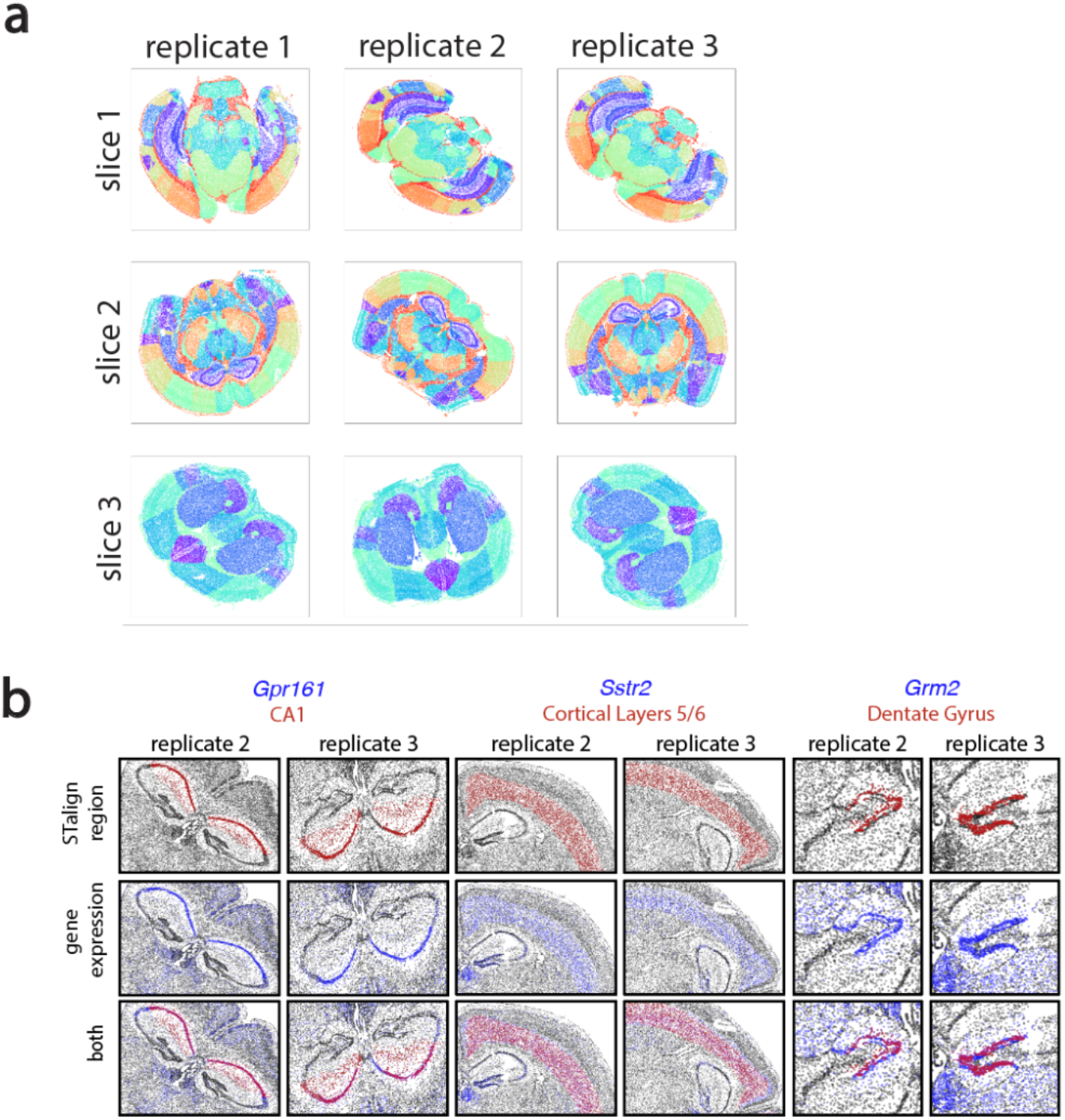
STalign-annotated brain regions. **a**. Brain regions annotated by STalign, represented by different colors, for three biological replicates of three brain slices. **b**. Examples of genes (blue) expressed in brain regions (red) obtained through 3D alignment of MERFISH slices using STalign. Based on gene expression, brain region annotations show consistency and accuracy across replicates of brain slices.

**Supplementary Figure 9.**
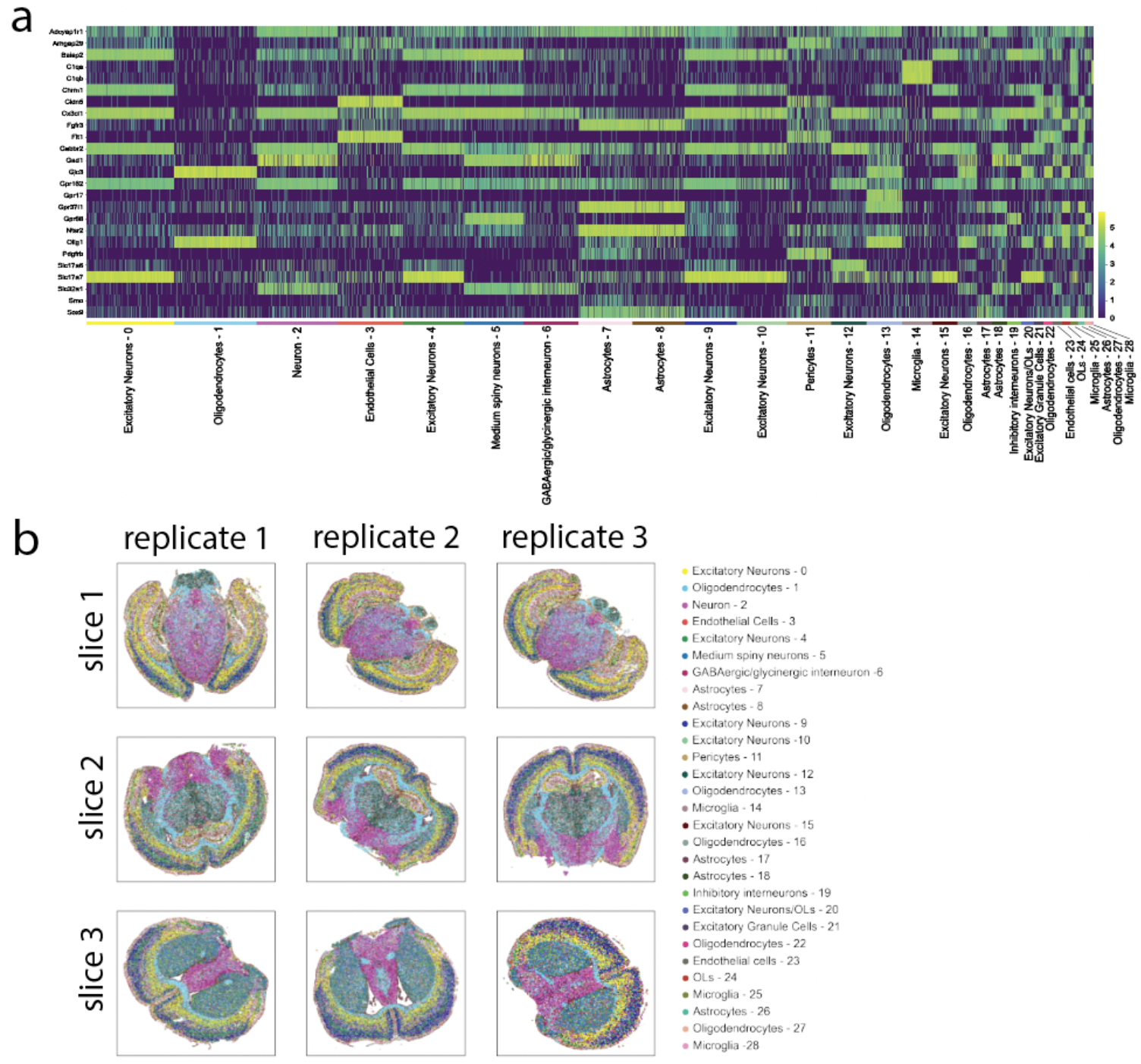
Cell types in MERFISH dataset. **a**. Heat map of differentially expressed genes in cell types defined by Leiden clustering for all cells across 9 MERFISH datasets. **b**. Cell types defined by differential gene expression and Leiden clustering, represented by different colors, for three biological replicates of three brain slices.

**Supplementary Figure 10.**
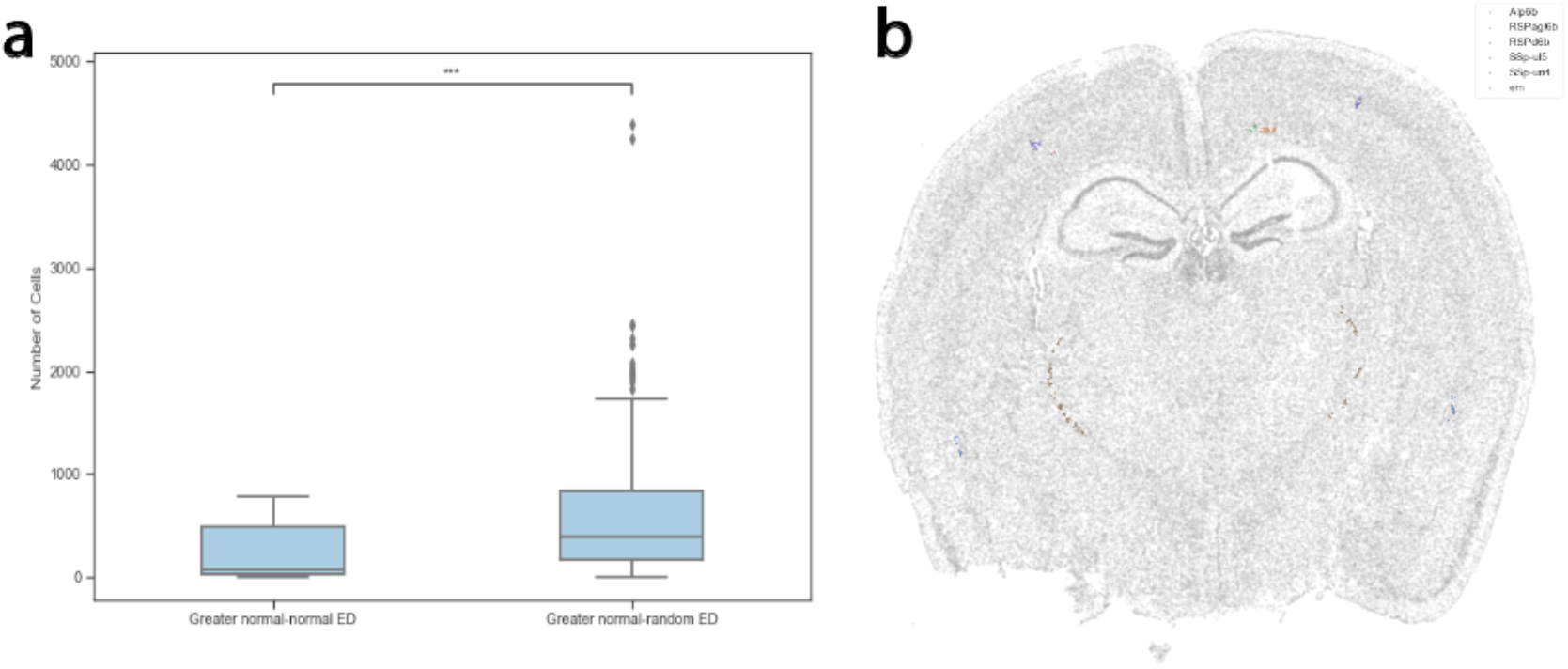
Analyzing brain regions with lower cell-type compositional similarity between replicates compared to size-matched random regions. **a**. Distribution of number of cells in brain regions for which Euclidean-distance (ED) was greater (left) or smaller (right) between replicates compared to matched randomly demarcated brain regions (center line, median; box limits, upper and lower quartiles; whiskers, 1.5x interquartile range; points, outliers) **b**. Representative MERFISH dataset (Slice 2 Replicate 3) of brain regions in the ‘Greater normal-normal ED’ suggestive of lower cell-type compositional similarity between replicates compared to size-matched random regions from (a) which were under 50μm in at least one dimension.

**Supplementary Figure 11.**
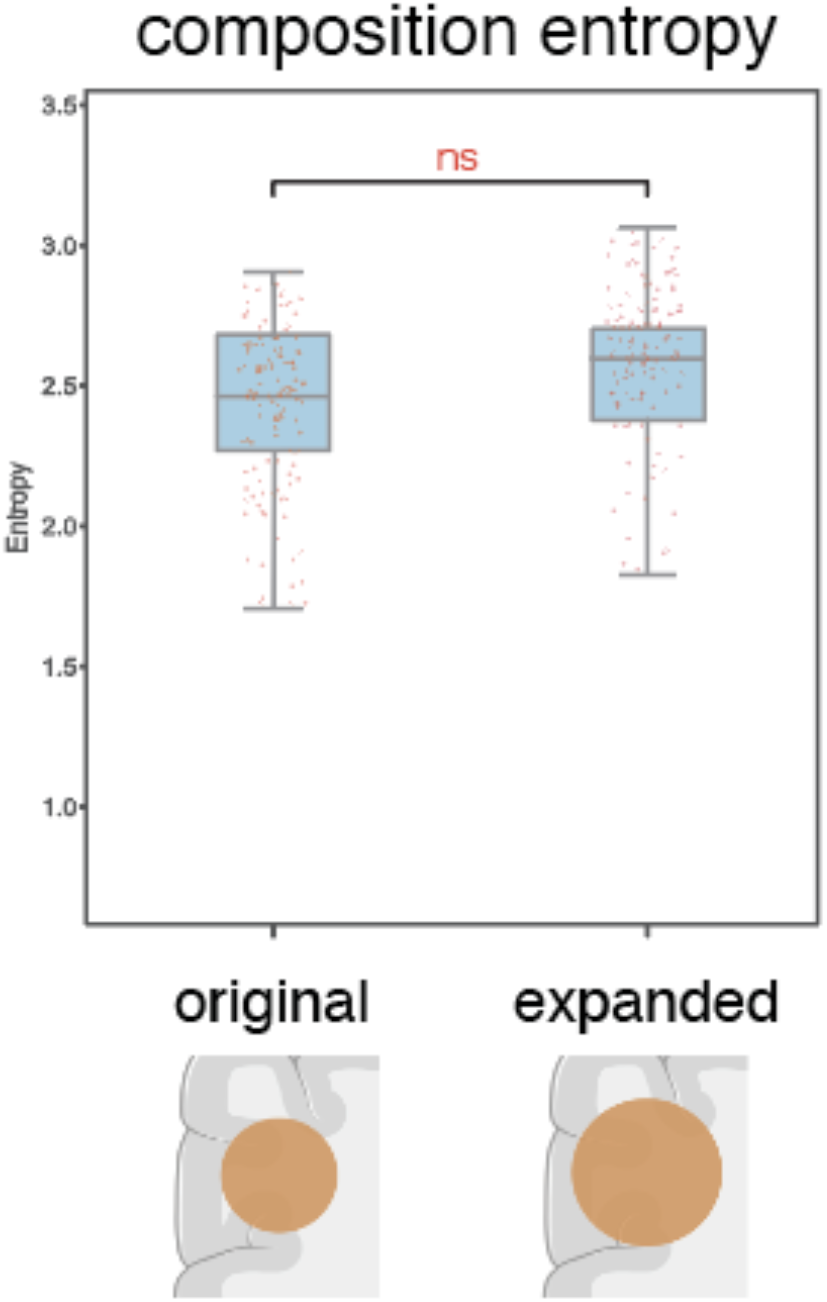
Entropy for size-matched and expanded random brain regions. Non-significant (ns) difference between distribution of cell-type composition entropy for randomly demarcated brain regions that were matched in size for STalign-annotated brain regions (left) versus regions expanded by 100 nearest neighbors (∼100μm) (center line, median; box limits, upper and lower quartiles; whiskers, 1.5x interquartile range; all data points shown)

## 1 Methods

### 1.1 Cell density data model

For single-cell resolution spatially resolved transcriptomics data, we model the point sets of detected cells in the framework of varifold measures [1]. While the theory extends to more complex spaces of features, here we focus on image varifolds [2] by utilizing the locations of cells only, termed the *marginal space measure* (after marginalizing out features other than spatial location) as defined in [3].

Briefly, these space measures are weighted sums of Dirac *δ* distributions 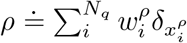 where 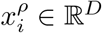 stores the spatial coordinate of the *i*th out of *N*_*q*_ cells, and 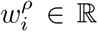 stores its weight. In this work, cell positional data is two dimensional, so *D* = 2, and with some abuse of notation we sometimes write (*x, y*) instead of *x*.

We aim to evaluate the similarity between two single-cell resolution spatially resolved transcriptomics datasets, which we call a source and a target. Note other commonly used terms for source are: template, atlas, or moving image, while another commonly used terms for target is: fixed image. To compute distances between datasets, we embed their corresponding space measures *ρ*_*S*_ and *ρ*_*T*_ respectively in the dual of a Reproducing Kernel Hilbert Space *V* ^*^ and use the standard operator norm (see for example [4]). For some choice of kernel function *k*, the norm squared is

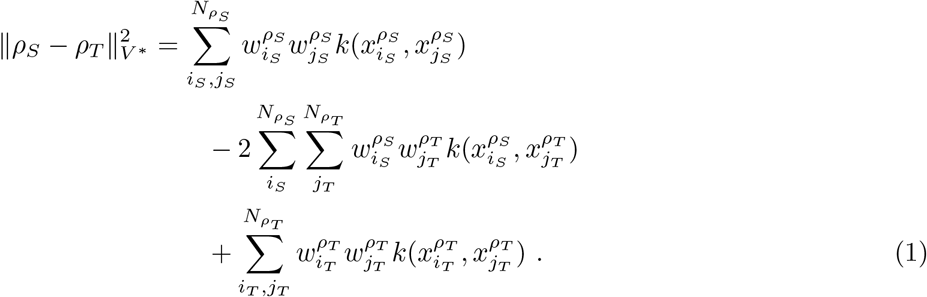

Here we chose *k* as a Gaussian with 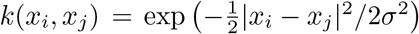, where|·| denotes the standard Euclidean norm, and *σ* is a user specified kernel width parameter.

The variables 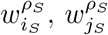, and 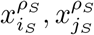 corresponds to the weights and spatial coordinates of the *i*th and *j*th cells in the source, while 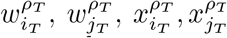 correspond to the weights and spatial coordinates of the cells in the target. For simplicity, in the main paper we write (*x*^*ρ S*^, *y*^*ρ S*^) and (*x*^*ρ T*^, *y*^*ρ T*^) for source and target points respectively.

In STalign, we initialize weights to 1, though applying nonlinear deformations will modify these weights as discussed below in section 1.4.

### 1.2 Rasterization

Since computation of this norm is of quadratic complexity in the number of points, we turn to a more efficient representation for computing optimal transformations through rasterization. We can reduce the complexity of our calculations significantly by approximating our space measures through sampling a density signal on a regular grid (known as rasterization), rather than keeping a list of points and weights.

By defining 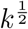 such that 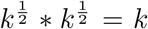 (where* refers to convolution), the above expression for norm squared (1) can be written as

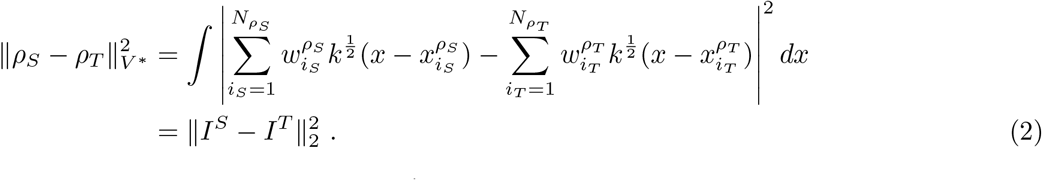

Note that when *k* is a radially symmetric Gaussian, 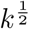 is also a radially symmetric Gaussian but with half the variance. Here we have defined the smooth density function

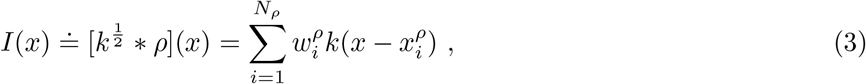

and ∥ · ∥_2_ is the *L*_2_ norm on functions.

Due to the smoothness introduced by 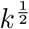, these functions can be accurately discretized by sampling them on a uniform pixel grid at a resolution rate defined by the user and comparable in size to *σ*.

Without rasterization, evaluating the function *I* at a given point would involve a sum over every *x*_*i*_, an order *N* complexity operation. After rasterization the function *I* can be evaluated at any point in order 1 complexity using bilinear interpolation. This allows the norm to be evaluated by summing over a pixel grid in order *P* complexity (where *P* is the number of pixels), rather than a double sum over the points *x*_*i*_ in order *N* ^2^ complexity.

For example, MERFISH Slice 2 Replicate 3 has 85958 cells, and the rasterized dataset has 336*×*256 = 86, 016 pixels. The Naive approach would involve 7,388,777,764 terms in the sum (pairs of cells), whereas in the rasterization approach there are only 86,016 terms in the sum (pixels). This is an approximately 86,000 times increase in efficiency which occurs for each iteration of optimization, ignoring the negligible time cost of rasterization itself, which occurs only once at the start of registration.

In this section we showed how a rasterized image *I* can be produced from a list of cell location, in a manner compatible with the theory of varifolds. However, our registration algorithm can be performed with any standard rasterized image type. For example, in the main manuscript we show examples where *I*^*T*^ is a red-green-blue image corresponding to a brightfield microscopy image of H&E stained tissue. How such images of different contrast profiles are handled is described in section 1.5.

### 1.3 Diffeomorphic transformation model

We estimate alignments between two rasterized datasets by applying a transformation *ϕ* : ℝ^*D*^ → ℝ^*D*^· *ϕ*(*x*) ≐ *Aφ*_1_(*x*), the composition of two transformations: a diffeomorphism (a smooth differentiable transformation with a smooth differentiable inverse) *φ*_1_ : ℝ^*D*^→ ℝ^*D*^ generated in the Large Deformation Diffeomorphic Metric Mapping (LDDMM) framework [5], and an affine transformation *A* (i.e. a 3×3 matrix in homogeneous coordinates whose upper left 2x2 block is a linear transform and upper right 2×1 block is a translation vector). In this notation *Aφ*_1_(*x*) denotes matrix multiplication of the matrix *A* and the vector *φ*_1_ in homogeneous coordinates.

In the LDDMM framework a diffeomormphism is generated by integrating a time varying velocity field *v*_*t*_ over time *t* ∈ [0, 1], by solving the ordinary differential equation

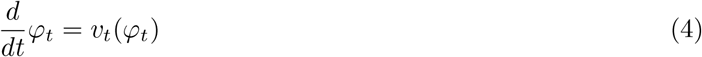

with initial condition *φ*_0_ = id. For identifying alignments, we optimize over *v*_*t*_ rather than *φ*_1_ directly, and to emphasize this dependence we use the superscript *φ*^*v*^ in the main text. Similarly, we use *ϕ*^*A,v*^ to emphasize the dependence of *ϕ* on both *A* and *v*_*t*_. As long as *v*_*t*_ a smooth function of space, 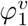is guaranteed to be diffeomorphic. We enforce this through regularization as described below in section 1.5.

While this section described how we parameterize our transformations, next we need to describe how they act to deform our datasets, in order to use them in an optimization problem.

### 1.4 Action of transformations on datasets

The action of a transformation *ϕ* on a space measure dataset *ρ* moves the spatial coordinate of each cell, and adjusts the weight of each cell based on the transformation’s Jacobian determinant.

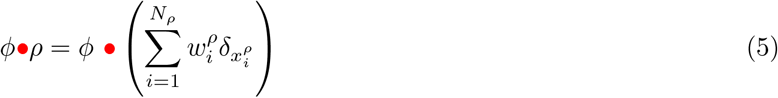

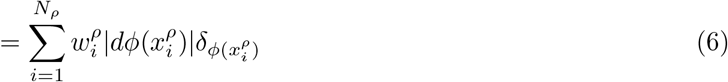

where *dϕ*(*x*) denotes the matrix of partial derivatives of the map *ϕ* at the point *x*, and | · |represents the determinant of a matrix.

We note that the standard image action [*ϕ·I*](*x*) = *I*(*ϕ*^−1^(*x*)) has been well studied theoretically (as a left group action), computationally (in terms of its efficient implementation through interpolation), and application-wise (in terms of its use in a variety of image registration platforms e.g. [5]). This image action for continuous image functions is not appropriate for space datasets and therefore the image action does not match the measure action defined in (5). However, in the dense tissue limit, the continuous image action is consistent with the measure action of (5) as proven in [3]. Since the applications shown here provide a dense approximation, this aforementioned consistency motivates us to leverage the continuous image action for its computational advantages. We write the image action as follows:

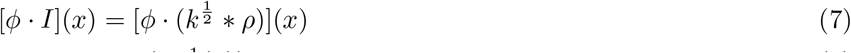

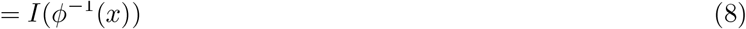

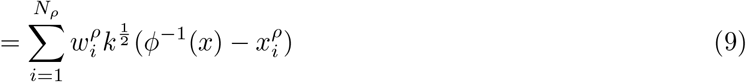

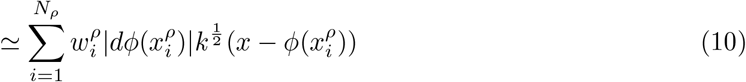

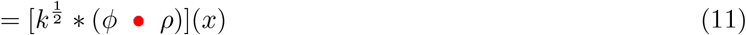

**Online Methods Figure 1:**
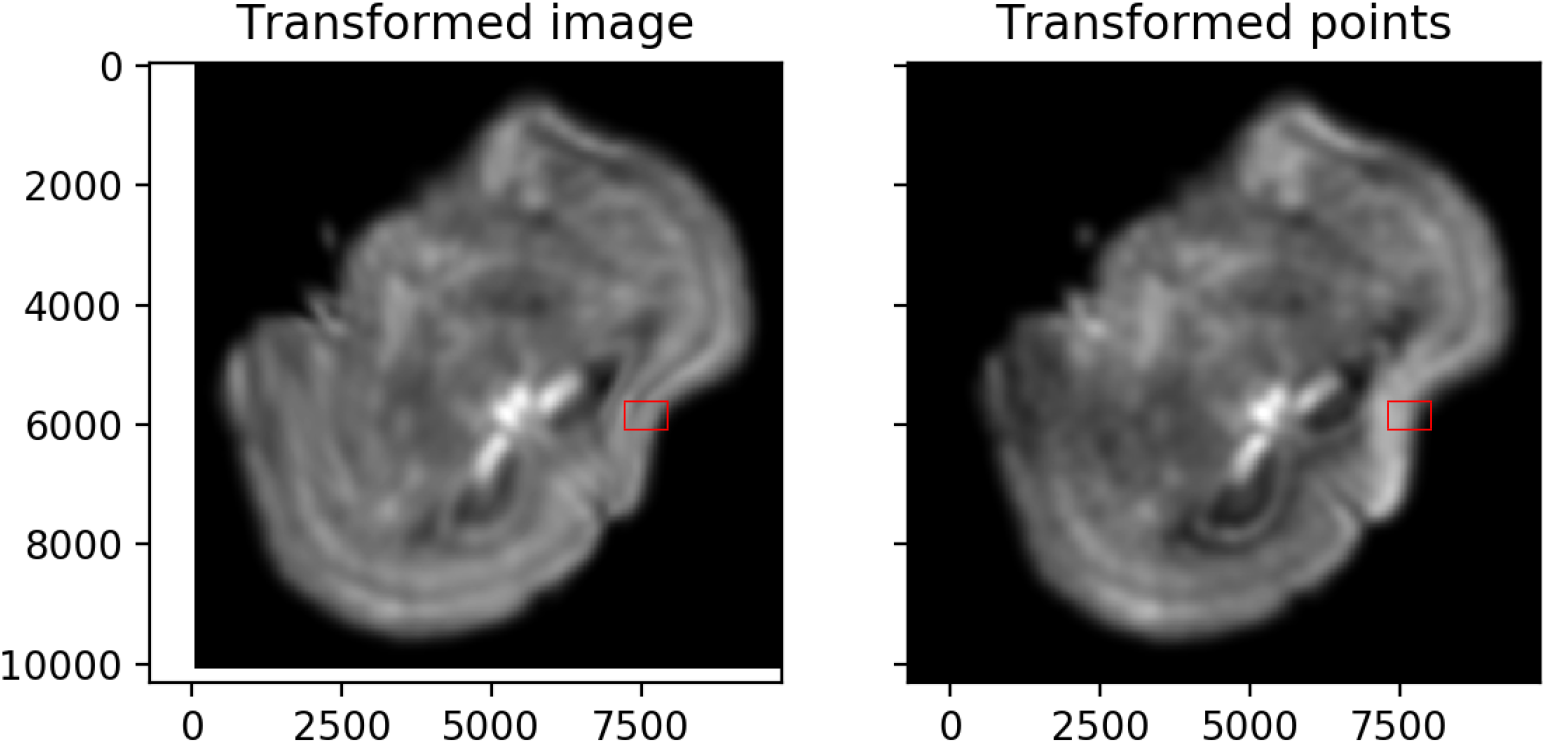
Rasterization followed by deformation with the image action (left), versus naively deforming the positions of cells followed by rasterization (right). Note the intensity changes that occur in regions of high deformation.

The approximate equality is accurate in our examples when 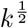 is narrow relative to the smoothness of *v*_*t*_ and wide relative to the spacing between cells. The approximate equality would be exact if 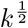 were a Dirac delta function. If cells are too far apart, a larger value of *σ* could be chosen. For a particularly sparse set of cells, a different method that does not include rasterization would be more appropriate, for example measure matching [6].

With this formulation, deformations can be applied to smooth density images *I* using interpolation in order *P* (number of pixels) complexity. Online Methods Figure 1 illustrates the importance of the Jacobian factor. Without including this factor, transforming a density image alters its brightness, which is typically undesirable: a bigger organ tends to have more cells with the same cell density, rather than the same number of cells with a lower density.

### 1.5 Image registration

We compute a spatial alignment between two ST datasets by minimizing the sum of two objective functions: a regularization term *R*, and a matching term *M*_*θ*_,

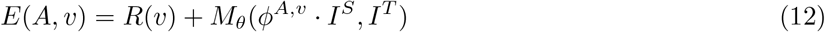

which we define below. Note that the computation of *I* is described in section 1.2, the parameterization of *ϕ*^*A,v*^ is described in 1.3, and the action *ϕ*^*A,v*^ *I*^*S*^ is described in the section 1.4. Recall that *ϕ* depends on both the velocity field *v* and the affine transform *A*.

Following the LDDMM framework, we regularize our diffeomorphism via

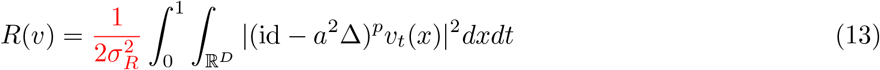

where id is an identity matrix, 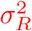 is a user tunable parameter that adjusts balance between matching accuracy and regularization, where large values correspond to less regularization and higher matching accuracy, and small values correspond to more regularization and lower matching accuracy. Δ is the Laplacian, *a* is a constant with units of length that controls spatial smoothness, and *p* = 2 is a power that must be large enough to guarantee that results are diffeomorphisms [7]. Note that small values of *a* may be overfitting of noise whereas large values of *a* may lead to low accuracy. In practice, we chose a value of *a* based on the spatial smoothness of deformations that we believe to be realistic. We then consider several values of 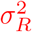 (starting with a value provided by one of our online examples), and chose the one that achieves a reasonable balance between regularization and accuracy.

Our matching takes the form of [8]

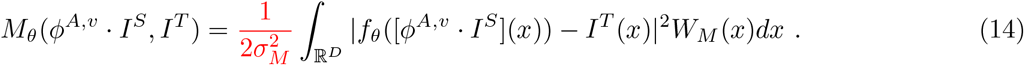

where 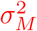 is a user tunable parameter that describes the amount of noise in our imaging data (see description of Gaussian Mixture modeling below).

Note that *I*^*T*^ need not correspond to a smooth density image as defined in section 1.2. For example, we include the case where it is a red green blue image corresponding to an H&E stain.

The function *f*_*θ*_ is a transformation of image contrast with unknown parameters *θ*. We use a polynomial for *f*_*θ*_, in which case the minimizing parameters *θ* can be found exactly by solving a weighted least squares problem. The purpose of this transformation is to model differences in contrast between images from the same modality due to calibration issues; and contrast/color differences between different modalities. In this work we found that first order polynomials were sufficient for accurate image registration. In other work in neuroimaging we have used 3rd order polynomials, which have enough degrees of freedom to map the intensity of gray matter, white matter, and background to arbitrary intensities [8]. Because there are many more pixels than degrees of freedom, it is unlikely that these polynomials will overfit the observed data *I*^*T*^. However, depending on the initialization of transformation parameters this is possible: if tissue in *I* and *I*^*T*^ do not overlap at all, parameters *θ* may be estimated to zero out imaging information and transform *I* into a constant function that looks like background only.

The term *W* is a positive weight that represents the probability that a given pixel in the target image can be matched accurately to one in the source image. For example, if tissue is missing in the target image but not the source image, pixels in the region of missing tissue would get a small weight. Similarly, if the target image included a signal not present in the source (e.g. a bright fluorescence signal).

To optimize *E*, we alternate between updating *W*_*M*_ with Gaussian mixture modeling, and jointly updating (*θ, ϕ*^*A,v*^) with gradient based methods, using expectation maximization algorithm as discussed in [8]. Briefly, we use 3 classes in our Gaussian mixture model (pixels to be matched, background, and artifact). Each is modeled as a Gaussian random variable with an unknown mean (optionally the mean can be assumed known and specified as an input parameter), a known variance (specified as an input parameter), and an unknown prior probability. While the means for background and artifact are constant values, the mean for pixels to be matched is equal to *f*_*θ*_([*ϕ*^*A,v*^ *I*^*S*^](*x*)) and is a function of space. Parameters are estimated by standard Gaussian mixture modeling techniques, and *W*_*M*_ (*x*) is computed as the posterior probability that the pixel at *x* belongs to the “pixels to be matched” class. If *g*(*x, μ, σ*^2^) is a multivariate normal with mean *μ* and covariance *σ*^2^ times identity, and *π*_*i*_ are prior probabilities for each class (*i* ∈ (matching, background, and artifact)), then

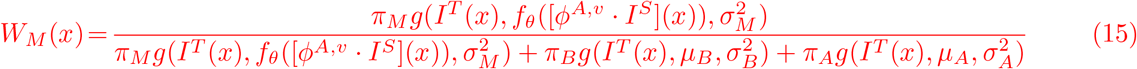

In the above expression, note that the denominator shows a mixture of three Gaussians, and the numerator shows the first class in the mixture. In our code we also define *W*_*B*_ and *W*_*A*_, which are posterior probabilities that a pixel belongs to the background or artifact classes. They are defined with same denominator as *W*_*M*_, but with numerators corresponding to the mixture component for their class. Recall that updating *ϕ* corresponds to updating the affine transformation matrix *A*, and the velocity field *v*_*t*_ which generates the deformation *φ*_1_ from (4).

After solving for the optimal transformation parameters *A* and *v*_*t*_, a transformation and its inverse are constructed by solving (4) sampled on a regular grid, using Semi-Lagrangian techniques [9]. With 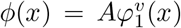 and 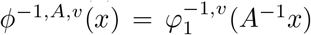 computed, cell locations 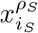 in the source image can be mapped into the target by calculating 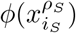 through linear interpolation. Similarly, a point 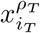 in the target image can be mapped to the atlas by calculating 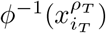 through linear interpolation.

For improved robustness, our software allows users to input pairs of corresponding points in the source and target images. These points can be used either to initialize the affine transformation *A* through least squares (steering our gradient based solution toward an appropriate local minima in this challenging nonconvex optimization problem); or can be used to drive the optimization problem itself by modifying to our objective function to be *E*(*A, v*) = *R*(*v*) + *M*_*θ*_(*ϕ*^*A,v*^ *·I*^*S*^, *I*^*T*^) + *P* (*ϕ*^*A,v*^(*X*^*S*^), *X*^*T*^) such that

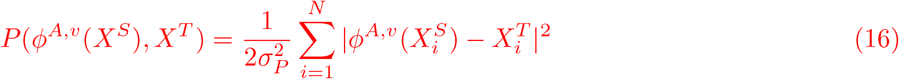

where 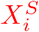 and 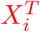 are the *i*th point of N corresponding points in the source and target respectively and 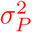 is a user tunable parameter that adjusts balance between matching corresponding landmark points, matching images, and regularization, where large values correspond to less accuracy matching points and small values correspond to more accuracy matching points. Landmark based optimization in the LDDMM framework has been studied extensively (see for example [10]).

#### 1.6 3D to 2D alignment

In addition to aligning spatially resolved transcriptomics datasets in which the cell positional information is 2D, we registered the 3D reconstructed Allen common coordinate framework (CCF) atlas (source) to each of the 9 MERFISH datasets (target). The image transformation is similar to the alignment discussed in the section 1.5 with a few exceptions:

It is important to note that all transformations are performed on the source 3D atlas. Since the 50 *μ*m

Nissl-stained Allen Brain Atlas CCF v3 was used as the source image, rasterization is not applied to the atlas. The affine transformation *A* for the 3D-2D alignment is a 4×4 3D matrix in homogeneous coordinates.

The space of dense 3D images in the orbit of the atlas, are defined via diffeomorphisms

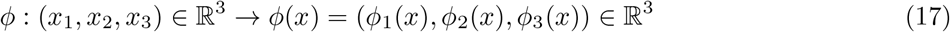

The diffeomorphism *ϕ* ∈ *D* acts on the atlas to generate the orbit of imagery ℐ,

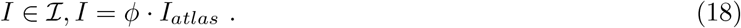

The velocity field *v*_*t*_ is still defined by (4), but *v*_*t*_ ∈ ℝ^3^.

The image *I*^*S*^ in the matching term *M* represents the transformed source atlas evaluated at *z* = 0, to enable comparison in the same dimension between the source and the target images.

