## supplementary software for "Alignment of spatial transcriptomics data using diffeomorphic metric mapping": point_annotator.py

```
''' Point annotator

ex

python point_annotator.py ../visium_data/Merfish_S2_R3.npz ../visium_data/tissue_hires_image.npz
'''

import argparse
from os.path import split,join,splitext
import numpy as np
import matplotlib.pyplot as plt
import matplotlib.transforms as mtransforms
from glob import glob

if __name__ == '__main__':
    print('hello world')

    parser = argparse.ArgumentParser(
        prog='point_annotator',
        description='Takes two images as an input (npz format x I), and provides an interface for annotating points',
    )

    parser.add_argument(
        'filename', nargs=2,
        help='Filename for source image data.  This should be a npz file containing "x,y" and "I"',
    )

    #parser.add_argument(
    #    'filename2',
    #    help='Filename for target image data.  This should be a npz file containing "x,y" and "J"',
    #)

    parser.add_argument(
        'output',nargs='*',
        default=None,
        help='Defaults to source image input name with a suffix "_points.npy"',
    )

    # parser.add_argument(
    #     '-o,','--output1',
    #     default=None,
    #     help='Defaults to source image input name with a suffix "_points.npy"',
    # )
    # parser.add_argument(
    #     '-o,','--output2',
    #     default=None,
    #     help='Defaults to target image input name with a suffix "_points.npy"',
    # )

    args = parser.parse_args()
    print(args)

    # load the source image
    # start with a special case
    if args.filename[0].isnumeric():
        ind = int(args.filename[0])
        files = glob('/home/dtward/bmaproot/nafs/dtward/merfish/jean_fan_2021/OneDrive_1_8-5-2021/*_rasterized.npz')
        files.sort()
        args.filename1 = files[ind]
        print(f'special case, using index {ind} to select file {args.filename1}')
        
    try:
        dataS = np.load(args.filename[0])
    except:
        raise Exception(f'Could not load source input image {args.filename[0]}')
    print(f'source image contains ')
    print([k for k in dataS])

    # load the target image
    # start with a special case
    if args.filename[1].isnumeric():
        ind = int(args.filename[1])
        files = glob('/home/dtward/bmaproot/nafs/dtward/merfish/jean_fan_2021/OneDrive_1_8-5-2021/*_rasterized.npz')
        files.sort()
        args.filename[1] = files[ind]
        print(f'special case, using index {ind} to select file {args.filename[1]}')
        
    try:
        dataT = np.load(args.filename[1])
    except:
        raise Exception(f'Could not load target input image {args.filename[1]}')
    print(f'target image contains ')
    print([k for k in dataT])

    # get the source output name
    if len(args.output) == 0:
        outputS = args.filename[0].replace('.npz','_points.npy')
    print(f'source output name {outputS}')

    # get the target output name
    if len(args.output) == 0:
        outputT = args.filename[1].replace('.npz','_points.npy')
    print(f'target output name {outputT}')

    # let's draw the source image
    xI = dataS['x']
    yI = dataS['y']
    I = dataS['I'].transpose(1,2,0).squeeze()

    dxI = xI[1] - xI[0]
    dyI = yI[1] - yI[0]

    # let's draw the target image
    xJ = dataT['x']
    yJ = dataT['y']
    J = dataT['I'].transpose(1,2,0).squeeze()

    dxJ = xJ[1] - xJ[0]
    dyJ = yJ[1] - yJ[0]

    # make plots bigger
    plt.rcParams["figure.figsize"] = (12,8)
    
    fig,ax = plt.subplots(1,2)
    ax[0].imshow(I,extent=(xI[0]-dxI/2.0,xI[-1]+dxI/2,yI[-1]+dyI/2,yI[0]-dyI/2))
    ax[1].imshow(J,extent=(xJ[0]-dxJ/2.0,xJ[-1]+dxJ/2,yJ[-1]+dyJ/2,yJ[0]-dyJ/2))
    ax[0].set_title('source') 
    ax[1].set_title('target')
    plt.show(block=False)

    # let's draw the source points if output has been created previously 
    dataS = {}
    try:
        dataS = np.load(outputS,allow_pickle=True).item()
    except:

        print(f'could not load previous')


    for k in dataS:
        points = dataS[k]
        trans_offset_0 = mtransforms.offset_copy(ax[0].transData, fig=fig, x=0.05, y=-0.05, units='inches')
        ax[0].scatter([p[0] for p in points],[p[1] for p in points], c='red', s=10)
        for i in range(len(points)):
            ax[0].text(points[i][0], points[i][1],f'{k}{i}', c='red', transform=trans_offset_0)
    plt.show(block=False)

    # let's draw the target points if output has been created previously 
    dataT = {}
    try:
        dataT = np.load(outputT,allow_pickle=True).item()
    except:

        print(f'could not load previous')

    for k in dataT:
        points = dataT[k]
        trans_offset_1 = mtransforms.offset_copy(ax[1].transData, fig=fig, x=0.05, y=-0.05, units='inches')
        ax[1].scatter([p[0] for p in points],[p[1] for p in points], c='red', s=10)
        for i in range(len(points)):
            ax[1].text(points[i][0], points[i][1],f'{k}{i}', c='red', transform=trans_offset_1)
    plt.show(block=False)
    
    # let's choose new points
    count = 0
    while True:
        name = input(f'Enter name for landmark structure {count} (or leave blank and hit enter to save): ')
        if len(name) == 0:
            break       
        fig.suptitle(f'annotate {name} by alternately selecting one point in source and one point in target with left clicks \n then press enter to store all points of the {name} \n (use delete or backspace to remove last point)')        
        #plt.show(block=False)
        plt.pause(0.001)
        points = plt.ginput(-1, timeout=60)
        if len(points) == 0:
            break

        print(points)
        pointsS = points[::2]
        pointsT = points[1::2]
        
        trans_offset_0 = mtransforms.offset_copy(ax[0].transData, fig=fig, x=0.05, y=-0.05, units='inches')
        ax[0].scatter([p[0] for p in pointsS],[p[1] for p in pointsS], c='red', s=10)
        for i in range(len(pointsS)):
            ax[0].text(pointsS[i][0], pointsS[i][1],f'{name}{i}', c='red', transform=trans_offset_0)

        trans_offset_1 = mtransforms.offset_copy(ax[1].transData, fig=fig, x=0.05, y=-0.05, units='inches')
        ax[1].scatter([p[0] for p in pointsT],[p[1] for p in pointsT], c='red', s=10)
        for i in range(len(pointsT)):
            ax[1].text(pointsT[i][0], pointsT[i][1],f'{name}{i}', c='red', transform=trans_offset_1)

        #plt.show(block=False)
        plt.pause(0.001)
        dataS[name] = pointsS
        dataT[name] = pointsT
        count += 1

        


    print(dataS)
    print(dataT)
    np.save(outputS,dataS,)
    np.save(outputT,dataT,)
    
    

    
```
