## supplementary software for "Alignment of spatial transcriptomics data using diffeomorphic metric mapping": STalign.py

```
''' Tools for aligning spatial transcriptomics data
'''

import numpy as np
import matplotlib.pyplot as plt

import torch
from torch.nn.functional import grid_sample

import numpy as np
import matplotlib.pyplot as plt
import pandas as pd # for csv.
from matplotlib import cm
from matplotlib.lines import Line2D
import os
from os.path import exists,split,join,splitext
from os import makedirs
import glob
import requests
from collections import defaultdict
import nrrd
import torch
from torch.nn.functional import grid_sample
import tornado
import sys
import numpy as np
import plotly.graph_objs as go
import plotly.express as px
import copy
import pandas as pd

def normalize(arr, t_min=0, t_max=1):
    """Linearly normalizes an array between two specifed values.
    
    Parameters
    ----------
    arr : numpy array
        array to be normalized
    t_min : int or float
        Lower bound of normalization range
    t_max : int or float
        Upper bound of normalization range
    
    Returns
    -------
    norm_arr : numpy array
        1D array with normalized arr values
        
    """
    
    diff = t_max - t_min
    diff_arr = np.max(arr) - np.min(arr)
    min_ = np.min(arr)
        
    norm_arr = ((arr - min_)/diff_arr * diff) + t_min
    
    return norm_arr


def rasterize(x, y, g=np.ones(1), dx=30.0, blur=1.0, expand=1.1, draw=10000, wavelet_magnitude=False,use_windowing=True):
    ''' Rasterize a spatial transcriptomics dataset into a density image
    
    Paramters
    ---------
    x : numpy array of length N
        x location of cells
    y : numpy array of length N
        y location of cells
    g : numpy array of length N
        RNA count of cells
        If not given, density image is created
    dx : float
        Pixel size to rasterize data (default 30.0, in same units as x and y)
    blur : float or list of floats
        Standard deviation of Gaussian interpolation kernel.  Units are in 
        number of pixels.  Can be aUse a list to do multi scale.
    expand : float
        Factor to expand sampled area beyond cells. Defaults to 1.1.
    draw : int
        If True, draw a figure every draw points return its handle. Defaults to False (0).
    wavelet_magnitude : bool
        If True, take the absolute value of difference between scales for raster images.
        When using this option blur should be sorted from greatest to least.
    
        
    Returns
    -------
    X  : numpy array
        Locations of pixels along the x axis
    Y  : numpy array
        Locations of pixels along the y axis
    M : numpy array
        A rasterized image with len(blur) channels along the first axis
    fig : matplotlib figure handle
        If draw=True, returns a figure handle to the drawn figure.
        
    Raises
    ------    
    Exception 
        If wavelet_magnitude is set to true but blur is not sorted from greatest to least.
        
        
    
    Examples
    --------
    Rasterize a dataset at 30 micron pixel size, with three kernels.
    
    >>> X,Y,M,fig = tools.rasterize(x,y,dx=30.0,blur=[2.0,1.0,0.5],draw=10000)
    
    Rasterize a dataset at 30 micron pixel size, with three kernels, using difference between scales.
    
    >>> X,Y,M,fig = tools.rasterize(x,y,dx=30.0,blur=[2.0,1.0,0.5],draw=10000, wavelet_magnitude=True)
        
        
    '''
    
    # set blur to a list
    if not isinstance(blur,list):
        blur = [blur]
    nb = len(blur)
    blur = np.array(blur)
    n = len(x)
    maxblur = np.max(blur) # for windowing
    
    
    
    if wavelet_magnitude and np.any(blur != np.sort(blur)[::-1]):
        raise Exception('When using wavelet magnitude, blurs must be sorted from greatest to least')
    
    minx = np.min(x)
    maxx = np.max(x)
    miny = np.min(y)
    maxy = np.max(y)
    minx,maxx = (minx+maxx)/2.0 - (maxx-minx)/2.0*expand, (minx+maxx)/2.0 + (maxx-minx)/2.0*expand
    miny,maxy = (miny+maxy)/2.0 - (maxy-miny)/2.0*expand, (miny+maxy)/2.0 + (maxy-miny)/2.0*expand
    X_ = np.arange(minx,maxx,dx)
    Y_ = np.arange(miny,maxy,dx)
    
    X = np.stack(np.meshgrid(X_,Y_)) # note this is xy order, not row col order

    W = np.zeros((X.shape[1],X.shape[2],nb))

    
    if draw: fig,ax = plt.subplots()
    count = 0
    
    g = np.resize(g,x.size)
    if(not (g==1.0).all()):
        g = normalize(g)
    
    for x_,y_,g_ in zip(x,y,g):
        # to speed things up I should index
        # to do this I'd have to find row and column indices
        #col = np.round((x_ - X_[0])/dx).astype(int)
        #row = np.round((y_ - X_[1])/dx).astype(int)
        #row0 = np.floor(row-blur*3).astype(int)
        #row1 = np.ceil(row+blur*3).astype(int)        
        #rows = np.arange(row0,row1+1)
        

        # this is incrementing one pixel at a time, it is way way faster, 
        # but doesn't use a kernel
        # I[c_,row,col] += 1.0
        # W[row,col] += 1.0
        if not use_windowing: # legacy version
            k = np.exp( - ( (X[0][...,None] - x_)**2 + (X[1][...,None] - y_)**2 )/(2.0*(dx*blur*2)**2)  )
            k /= np.sum(k,axis=(0,1),keepdims=True)
            k *= g_
            if wavelet_magnitude:
                for i in reversed(range(nb)):
                    if i == 0:
                        continue
                    k[...,i] = k[...,i] - k[...,i-1]

            W += k
        else: # use a small window
            r = int(np.ceil(maxblur*4))
            col = np.round((x_ - X_[0])/dx).astype(int)
            row = np.round((y_ - Y_[0])/dx).astype(int)
            
            row0 = np.floor(row-r).astype(int)
            row1 = np.ceil(row+r).astype(int)                    
            col0 = np.floor(col-r).astype(int)
            col1 = np.ceil(col+r).astype(int)
            # we need boundary conditions
            row0 = np.minimum(np.maximum(row0,0),W.shape[0]-1)
            row1 = np.minimum(np.maximum(row1,0),W.shape[0]-1)
            col0 = np.minimum(np.maximum(col0,0),W.shape[1]-1)
            col1 = np.minimum(np.maximum(col1,0),W.shape[1]-1)
            
           
            k =  np.exp( - ( (X[0][row0:row1+1,col0:col1+1,None] - x_)**2 + (X[1][row0:row1+1,col0:col1+1,None] - y_)**2 )/(2.0*(dx*blur*2)**2)  )
            k /= np.sum(k,axis=(0,1),keepdims=True)  
            k *= g_
            if wavelet_magnitude:
                for i in reversed(range(nb)):
                    if i == 0:
                        continue
                    k[...,i] = k[...,i] - k[...,i-1]
            W[row0:row1+1,col0:col1+1,:] += k #range of voxels -oka
            
        
            
        

        if draw:
            if not count%draw or count==(x.shape[0]-1):
                print(f'{count} of {x.shape[0]}')

                ax.cla()
                toshow = W-np.min(W,axis=(0,1),keepdims=True)
                toshow = toshow / np.max(toshow,axis=(0,1),keepdims=True)
                
                if nb >= 3:
                    toshow = toshow[...,:3]
                elif nb == 2:
                    toshow = toshow[...,[0,1,0]]
                elif nb == 1:
                    toshow = toshow[...,[0,0,0]]
                
                ax.imshow(np.abs(toshow))
                fig.canvas.draw()

        count += 1
    W = np.abs(W)
    # we will permute so channels are on first axis
    W = W.transpose((-1,0,1))
    extent = (X_[0],X_[-1],Y_[0],Y_[-1])
    
    # rename
    X = X_
    Y = Y_
    if draw:
        output = X,Y,W,fig
    else:
        output = X,Y,W
    return output
    

    
def rasterize_with_signal(x, y, s=None, dx=30.0, blur=1.0, expand=1.1, draw=0, wavelet_magnitude=False,use_windowing=True):
    ''' Rasterize a spatial transcriptomics dataset into a density image
    
    Paramters
    ---------
    x : numpy array of length N
        x location of cells
    y : numpy array of length N
        y location of cells
    s : numpy array of length N by M for M signals
        signal value should be length NxM
    dx : float
        Pixel size to rasterize data (default 30.0, in same units as x and y)
    blur : float or list of floats
        Standard deviation of Gaussian interpolation kernel.  Units are in 
        number of pixels.  Can be aUse a list to do multi scale.
    expand : float
        Factor to expand sampled area beyond cells. Defaults to 1.1.
    draw : int
        If True, draw a figure every draw points return its handle. Defaults to False (0).
    wavelet_magnitude : bool
        If True, take the absolute value of difference between scales for raster images.
        When using this option blur should be sorted from greatest to least.
    
        
    Returns
    -------
    X  : numpy array
        Locations of pixels along the x axis
    Y  : numpy array
        Locations of pixels along the y axis
    M : numpy array
        A rasterized image with len(blur) channels along the last axis
    fig : matplotlib figure handle
        If draw=True, returns a figure handle to the drawn figure.
        
    Raises
    ------    
    Exception 
        If wavelet_magnitude is set to true but blur is not sorted from greatest to least.
        
        
    
    Examples
    --------
    Rasterize a dataset at 30 micron pixel size, with three kernels.
    
    >>> X,Y,M,fig = tools.rasterize(x,y,dx=30.0,blur=[2.0,1.0,0.5],draw=10000)
    
    Rasterize a dataset at 30 micron pixel size, with three kernels, using difference between scales.
    
    >>> X,Y,M,fig = tools.rasterize(x,y,dx=30.0,blur=[2.0,1.0,0.5],draw=10000, wavelet_magnitude=True)
        
        
    '''
    
    # set blur to a list
    if not isinstance(blur,list):
        blur = [blur]
    nb = len(blur)
    blur = np.array(blur)
    n = len(x)
    maxblur = np.max(blur) # for windowing
    
    if len(blur)>1 and s is not None:
        raise Exception('when using a signal, we can only have one blur')
    if s is not None:
        s = np.array(s)
        if s.ndim == 1:
            s = s[...,None] # add a column of size 1
        
    
    
    if wavelet_magnitude and np.any(blur != np.sort(blur)[::-1]):
        raise Exception('When using wavelet magnitude, blurs must be sorted from greatest to least')
    
    minx = np.min(x)
    maxx = np.max(x)
    miny = np.min(y)
    maxy = np.max(y)
    minx,maxx = (minx+maxx)/2.0 - (maxx-minx)/2.0*expand, (minx+maxx)/2.0 + (maxx-minx)/2.0*expand
    miny,maxy = (miny+maxy)/2.0 - (maxy-miny)/2.0*expand, (miny+maxy)/2.0 + (maxy-miny)/2.0*expand
    X_ = np.arange(minx,maxx,dx)
    Y_ = np.arange(miny,maxy,dx)
    
    X = np.stack(np.meshgrid(X_,Y_)) # note this is xy order, not row col order

    if s is None:
        W = np.zeros((X.shape[1],X.shape[2],nb))
    else:
        W = np.zeros((X.shape[1],X.shape[2],s.shape[1]))
    

    
    if draw: fig,ax = plt.subplots()
    count = 0
    

    for x_,y_ in zip(x,y):
        # to speed things up I shoul index
        # to do this I'd have to find row and column indices
        #col = np.round((x_ - X_[0])/dx).astype(int)
        #row = np.round((y_ - X_[1])/dx).astype(int)
        #row0 = np.floor(row-blur*3).astype(int)
        #row1 = np.ceil(row+blur*3).astype(int)        
        #rows = np.arange(row0,row1+1)
        

        # this is incrementing one pixel at a time, it is way way faster, 
        # but doesn't use a kernel
        # I[c_,row,col] += 1.0
        # W[row,col] += 1.0
        if not use_windowing: # legacy version
            k = np.exp( - ( (X[0][...,None] - x_)**2 + (X[1][...,None] - y_)**2 )/(2.0*(dx*blur*2)**2)  )
            k /= np.sum(k,axis=(0,1),keepdims=True)*dx**2
            if wavelet_magnitude:
                for i in reversed(range(nb)):
                    if i == 0:
                        continue
                    k[...,i] = k[...,i] - k[...,i-1]
            if s is None:
                factor = 1.0
            else:
                factor = s[count]
            W += k * factor
        else: # use a small window
            r = int(np.ceil(maxblur*4))
            col = np.round((x_ - X_[0])/dx).astype(int)
            row = np.round((y_ - Y_[0])/dx).astype(int)
            
            row0 = np.floor(row-r).astype(int)
            row1 = np.ceil(row+r).astype(int)                    
            col0 = np.floor(col-r).astype(int)
            col1 = np.ceil(col+r).astype(int)
            # we need boundary conditions
            row0 = np.minimum(np.maximum(row0,0),W.shape[0]-1)
            row1 = np.minimum(np.maximum(row1,0),W.shape[0]-1)
            col0 = np.minimum(np.maximum(col0,0),W.shape[1]-1)
            col1 = np.minimum(np.maximum(col1,0),W.shape[1]-1)
            
           
            k = np.exp( - ( (X[0][row0:row1+1,col0:col1+1,None] - x_)**2 + (X[1][row0:row1+1,col0:col1+1,None] - y_)**2 )/(2.0*(dx*blur*2)**2)  )
            k /= np.sum(k,axis=(0,1),keepdims=True)*dx**2
            if wavelet_magnitude:
                for i in reversed(range(nb)):
                    if i == 0:
                        continue
                    k[...,i] = k[...,i] - k[...,i-1]
            if s is None:
                factor = 1.0
            else:
                factor = s[count]
            W[row0:row1+1,col0:col1+1,:] += k * factor
            
        
            
        

        if draw:
            if not count%draw or count==(x.shape[0]-1):
                print(f'{count} of {x.shape[0]}')

                ax.cla()
                toshow = W-np.min(W,axis=(0,1),keepdims=True)
                toshow = toshow / np.max(toshow,axis=(0,1),keepdims=True)
                
                if nb >= 3:
                    toshow = toshow[...,:3]
                elif nb == 2:
                    toshow = toshow[...,[0,1,0]]
                elif nb == 1:
                    toshow = toshow[...,[0,0,0]]
                
                ax.imshow(np.abs(toshow))
                fig.canvas.draw()

        count += 1
    W = np.abs(W)
    # we will permute so channels are on first axis
    W = W.transpose((-1,0,1))
    extent = (X_[0],X_[-1],Y_[0],Y_[-1])
    
    # rename
    X = X_
    Y = Y_
    if draw:
        output = X,Y,W,fig
    else:
        output = X,Y,W
    return output
    
        


def rasterizePCA(x, y, G):
    """Rasterize a spatial transcriptomics dataset into a density image and perform PCA on it.
    
    Parameters
    ----------
    x : numpy array of length N
        x location of cells
    y : numpy array of length N
        y location of cells
    G : pandas Dataframe of shape (N,M)
        gene expression level of cells for M genes
       
    Returns
    -------
    X : numpy array
        A rasterized image for each gene with the channel along the first axis
    Y : numpy array
        A rasterized image unraveled to 1-D for each gene; data is centered
    W : numpy array
        The eigenvalues for the principal components in descending order
    V : numpy array
        The normalized eigenvectors for the principal components
        V[:, i] corresponds to eigenvalue at W[i]
    Z : numpy array
        X rotated to align with the principal component axes
    nrows : int
        Row dimension of rasterized image
    ncols : int
        Column dimension of rasterized image
    
    Notes
    -----
    Each value/row at the same index in x, y, and G should all correspond to the same cell.
    x[i] <-> y[i] <-> G[i,:]
    
    """
    
    nrows=0
    ncols=0
    
    for i in range(G.shape[1]):
        g = np.array(G.iloc[:,i])
        
        XI,YI,I = tools.rasterize(x,y,g,dx=30.0,blur=1.0,expand=1.1, draw=0, 
                              wavelet_magnitude=True, use_windowing=True)
        
        if(i==0):
            # dimensions
            nrows=YI.size
            ncols=XI.size
            X = np.empty([G.shape[1], nrows, ncols])
            Y = np.empty([G.shape[1], nrows*ncols])
        
        # centers data
        X[i] = np.array(I)
        I_ = I.ravel()
        meanI = np.mean(I_)
        I_ -= meanI
        Y[i] = I_
        
        if(i % 50 == 0):
            print(f"{i} out of {G.shape[1]} genes rasterized.")
        
    S = np.cov(Y) # computes covariance matrix
    W,V = np.linalg.eigh(S) # W = eigenvalues, V = eigenvectors
    
    # reverses order to make it descending by eigenvalue
    W = W[::-1]
    V = V[:,::-1]
    Z = V.T @ Y
    
    return X, Y, W, V, Z, nrows, ncols


def make_scree(W, name, p=6):
    """Create a scree plot for a given set of eigenvalues.
    
    Parameters
    ----------
    W : numpy array
        Eigenvalues in descending order
    name : str
        Name of the dataset the eigenvalue originate from
        Will be included in plot title
    p : int
        Number of eigenvalue from W to plot. Defaults to 6
        
    Returns
    -------
    fig : matplotlib Figure
        Figure handle for scree plot
        
    Raises
    ------
    ValueError
        If p is larger than the number to eigenvalues in W.
    
    """
    
    if(p > W.size):
        raise ValueError("Cannot plot more eigenvalues than what is given.")
    
    fig, ax = plt.subplots(figsize=(8,6))
    
    ax.bar(range(p),W[:p],width=.9,tick_label=range(1,p+1))
    ax.plot(W[:p],'ko-',linewidth=1)

    ax.set_title("Scree Plot: First %s PCs (%s)" %(p,name), fontsize=18,fontweight='bold')
    ax.set_xlabel('Principal Components')
    ax.set_ylabel("Eigenvalues")
        
    return fig


def saveRasters(X, Y, W, V, Z, scree_fig, nrows, ncols, name, path):
    """
    Create an RGB image principal components 1-3 and 4-6.
    Save numpy arrays necessary for rasterizing and performing PCA.
    Save scree plot and RGB image as jpeg, and their figure handles.
    
    Parameters
    ----------
    X : numpy array
        A rasterized image for each gene with the channel along the first axis
    Y : numpy array
        A rasterized image unraveled to 1-D for each gene; data is centered
    W : numpy array
        The eigenvalues for the principal components in descending order
    V : numpy array
        The normalized eigenvectors for the principal components
        V[:, i] corresponds to eigenvalue at W[i]
    Z : numpy array
        X rotated to align with the principal component axes
    scree_fig : matplotlib Figure
        Figure handle for scree plot
    nrows : int
        Row dimension of rasterized image
    ncols : int
        Column dimension of rasterized image
    name : str
        Name of the dataset the arrays originate from
    path : str
        Absolute path to folder where files should be saved
        
    Returns
    -------
    None
    
    """
    
    I_pca = Z.reshape((Z.shape[0], nrows, ncols))
    I_rgb = np.array(I_pca[:6].transpose(1,2,0))
    I_rgb[...,:] = normalize(I_rgb[...,:])
    
    fig,axs = plt.subplots(1,2)
    axs[0].imshow(I_rgb[:,:,:3])
    axs[1].imshow(I_rgb[:,:,3:])
    
    fig.suptitle("Principal Components in RGB (%s)" %name, fontsize=18,fontweight='bold')
    axs[0].set_title("PCs 1-3")
    axs[1].set_title("PCs 4-6")
    
    scree_fig.savefig(f"{path}{name}_screefig.jpg")
    fig.savefig(f"{path}{name}_rgbfig.jpg")
    np.savez(f"{path}{name}_arrays", X=X, Y=Y, W=W, V=V, Z=Z, I_pca=I_pca, I_rgb=I_rgb,
             screefig=scree_fig, rgbfig = fig)
    
    print(f"Saved all {name} files!\n")
    
    

    
def interp(x,I,phii,**kwargs):
    '''
    Interpolate the 2D image I, with regular grid positions stored in x (1d arrays),
    at the positions stored in phii (3D arrays with first channel storing component)    
    
    Parameters
    ----------
    x : list of arrays
        List of arrays storing the pixel locations along each image axis. convention is row column order not xy.
    I : array
        Image array. First axis should contain different channels.
    phii : array
        Sampling array. First axis should contain sample locations corresponding to each axis.
    **kwargs : dict
        Other arguments fed into the torch interpolation function torch.nn.grid_sample
        
    
    Returns
    -------
    out : torch tensor
            The image I resampled on the points defined in phii.
    
    Notes
    -----
    Convention is to use align_corners=True.
    
    This uses the torch library.
    '''
    
    # first we have to normalize phii to the range -1,1    
    I = torch.as_tensor(I)
    phii = torch.as_tensor(phii)
    phii = torch.clone(phii)
    for i in range(2):
        phii[i] -= x[i][0]
        phii[i] /= x[i][-1] - x[i][0]
    # note the above maps to 0,1
    phii *= 2.0
    # to 0 2
    phii -= 1.0
    # done

    # NOTE I should check that I can reproduce identity
    # note that phii must now store x,y,z along last axis
    # is this the right order?
    # I need to put batch (none) along first axis
    # what order do the other 3 need to be in?    
    out = grid_sample(I[None],phii.flip(0).permute((1,2,0))[None],align_corners=True,**kwargs)
    # note align corners true means square voxels with points at their centers
    # post processing, get rid of batch dimension

    return out[0]

# build an interp function from grid sample
def interp3D(x,I,phii,**kwargs):
    '''
    Interpolate the 3D image I, with regular grid positions stored in x (1d arrays),
    at the positions stored in phii (4D arrays with first channel storing component)    
    
    Parameters
    ----------
    x : list of arrays
        List of arrays storing the pixel locations along each image axis. convention is row column order not xy.
    I : array
        Image array. First axis should contain different channels.
    phii : array
        Sampling array. First axis should contain sample locations corresponding to each axis.
    **kwargs : dict
        Other arguments fed into the torch interpolation function torch.nn.grid_sample
        
    
    Returns
    -------
    out : torch tensor
            The image I resampled on the points defined in phii.
    
    Notes
    -----
    Convention is to use align_corners=True.
    
    This uses the torch library.
    '''
    # first we have to normalize phii to the range -1,1    
    I = torch.as_tensor(I)
    phii = torch.as_tensor(phii)
    phii = torch.clone(phii)
    for i in range(3):
        phii[i] -= x[i][0]
        phii[i] /= x[i][-1] - x[i][0]
    # note the above maps to 0,1
    phii *= 2.0
    # to 0 2
    phii -= 1.0
    # done

    # NOTE I should check that I can reproduce identity
    # note that phii must now store x,y,z along last axis
    # is this the right order?
    # I need to put batch (none) along first axis
    # what order do the other 3 need to be in?    
    out = grid_sample(I[None],phii.flip(0).permute((1,2,3,0))[None],align_corners=True,**kwargs)
    # note align corners true means square voxels with points at their centers
    # post processing, get rid of batch dimension

    return out[0]


def clip(I):
    ''' clip an arrays values between 0 and 1.  Useful for visualizing
    
    Parameters
    ----------
    I : torch tensor
        A torch tensor to clip.
    
    Returns
    -------
    Ic : torch tensor
        Clipped torch tensor.
    
    '''
    Ic = torch.clone(I)
    Ic[Ic<0]=0
    Ic[Ic>1]=1
    return Ic


# timesteps will be along the first axis
def v_to_phii(xv,v):
    ''' Integrate a velocity field over time to return a position field (diffeomorphism).
    
    Parameters
    ----------
    xv : list of torch tensor 
        List of 1D tensors describing locations of sample points in v
    v : torch tensor
        5D (nt,2,v0,v1) velocity field
    
    Returns
    -------
    phii: torch tensor
        Inverse map (position field) computed by method of characteristics

    '''
    
    XV = torch.stack(torch.meshgrid(xv))
    phii = torch.clone(XV)
    dt = 1.0/v.shape[0]
    for t in range(v.shape[0]):
        Xs = XV - v[t]*dt
        phii = interp(xv,phii-XV,Xs)+Xs
    return phii

def v_to_phii_3D(xv,v):
    ''' Integrate a 3D velocity field over time to return a 3D position field (diffeomorphism).
    
    Parameters
    ----------
    xv : list of torch tensor 
        List of 1D tensors describing locations of sample points in v
    v : torch tensor
        5D (nt,3,v0,v1,v2) velocity field
    
    Returns
    -------
    phii: torch tensor
        Inverse map (position field) computed by method of characteristics

    '''
    
    XV = torch.stack(torch.meshgrid(xv))
    phii = torch.clone(XV)
    dt = 1.0/v.shape[0]
    for t in range(v.shape[0]):
        Xs = XV - v[t]*dt
        phii = interp3D(xv,phii-XV,Xs)+Xs
    return phii

def to_A(L,T):
    ''' Convert a linear transform matrix and a translation vector into an affine matrix.
    
    Parameters
    ----------
    L : torch tensor
        2x2 linear transform matrix
        
    T : torch tensor
        2 element translation vector (note NOT 2x1)
        
    Returns
    -------
    
    A : torch tensor
        Affine transform matrix
        
        
    '''
    O = torch.tensor([0.,0.,1.],device=L.device,dtype=L.dtype)
    A = torch.cat((torch.cat((L,T[:,None]),1),O[None]))
    return A
def to_A_3D(L,T):
    ''' Convert a linear transform matrix and a translation vector into an affine matrix.
    
    Parameters
    ----------
    L : torch tensor
        3x3 linear transform matrix
        
    T : torch tensor
        3 element translation vector (note NOT 2x1)
        
    Returns
    -------
    
    A : torch tensor
        Affine transform matrix
        
        
    '''
    O = torch.tensor([0.,0.,0.0,1.],device=L.device,dtype=L.dtype)
    A = torch.cat((torch.cat((L,T[:,None]),1),O[None]))
    return A

def extent_from_x(xJ):
    ''' Given a set of pixel locations, returns an extent 4-tuple for use with np.imshow.
    
    Note inputs are locations of pixels along each axis, i.e. row column not xy.
    
    Parameters
    ----------
    xJ : list of torch tensors
        Location of pixels along each axis
    
    Returns
    -------
    extent : tuple
        (xmin, xmax, ymin, ymax) tuple
    
    Examples
    --------
    
    >>> extent_from_x(xJ)
    >>> fig,ax = plt.subplots()
    >>> ax.imshow(J,extent=extentJ)
    
    '''
    dJ = [x[1]-x[0] for x in xJ]
    extentJ = ( (xJ[1][0] - dJ[1]/2.0).item(),
               (xJ[1][-1] + dJ[1]/2.0).item(),
               (xJ[0][-1] + dJ[0]/2.0).item(),
               (xJ[0][0] - dJ[0]/2.0).item())
    return extentJ

def L_T_from_points(pointsI,pointsJ):
    '''
    Compute an affine transformation from points.
    
    Note for an affine transformation (6dof) we need 3 points.
    
    Outputs, L,T should be rconstructed blockwize like [L,T;0,0,1]
    
    Parameters
    ----------
    pointsI : array
        An Nx2 array of floating point numbers describing atlas points in ROW COL order (not xy)
    pointsJ : array
        An Nx2 array of floating point numbers describing target points in ROW COL order (not xy)
    
    Returns
    -------
    L : array
        A 2x2 linear transform array.
    T : array
        A 2 element translation vector
    '''
    if pointsI is None or pointsJ is None:
        raise Exception('Points are set to None')
        
    nI = pointsI.shape[0]
    nJ = pointsJ.shape[0]
    if nI != nJ:
        raise Exception(f'Number of pointsI ({nI}) is not equal to number of pointsJ ({nJ})')
    if pointsI.shape[1] != 2:
        raise Exception(f'Number of components of pointsI ({pointsI.shape[1]}) should be 2')
    if pointsJ.shape[1] != 2:
        raise Exception(f'Number of components of pointsJ ({pointsJ.shape[1]}) should be 2')
    # transformation model
    if nI < 3:
        # translation only 
        L = np.eye(2)
        T = np.mean(pointsJ,0) - np.mean(pointsI,0)
    else:
        # we need an affine transform
        pointsI_ = np.concatenate((pointsI,np.ones((nI,1))),1)
        pointsJ_ = np.concatenate((pointsJ,np.ones((nI,1))),1)
        II = pointsI_.T@pointsI_
        IJ = pointsI_.T@pointsJ_
        A = (np.linalg.inv(II)@IJ).T        
        L = A[:2,:2]
        T = A[:2,-1]
    return L,T


def LDDMM(xI,I,xJ,J,pointsI=None,pointsJ=None,
          L=None,T=None,A=None,v=None,xv=None,
          a=500.0,p=2.0,expand=2.0,nt=3,
         niter=5000,diffeo_start=0, epL=2e-8, epT=2e-1, epV=2e3,
         sigmaM=1.0,sigmaB=2.0,sigmaA=5.0,sigmaR=5e5,sigmaP=2e1,
          device='cpu',dtype=torch.float64, muB=None, muA=None):
    ''' Run LDDMM between a pair of images.
    
    This jointly estimates an affine transform A, and a diffeomorphism phi.
    The map is off the form x -> A phi x
    
    
    Parameters
    ----------
    xI : list of torch tensor
        Location of voxels in atlas image I
    I : torch tensor
        Atlas image I, with channels along first axis        
    xJ : list of torch tensor
        Location of voxels in target image J
    J : torch tensor
        Target image J, with channels along first axis
    L : torch tensor
        Initial guess for linear transform (2x2 torch tensor). Defaults to None (identity).
    T : torch tensor
        Initial guess for translation (2 element torch tensor). Defaults to None (identity)
    A : torch tensor
        Initial guess for affine matrix.  Either L and T can be specified, or A, but not both.
        Defaults to None (identity).
    v : torch tensor
        Initial guess for velocity field
    xv : torch tensor
        pixel locations for velocity field
    a : float
        Smoothness scale of velocity field (default 500.0)
    p : float
        Power of Laplacian in velocity regularization (default 2.0)
    expand : float
        Factor to expand size of velocity field around image boundaries (default 2.0)
    nt : int
        Number of timesteps for integrating velocity field (default 3). Ignored if you input v.
    pointsI : torch tensor
        N x 2 set of corresponding points for matching in atlas image. Default None (no points).
    pointsJ : torch tensor
        N x 2 set of corresponding points for matching in target image. Default None (no points).
    niter : int
        Number of iterations of gradient descent optimization
    diffeo_start : int
        Number of iterations of gradient descent optimization for affine only, before nonlinear deformation.
    epL : float
        Gradient descent step size for linear part of affine.
    epT : float
        Gradient descent step size of translation part of affine.
    epV : float
        Gradient descent step size for velocity field.
    sigmaM : float
        Standard deviation of image matching term for Gaussian mixture modeling in cost function. 
        This term generally controls matching accuracy with smaller corresponding to more accurate.
        As an common example (rule of thumb), you could chose this parameter to be the variance of the pixels
        in your target image.
    sigmaB : float
        Standard deviation of backtround term for Gaussian mixture modeling in cost function. 
        If there is missing tissue in target, we may label some pixels in target as background,
        and not enforce matching here.
    sigmaA : float
        Standard deviation of artifact term for Gaussian mixture modeling in cost function. 
        If there are artifacts in target or other lack of corresponding between template and target, 
        we may label some pixels in target as artifact, and not enforce matching here.
    sigmaR: float
        Standard deviation for regularization. Smaller sigmaR means a smoother resulting transformation. 
        Regularization is of the form: 0.5/sigmaR^2 int_0^1 int_X |Lv|^2 dx dt. 
    sigmaP: float
        Standard deviation for matching of points.  
        Cost is of the form 0.5/sigmaP^2 sum_i (atlas_point_i - target_point_i)^2
    device: str
        torch device. defaults to 'cpu'. Can also be 'cuda:0' for example.
    dtype: torch dtype
        torch data type. defaults to torch.float64
    muA: torch tensor whose dimension is the same as the target image
        Defaults to None, which means we estimate this. If you provide a value, we will not estimate it.
        If the target is a RGB image, this should be a tensor of size 3.
        If the target is a grayscale image, this should be a tensor of size 1.
    muB: torch tensor whose dimension is the same as the target image
        Defaults to None, which means we estimate this. If you provide a value, we will not estimate it.
        
    Returns a dictionary
    -------
    {
    'A': torch tensor
        Affine transform
    'v': torch tensor
        Velocity field
    'xv': list of torch tensor
        Pixel locations in v
    'WM': torch tensor
        Resulting weight 2D array (matching)
    'WB': torch tensor
        Resulting weight 2D array (background)
    'WA': torch tensor
        Resulting weight 2D array (artifact)
    }
    
    '''
    
    
    
    
    # todo
    # implement local?
    # more iters

    #niter = 2000
    #diffeo_start = 100
    #epL = 5e-11
    #epT = 5e-4
    #epV = 5e1
    #niter = 5000

    # check initial inputs
    if A is not None:
        # if we specify an A
        if L is not None or T is not None:
            raise Exception('If specifying A, you must not specify L or T')
        L = torch.tensor(A[:2,:2],device=device,dtype=dtype,requires_grad=True)
        T = torch.tensor(A[:2,-1],device=device,dtype=dtype,requires_grad=True)   
    else:
        # if we do not specify A                
        if L is None: L = torch.eye(2,device=device,dtype=dtype,requires_grad=True)
        if T is None: T = torch.zeros(2,device=device,dtype=dtype,requires_grad=True)
    L = torch.tensor(L,device=device,dtype=dtype,requires_grad=True)
    T = torch.tensor(T,device=device,dtype=dtype,requires_grad=True)
    # change to torch
    I = torch.tensor(I,device=device,dtype=dtype)                         
    J = torch.tensor(J,device=device,dtype=dtype)
    
    #L = torch.eye(2,device=device,dtype=dtype,requires_grad=True)
    ##L.data[0,0] = -1.0
    ##L.data[:2,:2] *= 0.2
    #T = torch.zeros(2,device=device,dtype=dtype,requires_grad=True)
    ##T.data[0] = +2000.0
    ##T.data[1] = -xI[1][-1]/2.0*1.1

    # velocity
    #a = 500.0
    #p = 3.0
    #expand = 2.0
    if v is not None and xv is not None:
        v = torch.tensor(v,device=device,dtype=dtype,requires_grad=True)
        xv = [torch.tensor(x,device=device,dtype=dtype) for x in xv]
        XV = torch.stack(torch.meshgrid(xv),-1)
        nt = v.shape[0]        
    elif v is None and xv is None:
        minv = torch.as_tensor([x[0] for x in xI],device=device,dtype=dtype)
        maxv = torch.as_tensor([x[-1] for x in xI],device=device,dtype=dtype)
        minv,maxv = (minv+maxv)*0.5 + 0.5*torch.tensor([-1.0,1.0],device=device,dtype=dtype)[...,None]*(maxv-minv)*expand
        xv = [torch.arange(m,M,a*0.5,device=device,dtype=dtype) for m,M in zip(minv,maxv)]
        XV = torch.stack(torch.meshgrid(xv),-1)
        v = torch.zeros((nt,XV.shape[0],XV.shape[1],XV.shape[2]),device=device,dtype=dtype,requires_grad=True)
    else:
        raise Exception(f'If inputting an initial v, must input both xv and v')
    extentV = extent_from_x(xv)
    dv = torch.as_tensor([x[1]-x[0] for x in xv],device=device,dtype=dtype)
    
    
 
    fv = [torch.arange(n,device=device,dtype=dtype)/n/d for n,d in zip(XV.shape,dv)]
    extentF = extent_from_x(fv)
    FV = torch.stack(torch.meshgrid(fv),-1)
    LL = (1.0 + 2.0*a**2* torch.sum( (1.0 - torch.cos(2.0*np.pi*FV*dv))/dv**2 ,-1))**(p*2.0)

    K = 1.0/LL
    #fig,ax = plt.subplots()
    #ax.imshow(K,vmin=0.0,vmax=0.1,extent=extentF)
    
    #fig,ax = plt.subplots()
    #ax.imshow(K[0].cpu())
    DV = torch.prod(dv)
    Ki = torch.fft.ifftn(K).real
    fig,ax = plt.subplots()
    ax.imshow(Ki.clone().detach().cpu().numpy(),vmin=0.0,extent=extentV)
    ax.set_title('smoothing kernel')
    fig.canvas.draw()


    # nt = 3
    


    WM = torch.ones(J[0].shape,dtype=J.dtype,device=J.device)*0.5
    WB = torch.ones(J[0].shape,dtype=J.dtype,device=J.device)*0.4
    WA = torch.ones(J[0].shape,dtype=J.dtype,device=J.device)*0.1
    if pointsI is None and pointsJ is None:
        pointsI = torch.zeros((0,2),device=J.device,dtype=J.dtype)
        pointsJ = torch.zeros((0,2),device=J.device,dtype=J.dtype) 
    elif (pointsI is None and pointsJ is not None) or (pointsJ is None and pointsI is not None):
        raise Exception('Must specify corresponding sets of points or none at all')
    else:
        pointsI = torch.tensor(pointsI,device=J.device,dtype=J.dtype)
        pointsJ = torch.tensor(pointsJ,device=J.device,dtype=J.dtype)
    
    
    xI = [torch.tensor(x,device=device,dtype=dtype) for x in xI]
    xJ = [torch.tensor(x,device=device,dtype=dtype) for x in xJ]
    XI = torch.stack(torch.meshgrid(*xI,indexing='ij'),-1)
    XJ = torch.stack(torch.meshgrid(*xJ,indexing='ij'),-1)
    dJ = [x[1]-x[0] for x in xJ]
    extentJ = (xJ[1][0].item()-dJ[1].item()/2.0,
          xJ[1][-1].item()+dJ[1].item()/2.0,
          xJ[0][-1].item()+dJ[0].item()/2.0,
          xJ[0][0].item()-dJ[0].item()/2.0)
    
    #sigmaM = 0.2
    #sigmaB = 0.19
    #sigmaA = 0.3
    #sigmaR = 5e5
    #sigmaP = 2e-1
    
    if muA is None:
        estimate_muA = True
    else:
        estimate_muA = False
    if muB is None:
        estimate_muB = True
    else:
        estimate_muB = False
    
    fig,ax = plt.subplots(2,3)
    ax = ax.ravel()
    figE,axE = plt.subplots(1,3)
    Esave = []

    try:
        L.grad.zero_()
    except:
        pass
    try:
        T.grad.zero_()
    except:
        pass
    for it in range(niter):
        # make A
        A = to_A(L,T)
        # Ai
        Ai = torch.linalg.inv(A)
        # transform sample points
        Xs = (Ai[:2,:2]@XJ[...,None])[...,0] + Ai[:2,-1]    
        # now diffeo, not semilagrange here
        for t in range(nt-1,-1,-1):
            Xs = Xs + interp(xv,-v[t].permute(2,0,1),Xs.permute(2,0,1)).permute(1,2,0)/nt
        # and points
        pointsIt = torch.clone(pointsI)
        if pointsIt.shape[0] >0:
            for t in range(nt):            
                pointsIt += interp(xv,v[t].permute(2,0,1),pointsIt.T[...,None])[...,0].T/nt
            pointsIt = (A[:2,:2]@pointsIt.T + A[:2,-1][...,None]).T

        # transform image
        AI = interp(xI,I,Xs.permute(2,0,1),padding_mode="border")

        # transform the contrast
        B = torch.ones(1+AI.shape[0],AI.shape[1]*AI.shape[2],device=AI.device,dtype=AI.dtype)
        B[1:AI.shape[0]+1] = AI.reshape(AI.shape[0],-1)
        #B = torch.ones(10,AI.shape[1]*AI.shape[2],device=AI.device,dtype=AI.dtype)
        #B[1:4] = AI.reshape(AI.shape[0],-1)
        #B[4] = (AI[0][None]**2).reshape(1,-1)
        #B[5] = (AI[1][None]**2).reshape(1,-1)
        #B[6] = (AI[2][None]**2).reshape(1,-1)
        #B[7] = (AI[0][None]*AI[1][None]).reshape(1,-1)
        #B[8] = (AI[0][None]*AI[2][None]).reshape(1,-1)
        #B[9] = (AI[1][None]*AI[2][None]).reshape(1,-1)
        with torch.no_grad():    
            BB = B@(B*WM.ravel()).T
            BJ = B@((J*WM).reshape(J.shape[0],J.shape[1]*J.shape[2])).T
            small = 0.1
            coeffs = torch.linalg.solve(BB + small*torch.eye(BB.shape[0],device=BB.device,dtype=BB.dtype),BJ)
        fAI = ((B.T@coeffs).T).reshape(J.shape)

        # objective function
        EM = torch.sum((fAI - J)**2*WM)/2.0/sigmaM**2
        ER = torch.sum(torch.sum(torch.abs(torch.fft.fftn(v,dim=(1,2)))**2,dim=(0,-1))*LL)*DV/2.0/v.shape[1]/v.shape[2]/sigmaR**2
        E = EM + ER
        tosave = [E.item(), EM.item(), ER.item()]
        if pointsIt.shape[0]>0:
            EP = torch.sum((pointsIt - pointsJ)**2)/2.0/sigmaP**2
            E += EP
            tosave.append(EP.item())
        
        Esave.append( tosave )
        # gradient update
        E.backward()
        with torch.no_grad():            
            L -= (epL/(1.0 + (it>=diffeo_start)*9))*L.grad
            T -= (epT/(1.0 + (it>=diffeo_start)*9))*T.grad

            L.grad.zero_()
            T.grad.zero_()
            

            # v grad
            vgrad = v.grad
            # smooth it
            vgrad = torch.fft.ifftn(torch.fft.fftn(vgrad,dim=(1,2))*K[...,None],dim=(1,2)).real
            if it >= diffeo_start:
                v -= vgrad*epV
            v.grad.zero_()


        # update weights
        if not it%5:
            with torch.no_grad():
                # M step for these params
                if estimate_muA:
                    muA = torch.sum(WA*J,dim=(-1,-2))/torch.sum(WA)
                if estimate_muB:
                    muB = torch.sum(WB*J,dim=(-1,-2))/torch.sum(WB)
                #if it <= 200:
                #    muA = torch.tensor([0.75,0.77,0.79],device=J.device,dtype=J.dtype)
                #    muB = torch.ones(J.shape[0],device=J.device,dtype=J.dtype)*0.96

                if it >= 50:

                    W = torch.stack((WM,WA,WB))
                    pi = torch.sum(W,dim=(1,2))
                    pi += torch.max(pi)*1e-6
                    pi /= torch.sum(pi)


                    # now the E step, update the weights
                    WM = pi[0]* torch.exp( -torch.sum((fAI - J)**2,0)/2.0/sigmaM**2 )/np.sqrt(2.0*np.pi*sigmaM**2)**J.shape[0]
                    WA = pi[1]* torch.exp( -torch.sum((muA[...,None,None] - J)**2,0)/2.0/sigmaA**2 )/np.sqrt(2.0*np.pi*sigmaA**2)**J.shape[0]
                    WB = pi[2]* torch.exp( -torch.sum((muB[...,None,None] - J)**2,0)/2.0/sigmaB**2 )/np.sqrt(2.0*np.pi*sigmaB**2)**J.shape[0]
                    WS = WM+WB+WA
                    WS += torch.max(WS)*1e-6
                    WM /= WS
                    WB /= WS
                    WA /= WS


        # draw
        if not it%10:
            ax[0].cla()
            ax[0].imshow(   ((AI-torch.amin(AI,(1,2))[...,None,None])/(torch.amax(AI,(1,2))-torch.amin(AI,(1,2)))[...,None,None]).permute(1,2,0).clone().detach().cpu(),extent=extentJ)
            ax[0].scatter(pointsIt[:,1].clone().detach().cpu(),pointsIt[:,0].clone().detach().cpu())
            ax[0].set_title('space tformed atlas')
            
            ax[1].cla()    
            ax[1].imshow(clip(fAI.permute(1,2,0).clone().detach()/torch.max(J).item()).cpu(),extent=extentJ)
            ax[1].scatter(pointsIt[:,1].clone().detach().cpu(),pointsIt[:,0].clone().detach().cpu())
            ax[1].set_title('contrast tformed atlas')
            
            ax[5].cla()
            ax[5].imshow(clip( (fAI - J)/(torch.max(J).item())*3.0  ).permute(1,2,0).clone().detach().cpu()*0.5+0.5,extent=extentJ)
            ax[5].scatter(pointsIt[:,1].clone().detach().cpu(),pointsIt[:,0].clone().detach().cpu())
            ax[5].scatter(pointsJ[:,1].clone().detach().cpu(),pointsJ[:,0].clone().detach().cpu())
            ax[5].set_title('Error')

            ax[2].cla()
            ax[2].imshow(J.permute(1,2,0).cpu()/torch.max(J).item(),extent=extentJ)
            ax[2].scatter(pointsJ[:,1].clone().detach().cpu(),pointsJ[:,0].clone().detach().cpu())
            ax[2].set_title('Target')

            ax[4].cla()
            ax[4].imshow(clip(torch.stack((WM,WA,WB),-1).clone().detach()).cpu(),extent=extentJ)
            ax[4].set_title('Weights')


            toshow = v[0].clone().detach().cpu()
            toshow /= torch.max(torch.abs(toshow))
            toshow = toshow*0.5+0.5
            toshow = torch.cat((toshow,torch.zeros_like(toshow[...,0][...,None])),-1)   
            ax[3].cla()
            ax[3].imshow(clip(toshow),extent=extentV)
            ax[3].set_title('velocity')
            
            axE[0].cla()
            axE[0].plot(Esave)
            axE[0].legend(['E','EM','ER','EP'])
            axE[0].set_yscale('log')
            axE[1].cla()
            axE[1].plot([e[:2] for e in Esave])
            axE[1].legend(['E','EM'])
            axE[1].set_yscale('log')
            axE[2].cla()
            axE[2].plot([e[2] for e in Esave])
            axE[2].legend(['ER'])
            axE[2].set_yscale('log')


            fig.canvas.draw()
            figE.canvas.draw()
            
    return {
        'A': A.clone().detach(), 
        'v': v.clone().detach(), 
        'xv': xv, 
        'WM': WM.clone().detach(),
        'WB': WB.clone().detach(),
        'WA': WA.clone().detach()
    }


def LDDMM_3D_to_slice(xI,I,xJ,J,pointsI=None,pointsJ=None,
        L=None,T=None,A=None,v=None,xv=None,
        a=500.0,p=2.0,expand=1.25,nt=3,
        niter=5000,diffeo_start=0, epL=1e-6, epT=1e1, epV=1e3,
        sigmaM=1.0,sigmaB=2.0,sigmaA=5.0,sigmaR=1e8,sigmaP=2e1,
        device='cpu',dtype=torch.float64, muA=None, muB = None):
    ''' LDDMM for 3D to 2D slice mapping.
    
    muA: torch tensor whose dimension is the same as the target image
        Defaults to None, which means we estimate this. If you provide a value, we will not estimate it.
        If the target is a RGB image, this should be a tensor of size 3.
        If the target is a grayscale image, this should be a tensor of size 1.
    muB: torch tensor whose dimension is the same as the target image
        Defaults to None, which means we estimate this. If you provide a value, we will not estimate it.
        

    '''
    if muA is None:
        estimate_muA = True
    else:
        estimate_muA = False
    if muB is None:
        estimate_muB = True
    else:
        estimate_muB = False
        
    # check initial inputs and convert to torch
    if A is not None:
        # if we specify an A
        if L is not None or T is not None:
            raise Exception('If specifying A, you must not specify L or T')
        L = torch.tensor(A[:3,:3],device=device,dtype=dtype,requires_grad=True)
        T = torch.tensor(A[:3,-1],device=device,dtype=dtype,requires_grad=True)   
    else:
        # if we do not specify A                
        if L is None: L = torch.eye(3,device=device,dtype=dtype,requires_grad=True)
        if T is None: T = torch.zeros(3,device=device,dtype=dtype,requires_grad=True)
    
    L = torch.tensor(L,device=device,dtype=dtype,requires_grad=True)
    T = torch.tensor(T,device=device,dtype=dtype,requires_grad=True)
    # change to torch
    I = torch.tensor(I,device=device,dtype=dtype)                         
    J = torch.tensor(J,device=device,dtype=dtype)
    if J.ndim == 3:
        J = J[:,None] # add a z slice dimension


    if v is not None and xv is not None:
        v = torch.tensor(v,device=device,dtype=dtype,requires_grad=True)
        xv = [torch.tensor(x,device=device,dtype=dtype) for x in xv]
        XV = torch.stack(torch.meshgrid(xv),-1)
        nt = v.shape[0]        
    elif v is None and xv is None:
        minv = torch.as_tensor([x[0] for x in xI],device=device,dtype=dtype)
        maxv = torch.as_tensor([x[-1] for x in xI],device=device,dtype=dtype)
        minv,maxv = (minv+maxv)*0.5 + 0.5*torch.tensor([-1.0,1.0],device=device,dtype=dtype)[...,None]*(maxv-minv)*expand
        xv = [torch.arange(m,M,a*0.5,device=device,dtype=dtype) for m,M in zip(minv,maxv)]
        XV = torch.stack(torch.meshgrid(xv),-1)
        v = torch.zeros((nt,XV.shape[0],XV.shape[1],XV.shape[2],XV.shape[3]),device=device,dtype=dtype,requires_grad=True)
        
    else:
        raise Exception(f'If inputting an initial v, must input both xv and v')
    extentV = extent_from_x(xv[1:])
    dv = torch.as_tensor([x[1]-x[0] for x in xv],device=device,dtype=dtype)
    
    
 
    fv = [torch.arange(n,device=device,dtype=dtype)/n/d for n,d in zip(XV.shape,dv)]
    extentF = extent_from_x(fv[1:])
    FV = torch.stack(torch.meshgrid(fv),-1)
    LL = (1.0 + 2.0*a**2* torch.sum( (1.0 - torch.cos(2.0*np.pi*FV*dv))/dv**2 ,-1))**(p*2.0)

    K = 1.0/LL
    #fig,ax = plt.subplots()
    #ax.imshow(K,vmin=0.0,vmax=0.1,extent=extentF)
    
    #fig,ax = plt.subplots()
    #ax.imshow(K[0].cpu())
    DV = torch.prod(dv)
    Ki = torch.fft.ifftn(K).real
    fig,ax = plt.subplots()
    ax.imshow(Ki[Ki.shape[0]//2].clone().detach().cpu().numpy(),vmin=0.0,extent=extentV)
    ax.set_title('smoothing kernel')
    fig.canvas.draw()

    # steps
    epL = torch.tensor(epL,device=device,dtype=dtype)
    epT = torch.tensor(epT,device=device,dtype=dtype)

    # initialize weights
    WM = torch.ones(J[0].shape,dtype=J.dtype,device=J.device)*0.5
    WB = torch.ones(J[0].shape,dtype=J.dtype,device=J.device)*0.4
    WA = torch.ones(J[0].shape,dtype=J.dtype,device=J.device)*0.1

    # locations of pixels
    extentI = extent_from_x(xI[1:]) 
    xI = [torch.tensor(x,device=device,dtype=dtype) for x in xI]
    if len(xJ) == 2:
        xJ = [[0.0],xJ[0],xJ[1]]    
    extentJ = extent_from_x(xJ[1:])
    xJ = [torch.tensor(x,device=device,dtype=dtype) for x in xJ]
    XI = torch.stack(torch.meshgrid(*xI,indexing='ij'),-1)
    XJ = torch.stack(torch.meshgrid(*xJ,indexing='ij'),-1)
    dJ = [x[1]-x[0] for x in xJ[1:]]
    
    # metric TODO
    '''
    g = torch.zeros((12,12))
    count = 0
    for i in range(12):
        Ei = (torch.arange(16)==i).reshape((4,4))*1.0
        EiXI = (Ei[:3,:3]@XI[...,None])[...,0] + Ei[:3,-1]
        for j in range(12):
            Ej = (torch.arange(16)==j).reshape((4,4))*1.0
            EjXI = (Ej[:3,:3]@XI[...,None])[...,0] + Ej[:3,-1]
            g[i,j] = torch.mean(torch.sum(EiXI*EjXI,-1))

    gi = torch.inverse(g)
    fig,ax = plt.subplots(1,2)
    ax[0].imshow(g)
    ax[0].set_title('Metric') 
    ax[1].imshow(gi)    
    ax[1].set_title('Inverse metric') 
    fig.canvas.draw()
    '''

    # a figure
    fig,ax = plt.subplots(2,3)
    ax = ax.ravel()
    figE,axE = plt.subplots(1,3)
    axE = axE.ravel()
    Esave = []
    # zero gradients
    try:
        L.grad.zero_()
    except:
        pass
    try:
        T.grad.zero_()
    except:
        pass


    for it in range(niter):
        # make A
        A = to_A_3D(L,T)
        # Ai
        Ai = torch.linalg.inv(A)
        # transform sample points        
        Xs = (Ai[:-1,:-1]@XJ[...,None])[...,0] + Ai[:-1,-1]

        # now diffeo, not semilagrange here

        for t in range(nt-1,-1,-1):
            Xs = Xs + interp3D(xv,-v[t].permute(3,0,1,2),Xs.permute(3,0,1,2)).permute(1,2,3,0)/nt
        # # and points (not in 3D)        
        # pointsIt = torch.clone(pointsI)
        # if pointsIt.shape[0] >0:
        #     for t in range(nt):            
        #         pointsIt += interp(xv,v[t].permute(2,0,1),pointsIt.T[...,None])[...,0].T/nt
        #     pointsIt = (A[:2,:2]@pointsIt.T + A[:2,-1][...,None]).T
        
        # transform image
        AI = interp3D(xI,I,Xs.permute(3,0,1,2),padding_mode="border")
        

        # transform the contrast
        B = torch.ones(1+AI.shape[0],AI.shape[1]*AI.shape[2]*AI.shape[3],device=AI.device,dtype=AI.dtype)
        B[1:AI.shape[0]+1] = AI.reshape(AI.shape[0],-1)
        #B = torch.ones(10,AI.shape[1]*AI.shape[2],device=AI.device,dtype=AI.dtype)
        #B[1:4] = AI.reshape(AI.shape[0],-1)
        #B[4] = (AI[0][None]**2).reshape(1,-1)
        #B[5] = (AI[1][None]**2).reshape(1,-1)
        #B[6] = (AI[2][None]**2).reshape(1,-1)
        #B[7] = (AI[0][None]*AI[1][None]).reshape(1,-1)
        #B[8] = (AI[0][None]*AI[2][None]).reshape(1,-1)
        #B[9] = (AI[1][None]*AI[2][None]).reshape(1,-1)
        with torch.no_grad():    
            BB = B@(B*WM.ravel()).T
            BJ = B@((J*WM).reshape(J.shape[0],J.shape[1]*J.shape[2]*J.shape[3])).T
            small = 0.1
            coeffs = torch.linalg.solve(BB + small*torch.eye(BB.shape[0],device=BB.device,dtype=BB.dtype),BJ)
        fAI = ((B.T@coeffs).T).reshape(J.shape)

        # objective function
        EM = torch.sum((fAI - J)**2*WM)/2.0/sigmaM**2
        ER = torch.sum(torch.sum(torch.abs(torch.fft.fftn(v,dim=(1,2)))**2,dim=(0,-1))*LL)*DV/2.0/v.shape[1]/v.shape[2]/sigmaR**2
        E = EM + ER
        tosave = [E.item(), EM.item(), ER.item()]
        #if pointsIt.shape[0]>0:
        #    EP = torch.sum((pointsIt - pointsJ)**2)/2.0/sigmaP**2
        #    E += EP
        #    tosave.append(EP.item())
        
        Esave.append( tosave )
        # gradient update
        E.backward()
        with torch.no_grad():            
            L -= (epL/(1.0 + (it>=diffeo_start)*9))*L.grad
            T -= (epT/(1.0 + (it>=diffeo_start)*9))*T.grad

            L.grad.zero_()
            T.grad.zero_()
            

            # v grad
            vgrad = v.grad
            # smooth it            
            if it >= diffeo_start:
                vgrad = torch.fft.ifftn(torch.fft.fftn(vgrad,dim=(1,2,3))*K[...,None],dim=(1,2,3)).real
                v -= vgrad*epV
            v.grad.zero_()


        # update weights
        if not it%5:
            with torch.no_grad():
                # M step for these params
                if estimate_muA:
                    muA = torch.sum(WA*J,dim=(-1,-2,-3))/torch.sum(WA)
                if estimate_muB:
                    muB = torch.sum(WB*J,dim=(-1,-2,-3))/torch.sum(WB)
                #if it <= 200:
                #    muA = torch.tensor([0.75,0.77,0.79],device=J.device,dtype=J.dtype)
                #    muB = torch.ones(J.shape[0],device=J.device,dtype=J.dtype)*0.96

                if it >= 50:

                    W = torch.stack((WM,WA,WB))
                    pi = torch.sum(W,dim=(1,2,3))
                    pi += torch.max(pi)*1e-6
                    pi /= torch.sum(pi)


                    # now the E step, update the weights
                    WM = pi[0]* torch.exp( -torch.sum((fAI - J)**2,0)/2.0/sigmaM**2 )/np.sqrt(2.0*np.pi*sigmaM**2)**J.shape[0]
                    WA = pi[1]* torch.exp( -torch.sum((muA[...,None,None,None] - J)**2,0)/2.0/sigmaA**2 )/np.sqrt(2.0*np.pi*sigmaA**2)**J.shape[0]
                    WB = pi[2]* torch.exp( -torch.sum((muB[...,None,None,None] - J)**2,0)/2.0/sigmaB**2 )/np.sqrt(2.0*np.pi*sigmaB**2)**J.shape[0]
                    WS = WM+WB+WA
                    WS += torch.max(WS)*1e-6
                    WM /= WS
                    WB /= WS
                    WA /= WS


        # draw
        if not it%10:
            ax[0].cla()
            Ishow = ((AI-torch.amin(AI,(1,2,3))[...,None,None])/(torch.amax(AI,(1,2,3))-torch.amin(AI,(1,2,3)))[...,None,None,None]).permute(1,2,3,0).clone().detach().cpu()
            ax[0].imshow(  Ishow[0,...,0] ,extent=extentJ)
            #ax[0].scatter(pointsIt[:,1].clone().detach().cpu(),pointsIt[:,0].clone().detach().cpu())
            ax[0].set_title('space tformed atlas')

            ax[1].cla()    
            Ishow = clip(fAI.permute(1,2,3,0).clone().detach()/torch.max(J).item()).cpu()
            ax[1].imshow(Ishow[0,...,0],extent=extentJ,vmin=0,vmax=1)
            #ax[1].scatter(pointsIt[:,1].clone().detach().cpu(),pointsIt[:,0].clone().detach().cpu())
            ax[1].set_title('contrast tformed atlas')
            
            ax[5].cla()
            Ishow = clip( (fAI - J)/(torch.max(J).item())*3.0  ).permute(1,2,3,0).clone().detach().cpu()*0.5+0.5
            ax[5].imshow(Ishow[0,...,0],extent=extentJ)
            #ax[5].scatter(pointsIt[:,1].clone().detach().cpu(),pointsIt[:,0].clone().detach().cpu())
            #ax[5].scatter(pointsJ[:,1].clone().detach().cpu(),pointsJ[:,0].clone().detach().cpu())
            ax[5].set_title('Error')

            ax[2].cla()
            Ishow = J.permute(1,2,3,0).cpu()/torch.max(J).item()
            ax[2].imshow(Ishow[0,...,0],extent=extentJ,vmin=0,vmax=1)
            #ax[2].scatter(pointsJ[:,1].clone().detach().cpu(),pointsJ[:,0].clone().detach().cpu())
            ax[2].set_title('Target')

            ax[4].cla()
            ax[4].imshow(clip(torch.stack((WM,WA,WB),-1).clone().detach()).cpu()[0],extent=extentJ)
            ax[4].set_title('Weights')


            toshow = v[0].clone().detach().cpu() # initial velocity, components are rgb
            toshow /= torch.max(torch.abs(toshow))
            toshow = toshow*0.5+0.5
            #toshow = torch.cat((toshow,torch.zeros_like(toshow[...,0][...,None])),-1)   
            ax[3].cla()
            ax[3].imshow(clip(toshow)[toshow.shape[0]//2],extent=extentV)
            ax[3].set_title('velocity')
            
            axE[0].cla()
            axE[0].plot(Esave)
            axE[0].legend(['E','EM','ER','EP'])
            axE[0].set_yscale('log')
            axE[1].cla()
            axE[1].plot([e[:2] for e in Esave])
            axE[1].legend(['E','EM'])
            axE[1].set_yscale('log')
            axE[2].cla()
            axE[2].plot([e[2] for e in Esave])
            axE[2].legend(['ER'])
            axE[2].set_yscale('log')


            fig.canvas.draw()
            figE.canvas.draw()
            
    return {
        'A': A.clone().detach(), 
        'v': v.clone().detach(), 
        'xv': xv, 
        'WM': WM.clone().detach(),
        'WB': WB.clone().detach(),
        'WA': WA.clone().detach(),
        'Xs': Xs.clone().detach()
    }


def build_transform(xv,v,A,direction='b',XJ=None):
    ''' Create sample points to transform atlas to target from affine and velocity.
    
    Parameters
    ----------
    xv : list of array
        Sample points for velocity
    v : array
        time dependent velocity field
    A : array
        Affine transformation matrix
    direction : char
        'f' for forward and 'b' for backward. 'b' is default and is used for transforming images.
        'f' is used for transforming points.
    XJ : array
        Sample points for target (meshgrid with ij index style).  Defaults to None 
        to keep sampling on the xv.
    
    Returns
    -------
    Xs : array
        Sample points in mehsgrid format.
    
    
    '''
    
    A = torch.tensor(A)
    if v is not None: v = torch.tensor(v) 
    if XJ is not None:
        # check some types here
        if isinstance(XJ,list):
            if XJ[0].ndim == 1: # need meshgrid
                XJ = torch.stack(torch.meshgrid([torch.tensor(x) for x in XJ],indexing='ij'),-1)
            elif XJ[0].ndim == 2: # assume already meshgrid
                XJ = torch.stack([torch.tensor(x) for x in XJ],-1)
            else:
                raise Exception('Could not understand variable XJ type')
            
        # if it is already in meshgrid form we just need to make sure it is a tensor
        XJ = torch.tensor(XJ)
    else:
        XJ = torch.stack(torch.meshgrid([torch.tensor(x) for x in xv],indexing='ij'),-1)
        
    if direction == 'b':
        Ai = torch.linalg.inv(A)
        # transform sample points
        Xs = (Ai[:-1,:-1]@XJ[...,None])[...,0] + Ai[:-1,-1]    
        # now diffeo, not semilagrange here
        if v is not None:
            nt = v.shape[0]
            for t in range(nt-1,-1,-1):
                Xs = Xs + interp(xv,-v[t].permute(2,0,1),Xs.permute(2,0,1)).permute(1,2,0)/nt
    elif direction == 'f':
        Xs = torch.clone(XJ)
        if v is not None:
            nt = v.shape[0]
            for t in range(nt):
                Xs = Xs + interp(xv,v[t].permute(2,0,1),Xs.permute(2,0,1)).permute(1,2,0)/nt
        Xs = (A[:2,:2]@Xs[...,None])[...,0] + A[:2,-1]    
            
    else:
        raise Exception(f'Direction must be "f" or "b" but you input {direction}')
    return Xs 

def build_transform3D(xv,v,A,direction='b',XJ=None):
    ''' Create sample points to transform atlas to target from affine and velocity.
    
    Parameters
    ----------
    xv : list of array
        Sample points for velocity
    v : array
        time dependent velocity field
    A : array
        Affine transformation matrix
    direction : char
        'f' for forward and 'b' for backward. 'b' is default and is used for transforming images.
        'f' is used for transforming points.
    XJ : array
        Sample points for target (meshgrid with ij index style).  Defaults to None 
        to keep sampling on the xv.
    
    Returns
    -------
    Xs : array
        Sample points in mehsgrid format.
    
 
    
    '''
    
    A = torch.tensor(A)
    v = torch.tensor(v)
    if XJ is not None:
        # check some types here
        if isinstance(XJ,list):
            if XJ[0].ndim == 1: # need meshgrid
                XJ = torch.stack(torch.meshgrid([torch.tensor(x) for x in XJ],indexing='ij'),-1)
            elif XJ[0].ndim == 3: # assume already meshgrid
                XJ = torch.stack([torch.tensor(x) for x in XJ],-1)
            else:
                raise Exception('Could not understand variable XJ type')
            
        # if it is already in meshgrid form we just need to make sure it is a tensor
        XJ = torch.tensor(XJ)
    else:
        XJ = torch.stack(torch.meshgrid([torch.tensor(x) for x in xv],indexing='ij'),-1)
        
    if direction == 'b':
        Ai = torch.linalg.inv(A)
        # transform sample points
        Xs = (Ai[:-1,:-1]@XJ[...,None])[...,0] + Ai[:-1,-1]    
        # now diffeo, not semilagrange here
        nt = v.shape[0]
        for t in range(nt-1,-1,-1):
            Xs = Xs + interp3D(xv,-v[t].permute(3,0,1,2),Xs.permute(3,0,1,2)).permute(1,2,3,0)/nt
    elif direction == 'f':
        Xs = torch.clone(XJ)
        nt = v.shape[0]
        for t in range(nt):
            Xs = Xs + interp(xv,v[t].permute(3,0,1,2),Xs.permute(3,0,1,2)).permute(1,2,0)/nt
        Xs = (A[:-1,:-1]@Xs[...,None])[...,0] + A[:-1,-1]    
            
    else:
        raise Exception(f'Direction must be "f" or "b" but you input {direction}')
    return Xs 

def transform_image_atlas_with_A(A, XI, I, XJ):
    '''
    Transform an image with an affine matrix
    
    Parameters
    ----------
    
    A  : torch tensor
         Affine transform matrix
        
    XI : list of numpy arrays
         List of arrays storing the pixel location in image I along each image axis. 
         convention is row column order not xy. i.e, 
         locations of pixels along the y axis (rows) followed by
         locations of pixels along the x axis (columns)  
    
    I  : numpy array
         A rasterized image with len(blur) channels along the first axis
        
    XJ : list of numpy arrays
         List of arrays storing the pixel location in image I along each image axis. 
         convention is row column order not xy. i.e, 
         locations of pixels along the y axis (rows) followed by
         locations of pixels along the x axis (columns)         
    
    Returns
    -------
    AI : torch tensor
        image I after affine transformation A, with channels along first axis
              
    '''
    xv = None
    v = None
    AI= transform_image_atlas_to_target(xv, v, A, XI, I, XJ=XJ)
    return AI

def transform_image_atlas_to_target(xv,v,A,xI,I,XJ=None):
    '''
    Transform an image
    '''
    phii = build_transform(xv,v,A,direction='b',XJ=XJ)    
    phiI = interp(xI,I,phii.permute(2,0,1),padding_mode="border")
    return phiI
    
    
def transform_image_target_to_atlas(xv,v,A,xJ,J,XI=None):
    '''
    Transform an image
    '''
    phi = build_transform(xv,v,A,direction='f',XJ=XI)    
    phiiJ = interp(xJ,J,phi.permute(2,0,1),padding_mode="border")
    return phiiJ
    
def transform_points_atlas_to_target(xv,v,A,pointsI):
    '''
    Transform points.  Note points are in row column order, not xy.
    '''
    #phi = build_transform(xv,v,A,direction='f',XJ=XI)
    if isinstance(pointsI,torch.Tensor):
        pointsIt = torch.clone(pointsI)
    else:
        pointsIt = torch.tensor(pointsI)
    nt = v.shape[0]
    for t in range(nt):            
        pointsIt += interp(xv,v[t].permute(2,0,1),pointsIt.T[...,None])[...,0].T/nt
    pointsIt = (A[:2,:2]@pointsIt.T + A[:2,-1][...,None]).T
    return pointsIt
def transform_points_target_to_atlas(xv,v,A,pointsI):
    '''
    Transform points.  Note points are in row column order, not xy.
    '''
    
    if isinstance(pointsI,torch.Tensor):
        pointsIt = torch.clone(pointsI)
    else:
        pointsIt = torch.tensor(pointsI)
    Ai = torch.linalg.inv(A)
    pointsIt = (Ai[:2,:2]@pointsIt.T + Ai[:2,-1][...,None]).T
    nt = v.shape[0]
    for t in range(nt):            
        pointsIt += interp(xv,-v[t].permute(2,0,1),pointsIt.T[...,None])[...,0].T/nt
    return pointsIt

#given two sets of points, caluclates the target registratoin error (TRE) and their mean/std
def calculate_tre(pointsI, pointsJ):
    TRE_i = np.sqrt(np.sum((pointsI - pointsJ)**2,axis=1))
    meanTRE = np.mean(TRE_i)
    stdTRE = np.std(TRE_i)
    return meanTRE, stdTRE


#Download and store ontology
def download_aba_ontology(url,file_name): 
	''' Create 3D altas ontology
	    
	    Parameters
	    ----------
	    url : Link to url contain atlas ontology
	    file_name : File name to save atlas ontology
	    
	    Returns
	    -------
	    ontology_name : File name to store ontologies
	    namesdict : Dictionary of all brain rgeion names
	'''
	    
   
	r = requests.get(url)
	print(r)
	with open(file_name,'w') as f:
        	f.write(r.text)
	ontology_name = file_name

	O = pd.read_csv(ontology_name)

	# store the ontology in a dictionary
	namesdict = defaultdict(lambda: 'unk')
	namesdict[0] = 'bg'

	# we need to add the structure names from the structure_id
	for i,n in zip(O['id'],O['acronym']):
        	namesdict[i] = n
	return ontology_name,namesdict

def download_aba_image_labels(imageurl, labelurl, imagefile, labelfile):
	''' Create 3D altas image and region annotations
		    
		    Parameters
		    ----------
		    imageurl : array
		    		Link containing atlas cell stained image.
		    labelurl : array
		    		Link containing atlas altas images for each voxel in imageurl.
		    imagefile : array
		    		File location to save imageurl information.
		    labelfile : array
		    		File location to save labelurl information.
		    
		    Returns
		    -------
		    imagefile : array
		    		File location to save imageurl information.
		    labelfile : array
		    		File location to save labelurl information.	    
		    
	'''
	url = imageurl
	r = requests.get(url, stream=True)
	with open(imagefile, 'wb') as f:
            for chunk in r.iter_content(chunk_size=1024): 
                if chunk: # filter out keep-alive new chunks
                    f.write(chunk)
	imagefile = imagefile

	url = labelurl#'http://download.alleninstitute.org/informatics-archive/current-release/mouse_ccf/annotation/ccf_2017/annotation_50.nrrd'
	r = requests.get(url, stream=True)
	with open(labelfile, 'wb') as f:
            for chunk in r.iter_content(chunk_size=1024): 
                if chunk: # filter out keep-alive new chunks
                    f.write(chunk)
	labelfile = labelfile
	return imagefile, labelfile

def analyze3Dalign(labelfile, xv,v,mat, xJ, dx, scale_x, scale_y, x, y, X_, Y_, namesdict, device='cpu'):
	''' Create dataframe with region annotations and 3D coordinates of target brain slice
		    
		    Parameters
		    ----------
		    labelfile : array
		    		File name of 3D atlas with region annotations.
		    xv : list of array
        		Sample points for velocity
    		    v : array
        		time dependent velocity field
    		    A : array
        		Affine transformation matrix
    		    direction : char
        		'f' for forward and 'b' for backward. 'b' is default and is used for transforming images.
        		'f' is used for transforming points.
    		    XJ : array
        		Sample points for target (meshgrid with ij index style).  Defaults to None 
       		to keep sampling on the xv.
    
		    dx : int
		    	Step of rasterized image.
		    scale_x : double
		    	Value that x positions of atlas were scaled by
		    scale_y : double
		    	Value that y positions of atlas were scaled by
		    x : array
		    	X positions of target brain slice
		    y : array
		    	Y positions of target brain slice
		    X_ : array
		    	-
		    Y_ : array
		    	-
		    namesdict : Dictionary
		    		Dictionary of brain regions corresponding to region numbers
		    
		    Returns
		    -------
		    df : Dataframe containing each cell in original target brain slice, region annotations and 3D coordinates in terms of the altas.
	'''
	# map the annotations
	#xS = [np.arange(n)*d - (n-1)*d/2.0 for n,d in zip(nxS,dxS)]
	vol,hdr = nrrd.read(labelfile)
	L = vol
	dxL = np.diag(hdr['space directions'])
	nL = L.shape
	xL = [np.arange(n)*d - (n-1)*d/2 for n,d in zip(nL,dxL)]

	# next we'll chose a set of points to sample on
	# if we wanted to use the same resolution as our rasterized image, we would do this
	res = np.array(dx)
	XJ = np.stack(np.meshgrid(np.zeros(1),xJ[0],xJ[1],indexing='ij'),-1)
	# if we want to use a different resolution
	res = 10.0
	XJ = np.stack(np.meshgrid(np.zeros(1), np.arange(xJ[0][0],xJ[0][-1],res), np.arange(xJ[1][0],xJ[1][-1],res), indexing='ij'),-1)


	tform = build_transform3D(xv,v,mat,direction='b',XJ=torch.tensor(XJ,device=mat.device))

	AphiL = interp3D(
        	xL, 
        	torch.tensor(L[None].astype(np.float64),dtype=torch.float64,device=tform.device),
        	tform.permute(-1,0,1,2),
        	mode='nearest',
	)[0,0].int()
	AphiL = AphiL.detach().cpu()

	# now look at the cells and assign
	q = np.stack((y,x))
	qi = np.round(((q - np.stack([xJ[0][0],xJ[1][0]])[...,None])/res)).astype(int)
	# by definition, no points should be out of bounds
	# but if I change resolutions, there may be points out of bounds
	df_ = pd.DataFrame()
	labels = AphiL[qi[0],qi[1]]
	col = ((x - X_[0])/dx).astype(int)
	row = ((y - Y_[0])/dx).astype(int)
	df_['coord0'] = tform[0,row,col,0].detach().cpu()
	df_['coord1'] = tform[0,row,col,1].detach().cpu()
	df_['coord2'] = tform[0,row,col,2].detach().cpu()
	df_['x'] = x
	df_['y'] = y
	df_['struct_id'] = labels
    
	all_names = [namesdict[i] for i in df_['struct_id']]
	df_['acronym'] = all_names  
    
	return df_ 

def plot_brain_regions(df):
	''' Plot all brain regions in target brain slice with different color.
		    
		    Parameters
		    ----------
		    df : DataFrame
		    	Dataframe containing each cell in original target brain slice, region annotations and 3D coordinates in terms of the altas.
		    
		    Returns
		    -------
		    None
		    
	'''
	#plot brain regions
	brain_regions = np.unique(df['acronym'])
	fig,ax = plt.subplots()
	for i in range(len(brain_regions)):
            region_df = df[df['acronym']==brain_regions[i]]

            ax.scatter(region_df['x'], region_df['y'], label = brain_regions[i],s= 0.1)   
            ax.legend()

def plot_subset_brain_regions(df, brain_regions):
	''' Plot subset of brain regions in target brain slice with different color.
		    
		    Parameters
		    ----------
		    df : DataFrame
		    	Dataframe containing each cell in original target brain slice, region annotations and 3D coordinates in terms of the altas.
		    brain_regions : array
		    		Subset of brain regions of interest (i.e. ['CA1', 'CP', 'DG-sg'])
		    
		    Returns
		    -------
		    None
		    
	'''
    	#plot brain regions
	brain_regions = brain_regions
	fig,ax = plt.subplots()
	ax.scatter(df['x'], df['y'],color = 'grey',s= 0.1)   
	for i in range(len(brain_regions)):
	    region_df = df[df['acronym']==brain_regions[i]]
	    ax.scatter(region_df['x'], region_df['y'], label = brain_regions[i],s= 0.1)   
	    ax.legend()
```
